# Honoring the Afro-Colombian musical culture with the naming of *Epipedobates currulao* sp. nov. (Anura, Dendrobatidae), a frog from the Pacific rainforests

**DOI:** 10.1101/2024.03.23.586415

**Authors:** Mileidy Betancourth-Cundar, Juan Camilo Ríos-Orjuela, Andrew J. Crawford, David C. Cannatella, Rebecca D. Tarvin

**Affiliations:** Departamento de Ciencias Biológicas, Universidad de los Andes, Bogotá, 111711, Colombia; Department of Biology, Stanford University, Palo Alto, CA, 94305, USA; Grupo de Morfología y Ecología Evolutiva, Instituto de Ciencias Naturales, Universidad Nacional de Colombia, Sede Bogotá, 7495, Colombia; Museo de Historia Natural C.J. Marinkelle, Universidad de los Andes, Bogotá, 111711, Colombia; Department of Integrative Biology and Biodiversity Center, University of Texas, Austin, TX 78712, USA; Museum of Vertebrate Zoology and Department of Integrative Biology, University of California, Berkeley, Berkeley, CA 94720, USA

**Keywords:** bioacoustics, Chocó, music, megadiverse, poison frogs, new species, alpha taxonomy, DNA barcoding

## Abstract

The number of amphibian species described yearly shows no signs of slowing down, especially in tropical regions, implying that the biodiversity of amphibians remains woefully underestimated. We describe a new species of poison frog from the Pacific lowlands of southwestern Colombia: *Epipedobates currulao* sp. nov., named for the Pacific music and dance genre known as *bambuco viejo* or *currulao*. This species inhabits lowland forests from 0–260 m a.s.l. This taxon differs from congeners by having a combination of bright yellow blotches in the dorsal anterior region of the thigh and upper arm, homogenous dark-brown dorsal coloration, and advertisement calls of long duration and many pulses. We also describe the courtship call of *E. currulao*, which is lower in frequency and shorter in duration than its advertisement call. Molecular phylogenetic analyses confirm the monophyly of the populations sampled and its position as the sister species of *Epipedobates narinensis*, which occurs in southwestern Colombia. Among species of *Epipedobates*, the new species has been previously confused with *E. boulengeri*, but we find that the two species are allopatric and represent two divergent clades (1.77% divergent for *12S–16S* and 5.39% for *CYTB*). These species can be distinguished by the presence of a bright yellow blotch on the dorsal anterior region of the thigh and on the upper arm of *E. currulao*, blotches that are either more white than yellow or absent in *E. boulengeri*. In addition, the advertisement calls are distinct, with *E. currulao* having a single but long call in each call series while *E. boulengeri* has 2–6 calls in a series with each call being much shorter in length. *Epipedobates currulao* is the most northern species of *Epipedobates*, which extends southwards along the western edge of the Andes. Known as the Chocó, this biogeographic region has been largely converted to agriculture in Ecuador and is experiencing widespread transformation in Colombia, which may endanger *E. currulao* and biodiversity in the region. A Spanish translation of the main text is available in the Supplementary Materials.

**Resumen:** El número de especies de anfibios descritas cada año continúa aumentando, especialmente en las regiones tropicales, lo que implica que la biodiversidad de anfibios sigue siendo subestimada. Describimos una nueva especie de rana venenosa de las tierras bajas del Pacífico del suroccidente de Colombia: *Epipedobates currulao* sp. nov., nombrada así por el género de música y danza del Pacífico conocido como *bambuco viejo* o *currulao*. Las ranas de esta especie habitan en bosques de tierras bajas desde el nivel del mar hasta los 260 m. Este taxón se diferencia de sus congéneres por tener una combinación de manchas amarillas brillantes en la región dorsal anterior del muslo y los brazos, una coloración dorsal homogénea marrón oscuro y cantos de advertencia más largos y en consecuencia con mayor número de pulsos. También describimos la llamada de cortejo de *E. currulao,* con menor frecuencia pico y duración que la llamada de advertencia. Los análisis filogenéticos confirman la monofilia de la especie y su posición como hermana de *Epipedobates narinensis*, la cual se distribuye en el suroccidente de Colombia. Entre las especies de Epipedobates, la nueva especie ha sido previamente asignada a *E. boulengeri,* pero las dos especies son alopátricas y representan dos clados filogenéticamente divergentes (1.77% divergentes para *12S–16S* y 5.39% para *CYTB*). Estas especies se pueden distinguir fenotípicamente por la presencia de una mancha amarilla brillante en la región dorsal anterior del muslo y en la parte superior del brazo en *E. currulao*, que son más blancas que amarillas o están ausentes en *E. boulengeri.* Además, los cantos de advertencia son distintos, *E. currulao* tiene una única y larga llamada en una serie de llamadas, mientras que *E. boulengeri* tiene de 2 a 6 llamadas por serie, siendo cada llamada mucho más corta. *Epipedobates currulao* es la especie distribuida más al norte del género *Epipedobates*, el cual se extiende hacia el sur a lo largo del flanco occidental de la cordillera de los Andes. Esta región conocida como el Chocó biogeográfico, ha sido fuertemente transformada por agricultura en Ecuador y está experimentando una transformación generalizada de sus bosques en Colombia, lo cual pone en peligro a *E. currulao* y toda su biodiversidad en un futuro cercano. Una traducción al español del texto principal está disponible en el material suplementario.

## INTRODUCTION

*Epipedobates* is a taxon of neotropical poison frogs (Anura, Dendrobatidae) with nine putative species (Grant et al. 2006, 2017; López-Hervas et al. 2024): *Epipedobates* aff. *espinosai* (see López-Hervas et al. 2024)*, E. anthonyi* (Noble, 1921), *E. boulengeri* (Barbour, 1909), *E.* sp. 1 (see López-Hervas et al. 2024)*, E. espinosai* (Funkhouser, 1956), *E. machalilla* (Coloma, 1995), *E. maculatus* (Peters, 1873)*, E. narinensis (*Mueses-Cisneros et al. 2008), and *E. tricolor* (Boulenger, 1899). Frogs of the genus *Epipedobates* inhabit dry and humid tropical forests from 0 to 1800 m a.s.l. across the Pacific lowlands and foothills of the western side of the Andes of Colombia, Ecuador, and northern Peru (Grant et al. 2006, 2017). A recent assessment of genetic and phenotypic diversity in *Epipedobates boulengeri* found several distinct evolutionary lineages with similar phenotypes, i.e., cryptic species (López-Hervas et al. 2024). It is thus no surprise that prior phylogenetic studies, which sampled different subsets of lineages that we now know correspond to different species, failed to resolve some phylogenetic relationships in *Epipedobates* (Clough & Summers 2000; Grant et al. 2017; Santos et al. 2014; Tarvin et al. 2017; Vences et al. 2003).

Previous studies of *Epipedobates boulengeri* demonstrated high genetic diversity and interpopulation variation in acoustic traits and larval morphology, indicating that the lineages assigned to *Epipedobates boulengeri* most likely represented a species complex (López-Hervas et al. 2024; Lötters et al. 2003; Santos et al. 2009; Tarvin et al. 2017). The most recent phylogenetic assessment (López-Hervas et al. 2024) concluded that *Epipedobates boulengeri* contained representatives of four genetically distinct lineages: *E. boulengeri* (*sensu stricto*), distributed in southwestern Colombia and the northwestern edge of Ecuador; *E. espinosai,* distributed in central-northwestern Ecuador; *E.* sp. 1, distributed in southwestern Colombia, and *E.* aff. *espinosai*, distributed in northwestern Ecuador and up into the southern end of the Andean foothills of Colombia (López-Hervas et al. 2024). Phenotypic evidence supports this cryptic diversity, for example Lötters et al. (2003) found differences in the number of notes and the length of the advertisement call between specimens referred to as *E. boulengeri* from Colombia (Anchicayá, Valle del Cauca, corresponding to *E.* sp. 1) and *E. boulengeri* from Ecuador (Lita, Imbabura, corresponding to *E.* aff. *espinosai*). In addition, Anganoy-Criollo and Cepeda-Quilindo (2017) described the external morphology of the tadpoles of *E. boulengeri* from Colombia (Tumaco, Nariño, possibly representing *E. boulengeri sensu stricto*) and from Ecuador (Carchi, Esmeraldas, Imbabura, and Santo Domingo provinces, identification uncertain, possibly corresponding to *E.* aff. *espinosai*, *E. boulengeri sensu stricto*, or *E. espinosai*). Although they only examined tadpoles from the southern half of the species distribution area of *E. boulengeri* (i.e., excluding the lineage described as *E.* sp. 1 by López-Hervas et al. 2024), they found that the Ecuadorian populations differed from those of Colombia in body shape, tips of marginal papillae, and pattern of tail coloration. Thus, previous evidence pointed strongly to *E. boulengeri* as a species complex.

In this study, we describe a new species of *Epipedobates* from the Colombian Pacific lowlands corresponding to *E.* sp. 1 as assigned by López-Hervas et al. (2024). Our acoustic, morphological, and molecular analyses are consistent with work by López-Hervas et al. (2024), which identified this frog as a distinct and previously unrecognized species. We name the new species *Epipedobates currulao* sp. nov. and provide a detailed description of its calls, morphology, behavior, and natural history. As with many other amphibian species (Myers et al. 2000; Powers and Jetz 2019; Warren et al. 2013), *Epipedobates currulao* sp. nov. is under threat from habitat loss, pollution, and climate change. By identifying and describing this new species, we can better understand the diversity, evolution, geographic distribution, habitat requirements, and conservation needs of *Epipedobates* species in Colombia. Our findings will stimulate further research and conservation efforts to protect the unique and irreplaceable amphibian biodiversity of the Chocó biogeographical region.

## METHODS

### Ethics statement

Scientific collecting procedures involving live animals followed protocols approved by the IACUC at the University of Texas at Austin (AUC-2012-00032), the University of California, Berkeley (AUP-2019-08-12457), and *CICUAL* of the Universidad de los Andes (POE 18–003). Research and field collection of samples were authorized by the Autoridad Nacional de Licencias Ambientales (ANLA) of Colombia under the *permiso marco resolución* No. 1177 to the Universidad de los Andes. Specimens of *E. boulengeri* from the type locality were collected under Res. 061-2016 by Parques Nacionales Naturales de Colombia. Samples were exported under the following permits: CITES No. CO39282, CO41443, CO46948, and ANLA No. 00561.

### Scientific collecting

We collected genetic, morphological, and acoustic data, as well as whole specimens, of different species and populations of *Epipedobates* in Colombia (Fig. 1; see also López-Hervas et al. 2024) during 2014, 2016, and 2022. Animals were euthanized by an overdose of lidocaine. Sex for most of the specimens was determined at the time of collection based on whether individuals were calling or not and was confirmed by direct examination of the gonads after euthanasia. Before fixation, liver and muscle samples were taken for molecular genetic analyses and stored in 95% ethanol. In some cases, skin was removed for quantification of defensive alkaloid content (see Suppl. material 1). Voucher specimens were fixed in 10% neutral buffered formalin and transferred directly to 70% ethanol for long-term storage in the amphibian collection of the Museo de Historia Natural C.J. Marinkelle at the Universidad de los Andes in Bogotá, Colombia (ANDES:A) and the Herpetology Collection at the Museum of Vertebrate Zoology at the University of California, Berkeley in Berkeley, California, USA (MVZ:Herp).

**Figure 1.**
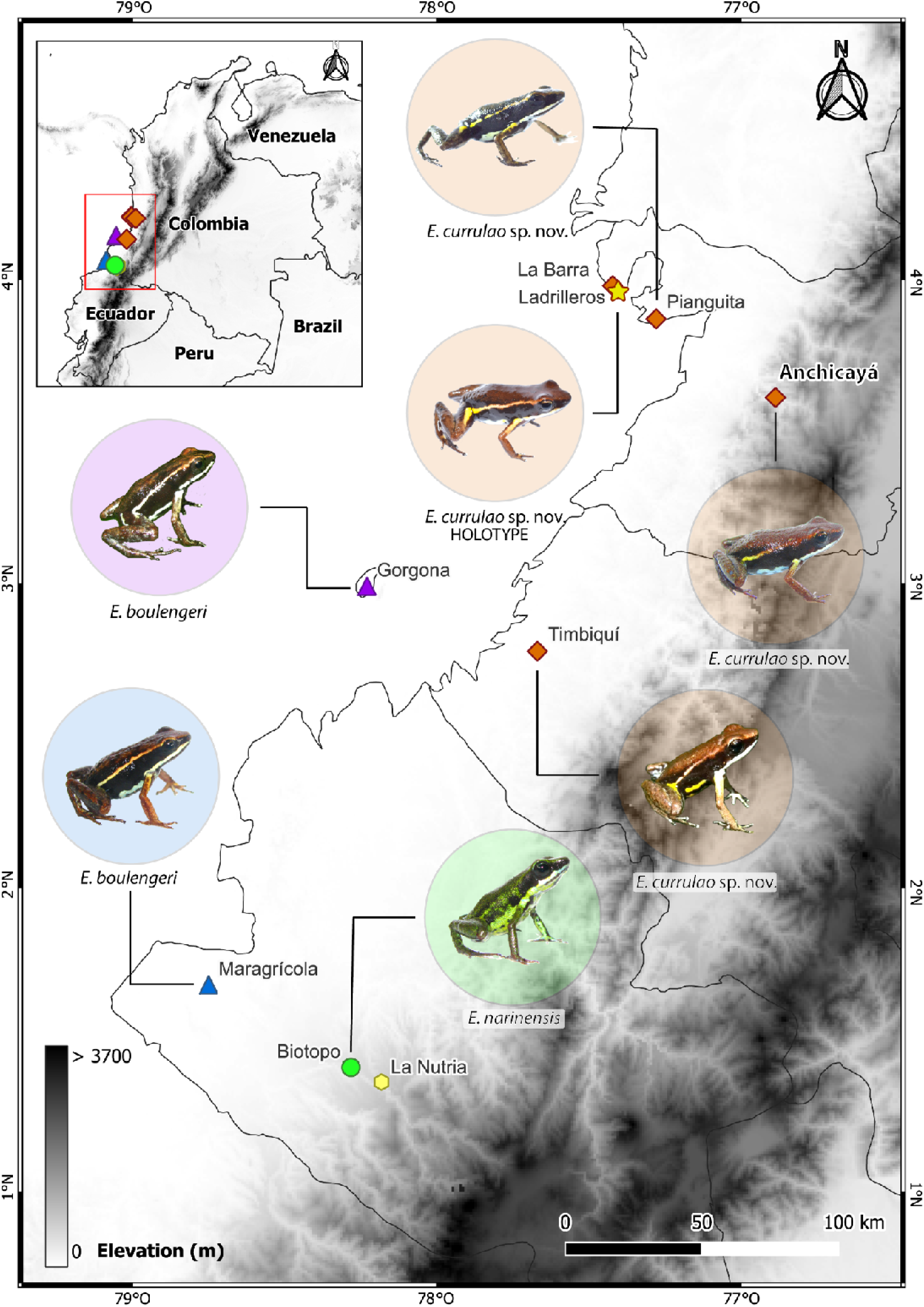
Map of research area. The southern half of the Pacific lowlands of Colombia, part of the Chocó biogeographical region, hosts at least three species of *Epipedobates*. Photograph of the frog from Timbiquí (Fundación ProAves y la Reserva Ranita Terribilis) was obtained from iNaturalist (observation No. 135253843); other images were taken by the authors.

### Morphology

Following previous descriptions of new *Epipedobates* species (Cisneros-Heredia and Yánez-Muñoz 2010; Mueses-Cisneros et al. 2008), we took 13 standardized morphometric measurements after fixation (Watters et al. 2016): Snout-vent-length (SVL); forearm length (FAL), taken from tip of the flexed elbow and the proximal edge of the palmar tubercle; hand length (HaL), measured from the proximal edge of the large medial palmar tubercle to the tip of the longest (fourth) finger; tibia length (TL), taken from the outer surface of the flexed knee to the heel inflection; foot length (FL), taken from the proximal edge of the external metatarsal tubercle to the end of the fourth toe; head width (HW), taken between the angles of the jaws; head length (HL), distance from the tip of the snout to the angle of the jaw; distance from the center of one nares to the anterior edge of the eye (NED); eye width (EW); eye-nares distance (END); internarial distance (IND); interocular distance (IOD); and horizontal diameter of the tympanum (TD). All measurements were taken under a dissecting microscope using a digital caliper at a resolution of 0.01 mm. We conducted a Kruskal-Wallis test to compare body size (snout-vent-length) across species and a post-hoc paired comparison using a Wilcoxon rank-sum test. We also conducted an ANOVA to assess sexual dimorphism in body size, previously checked its assumptions.

We also reviewed and summarized species-level phenotypic variation as described in López-Hervas et al. (2024) to provide a species-level assessment of color pattern variation that can be used to visually distinguish species.

### Molecular and phylogenetic analyses

DNA was extracted from liver and muscle tissue using the Qiagen DNeasy Blood & Tissue kit (Valencia, CA) following the manufacturer’s protocol. We sequenced two mitochondrial gene fragments from 5 individuals of *Epipedobates currulao* sp. nov. collected from the type locality: a fragment including parts of the *12S* and *16S* mitochondrial rRNA genes and the intervening valine tRNA gene (*12S–16S; 692 bp*), and cytochrome b (*CYTB*; 659 bp). We used the same primers and protocols as described in López-Hervas et al. (2024). The 12S–16S region was sequenced using forward primer 12Sa (5’-AACTGGGATTAGATACCCCACTAT-3’) and reverse primer 16SH-H (5’-TACCTTTTGCATCATGGTCTAGC-3’) with PCR conditions: 2 min at 94°C for the initial denaturation followed by 35 cycles of 30 s at 94°C, 30 s at 46.5°C, and 1 min at 72°C and a final extension time of 7 min at 72°C (Santos and Cannatella, 2011). The CYTB region was sequenced using forward primer CytbDen3-L (5’-AAYATYTCCRYATGATGRAAYTTYGG-3’) and reverse primer CytbDen1-H (5’-GCRAANAGRAAGTATCATTCNGGYT-3’) using the same PCR conditions as described for *12S–16S*. PCR products were sequenced in both directions at the QB3 Functional Genomics Laboratory at the University of California, Berkeley (RRID:SCR_022170). The chromatograms were trimmed and consensus contigs were obtained using *sangeranalyseR* (Chao et al. 2021) in *R* v.4.3.1 (R Core Team, 2023). New sequences were deposited in GenBank (accession numbers OR789875–OR789884).

We aligned the five new sequences using MUSCLE v3.8.31 (Edgar 2004) in AliView (Larsson 2014) with existing *12S–16S* and *CYTB* data for 108 other *Epipedobates* individuals, including one outgroup (*Silverstoneia nubicola*) and 4 individuals assigned to *Epipedobates* sp. 1 by López-Hervas et al. (2024), which correspond here to *Epipedobates currulao* sp. nov. Using the resulting alignment (Suppl. material 2), we calculated mean pairwise p-distances (uncorrected) between species for each gene with the R script provided by López-Hervas et al. (2024) with the ‘dist.dna’ function in *ape* v5.7.1 (Paradis and Schliep 2019), with model set as “raw” and pairwise.deletion set to TRUE. Following López-Hervas et al. (2024), we excluded the individuals from the *E.* aff. *espinosai* Bilsa and La Tortuga localities, which show evidence of introgression with *E. machalilla*.

We complement our molecular dataset with three nuclear markers (total length of 2227 bp), namely histone H3 (*H3*), bone morphogenetic protein 2 (*BMP2*), and voltage-gated potassium channel 1.3 (*K_V_1.3*), as well as the mitochondrial control region (*CR*; 1031 bp) from previously published sequences of the same 108 individuals referenced above (Suppl. material 1). We also added 28 sequences of another 23 individuals of *Epipedobates* that were previously published (Grant et al. 2006, 2017; Santos and Cannatella 2011; Santos et al. 2003, 2009) to the alignment (see Suppl. material 1 for GenBank numbers). These sequences were first trimmed to the gene regions present in the alignment and then aligned using MUSCLE in AliView. A maximum likelihood phylogeny was estimated using this expanded alignment in IQ-TREE v2.3.3 (Kalyaanamoorthy et al. 2017; Minh et al. 2020) and parameters described in López-Hervas et al. (2024). Briefly, we partitioned protein-coding genes by gene and by codon position. We excluded the first and second codon positions of *BMP* and *H3*, which were invariant across all samples. We selected the model of evolution and data partitions under the Bayesian Information Criterion using option -m TESTMERGE, base frequency parameters including equal and estimated (-mfreq F, FO), and rate heterogeneity parameters including equal, gamma, and invariant (-mrate E, G, I). We ran the analysis three times and checked for consistency across runs. To assess branch support, we performed 10,000 ultrafast bootstrap replicates (Minh et al. 2013) and plotted them onto the optimal likelihood tree (Suppl. material 2).

### Call recording and bioacoustic analyses

We recorded the advertisement calls of twelve male frogs of *Epipedobates currulao* sp. nov. from three Valle de Cauca localities under natural conditions: Anchicayá (four individuals), Pianguita (six), and from the type locality, Ladrilleros (two). We also recorded five males of *E. boulengeri* from the type locality (Isla Gorgona, Cauca, Colombia), five males of *E. boulengeri* from Maragrícola (Tumaco, Nariño, Colombia) and six males of *E. narinensis* from its type locality (Biotopo Natural Reserve, Nariño, Colombia). All recordings were deposited in Fonozoo (FZ-SOUND-CODE 14657-14685). Recordings were taken during the morning (7–13 h) using a unidirectional microphone (Sennheiser K6/ME66) connected to a digital recorder (Marantz PMD660/Zoom H4n Pro) and positioned 50–150 cm in front of a calling male. Immediately after recording, we measured the temperature of the substrate using an infrared thermometer (Oakton model 35629). We digitized the recordings at a minimum of 16-bit resolution and 44.1 kHz sampling rate in RAVEN PRO 1.6 (Bioacoustics Research Program 2014). For the terminology and procedures for measuring call traits we follow Köhler et al. (2017). The spectrograms and oscillograms were graphed using Seewave R package (Sueur et al. 2008) with an FFT window using the Blackman algorithm, window length of 256 samples and overlap of 90%.

We defined a call as the sound produced during a single abdominal muscular contraction (Erdtmann and Amézquita 2009; Köhler et al. 2017). In *Epipedobates*, such contractions produce a single “buzz” composed of several pulses (Brown et al. 2011; Erdtmann and Amézquita 2009; Myers and Daly 1976). We analyzed five consecutive calls per individual by measuring their peak frequency (the frequency with the highest amplitude), low frequency (the lower frequency of the call), high frequency (the upper frequency of the call) and 90% bandwidth (-10 dB threshold; containing the frequency range that encompasses 90% of the sound energy in each call). High and low frequencies of the calls were measured at 20 dB (re 20 mPA) below the peak intensity of the peak frequency. At this value the signal energy could still be clearly distinguished from the background noise. We also measured the following temporal variables: call duration, number of pulses per call and inter-call interval. For *E. currulao* sp. nov., we also measured the pulse rate, pulse duration, and inter-pulse interval. Call parameters for the calls of each individual were averaged to statistically compare among individuals. To avoid redundancy among the acoustic variables, we used a Principal Component Analysis (PCA) to reduce the number of parameters implemented in the *‘PCA’* function of the ‘FactoMineR’ R package (Lê et al. 2008). In order to identify whether *E. currulao* sp. nov. could be differentiated based on advertisement calls, we ran a LDA on the first 5 PCs using the *lda’* function of the MASS R package (Venables and Ripley 2002). We utilized a Kruskal-Wallis test to examine the relevant call traits (peak frequency, call duration and number of pulses per call) that differentiate *E. currulao* sp. nov. from other species and a post-hoc paired comparison using a Wilcoxon rank-sum test. We visually represent these traits using the ‘pirateplot’ function “yarrr” R package (Phillips 2017). Pirate plots show all the raw data, their distribution, the mean, and the 95% highest-density intervals (HDI) of the mean of each estimate (Kampstra 2008). We used 999 iterations to calculate the HDI.

### Repositories, Institutional acronyms, and Institutional abbreviations

Throughout our paper we use globally unique identifiers (GUIDs, Globally Unique Identifiers Task Group 2011) to refer to the voucher specimens at QCAZ (QCAZ:A:XXXX, Museo de Zoología de la Pontificia Universidad Católica del Ecuador, Quito, Ecuador), the MVZ (MVZ:Herp:XXXX, herpetological collections of the Museum of Vertebrate Zoology, University of California, Berkeley, USA), and the amphibian collection at the Museo de Historia Natural C.J. Marinkelle at the Universidad de los Andes, Bogotá, Colombia (ANDES:A:XXXX), in an effort to facilitate future machine-readability of this text. Field series abbreviations are as follows: RDT (Rebecca D. Tarvin), AJC (Andrew J. Crawford), MBC (Mileidy Betancourth-Cundar).

## RESULTS

### Taxonomy

***Epipedobates currulao* sp. nov.**

*Epipedobates boulengeri*: Silverstone (1976), Lötters et al. (2003), Vargas-S & Bolaños-L (1999), Castro-Herrera and Vargas-Salinas (2008), Lynch and Suárez-Mayorga (2004), Lötters et al. (2007).

*Epipedobates* sp. 1: Lopez-Hervas et al. (2024).

Proposed English common name: Currulao Nurse Frog

Proposed Spanish common name: Rana nodriza de currulao

http://zoobank.org/ [registration number to be added once paper is accepted]

### Material examined

#### Holotype

**COLOMBIA** • ♀; Ladrilleros, Buenaventura, Valle del Cauca; 3.945221, −77.364993; 28 m. a.s.l.; 6 Aug. 2022; Rebecca D. Tarvin, Mileidy Betancourth-Cundar, Juan Camilo Ríos-Orjuela leg.; ANDES:A:5255.

#### Paratypes

**COLOMBIA** • 4 ♀♀, 6 ♂♂, 1 ND; same data as for holotype; ANDES:A:5254, 5256–5265 • 3 ♀♀, 2 ♂♂; same data as for holotype; Genbank: OR789880–84 and OR789875–79; MVZ:Herp:295432–295436. 1 ND; Ladrilleros, Buenaventura, Valle del Cauca; 3.950882, −77.358293; 53 m a.s.l.; 26 Nov. 2014; Rebecca D. Tarvin and Fray Arriaga leg.; Genbank: OR489012, OR179791, OR734704, OR179836, and OR179880; ANDES:A:2464.

#### Other material

##### Epipedobates currulao

**COLOMBIA** • 2 ♀♀, 5 ND; Type locality, Ladrilleros, Buenaventura, Valle del Cauca; 3.950882, −77.358293; 53 m a.s.l.; 26 Nov. 2014; Rebecca D. Tarvin and Fray Arriaga leg.; GenBank: OR489011, OR179790, OR734703, OR179835, OR179875; ANDES:A:2458–2463, 2465. The skin was removed prior to preservation for evaluation of alkaloid content from these individuals. • *E. currulao* 1 ♀; La Barra, Buenaventura, Valle del Cauca; 3.985064, –77.376723; 15 m a.s.l.; 26 Nov. 2014; Rebecca D. Tarvin and Fray Arriaga leg.; GenBank: OR489010, OR179789, OR734702, OR179834, OR179851; ANDES:A:2455. • *E. currulao* 5 ♂♂; Pianguita, Buenaventura, Valle del Cauca; 3.841954, –77.198718; 17 m.a.s.l.; 12 Sep. 2016; Rebecca D. Tarvin, Mileidy Betancourth-Cundar, Sandra V. Flechas leg.; ANDES:A:3690–94. The skin was removed prior to preservation for evaluation of alkaloid content from these individuals, with the exception of ANDES:A:3691. • *E. currulao 5* ♀♀, 3 ♂♂, Danubio, Dagua, Valle del Cauca; 3.611528, –76.885194; 705 m a.s.l.; 5 Nov. 2016; Mileidy Betancourth-Cundar, Adolfo Amézquita, and Ivan Beltrán leg.; GenBank: OR489013, OR179792, OR734705, OR179837; ANDES:A:3713–20. The skin was removed prior to preservation for evaluation of alkaloid content from these individuals.

##### Epipedobates boulengeri

**COLOMBIA** • 3 ♀♀, 4 ♂♂, 1 ND; Isla Gorgona, Guapi, Cauca; 2.96465, -78.173685; 20 m a.s.l.; 12 Sep. 2016; Rebecca D. Tarvin, Mileidy Betancourth-Cundar and Sandra V. Flechas leg.; GenBank: OR OR488992, OR179771, OR734681, OR179812; ANDES:A:3695-3702. • 1 ND; Isla Gorgona, Guapi, Cauca; 08 Sep. 2005; ANDES:A:0560; Collector unknown. • *E. boulengeri* 3 ♀♀, 4 ♂♂, 1 ND; Isla Gorgona, Guapi, Cauca; 2.96465, –78.173685; 20 m a.s.l.; 12 Sep. 2016; Rebecca D. Tarvin, Mileidy Betancourth-Cundar, Sandra V. Flechas leg.; GenBank: OR OR488992, OR179771, OR734681, OR179812; ANDES:A:3695-3702. • *E. boulengeri* 8 ♀♀; Maragrícola, Tumaco, Nariño; 1.68084, –78.74924; 7 m a.s.l.; 09 Dec. 2014; Rebecca D. Tarvin, Mileidy Betancourth-Cundar, Cristian Flórez leg.; GenBank: OR488990, OR179769, OR734679, OR179810, OR179862, OR488991, OR179770, OR734680, OR179811, OR179863; ANDES:A:2468–75. • *E. boulengeri* 1 ND; La Nutria, El Diviso, Barbacoas, Nariño; 1.36083, –78.18076; 762 m a.s.l.; 10 Dec. 2014; Rebecca D. Tarvin, Mileidy Betancourth-Cundar, Cristian Flórez leg.; GenBank: OR488977, OR179756, OR734663, OR179794, OR179840; ANDES:A:2476.

##### Epipedobates narinensis

**COLOMBIA** • 3 ♀♀, 5 ♂♂, 1 ND; Reserva Natural Biotopo, Berlín, Barbacoas, Nariño; 1.408999, -78.281246; 518 m a.s.l.; 25 Sep. 2016; Rebecca D. Tarvin, Mileidy Betancourth-Cundar and Cristian Flórez leg.; GenBank: OR489008, OR179787, OR734700, OR179832, OR489009, OR179788, OR734701, OR179833; ANDES:A:3703–3711. • *E. narinensis* 10 ♂♂; Reserva Natural Biotopo, Berlín, Barbacoas, Nariño; 1.411263, –78.285099; 600 m a.s.l.; 22 Jul. 2006; Viviana Moreno-Quintero, Jonh Jairo Mueses-Cisneros, Luisa Mercedes Bravo, Carol Narváez, and Bienvenido Cortés leg.; ICN-A:53344 (holotype), ICN-A:53336–53340, 53342–53346 (paratypes).

##### Andinobates minutus

**COLOMBIA** • 4ND; La Barra, Buenaventura, Valle del Cauca; 3.985064, –77.376723; 15 m a.s.l.; 25 Nov. 2014; Rebecca D. Tarvin and Fray Arriaga leg.; ANDES:A 2451–54. • *A. minutus* 1ND; Ladrilleros, Buenaventura, Valle del Cauca; 3.945221, −77.364993; 28 m a.s.l; 6 Aug. 2022; Rebecca D. Tarvin, Mileidy Betancourth-Cundar, Juan Camilo Ríos-Orjuela leg.; ANDES:A:5266.

##### Epipedobates aff. Espinosai

**ECUADOR** • 1 ND; Lita, Carchi; 12 Aug. 1992; M. Bueno leg.; ICN-A:32504.

### Diagnosis

*Epipedobates currulao* is a small dendrobatid frog (SVL mean = 17.99 mm and SD = 0.95 mm, N = 16 frogs; Tables 1, 2) with uniformly brown dorsal coloration, black sides, a white to yellow oblique lateral stripe, a bright yellow blotch on the anteriodorsal side of thigh and on the upper arm, and a pale-blue or turquoise venter with black mottling (Fig. 2, Suppl. material 3). Calls of *E. currulao* sp. nov are long with a call duration of 0.67–3.88 s (mean = 2.21, SD = 0.54 s, N = 15) and 22–122 pulses per call (mean = 73.98, SD = 18.77, N = 15). They occur in call series of only one call (Tables 3, 4).

**Figure 2.**
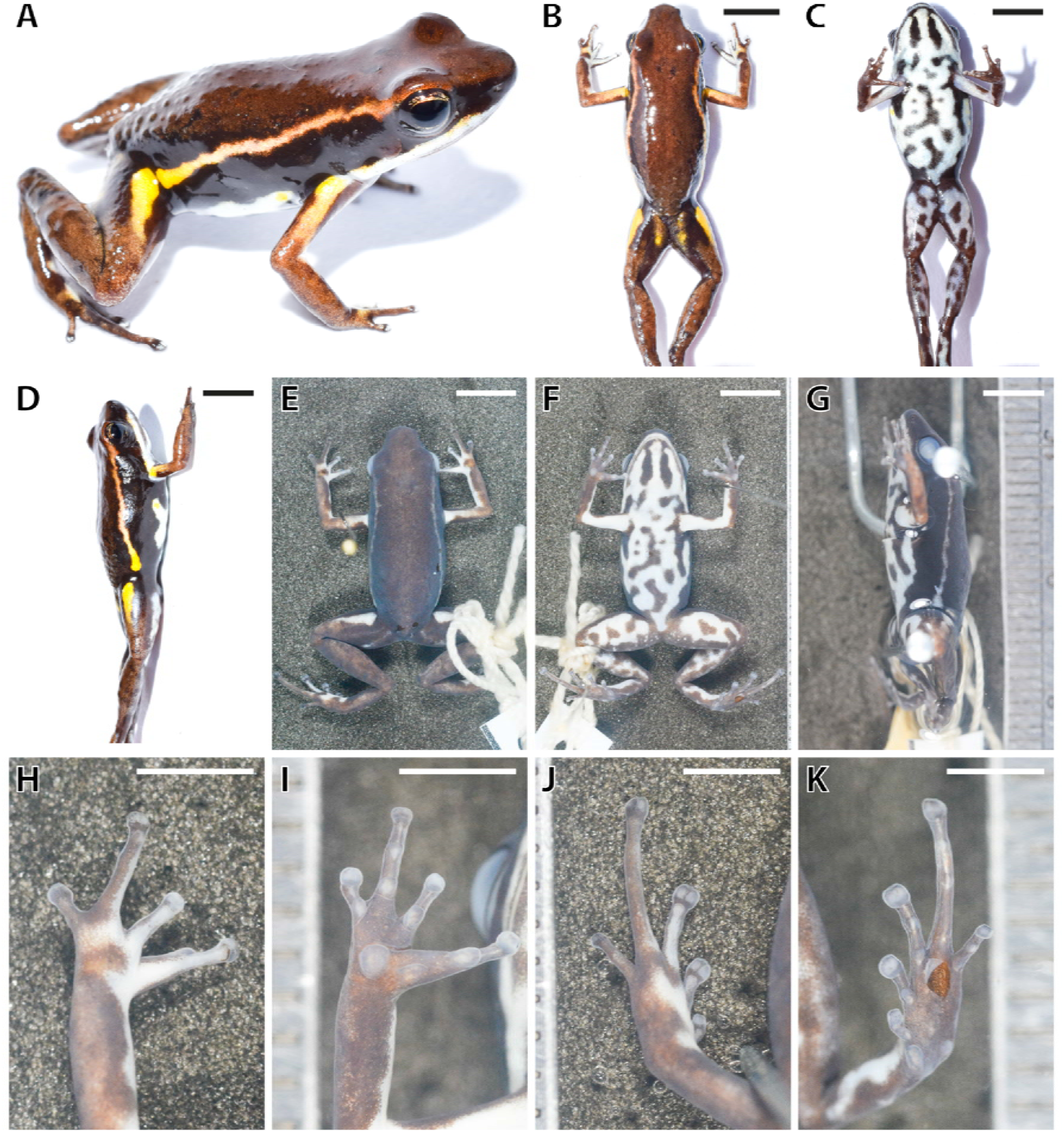
Images in life and in preservative of the holotype of *Epipedobates currulao* sp. nov.. **A** Full specimen in life; **B** Dorsal view in life; **C** Ventral view in life; **D** Lateral view in life; **E** Dorsal view in preservative (70% ethanol); **F** Ventral view in preservative; **G** Lateral view in preservative; **H** Dorsal hand in preservative; **I** Ventral hand in preservative; **J** Dorsal foot in preservative; **K** Ventral foot in preservative. Scale bars represent 5 mm (B to G) or 2.5 mm (H to K).

**Table 1.** Morphometric characters and summary statistics of the males of type series of *Epipedobates currulao* sp. nov. Measurements (in mm) of adult males; values are given as mean, SD, and range. See Methods section for explanation of abbreviations.

| Museum Code | Collector Code | Sex | SVL | FAL | HaL | TL | FL | HW | HL | NED | EW | END | IND | IOD | TD |
| --- | --- | --- | --- | --- | --- | --- | --- | --- | --- | --- | --- | --- | --- | --- | --- |
| ANDES:A:5254 | MBC753 | Male | 17.20 | 4.30 | 3.70 | 7.70 | 7.00 | 5.4 | 6.3 | 2.5 | 2.10 | 1.60 | 2.10 | 2.90 | 0.90 |
| ANDES:A:5256 | AJC07731 | Male | 17.20 | 4.10 | 3.90 | 7.80 | 6.90 | 5.80 | 6.90 | 2.90 | 2.10 | 1.60 | 2.20 | 3.30 | 1.00 |
| ANDES:A:5260 | AJC07735 | Male | 17.40 | 4.40 | 3.70 | 7.90 | 7.00 | 5.60 | 6.40 | 3.00 | 2.50 | 1.60 | 2.50 | 3.80 | 1.00 |
| ANDES:A:5261 | AJC07736 | Male | 18.70 | 4.50 | 4.30 | 8.50 | 7.30 | 5.90 | 7.00 | 3.30 | 2.70 | 1.70 | 2.50 | 3.80 | 1.00 |
| ANDES:A:5263 | AJC07738 | Male | 17.50 | 4.60 | 3.80 | 7.70 | 6.60 | 6.00 | 7.30 | 3.00 | 2.40 | 1.60 | 2.30 | 3.50 | 0.90 |
| MVZ:Herp:29543<br>4 | AJC07742 | Male | 16.70 | 4.00 | 3.60 | 7.50 | 6.70 | 4.90 | 6.70 | 2.80 | 1.90 | 1.70 | 2.30 | 3.20 | 0.90 |
| MVZ:Herp:29543<br>6 | AJC07744 | Male | 16.70 | 3.80 | 3.70 | 7.10 | 6.70 | 5.70 | 6.90 | 2.70 | 2.40 | 1.70 | 2.10 | 3.40 | 1.00 |
| ANDES:A:5259 | RDT0935 | Male | 16.90 | 4.40 | 3.80 | 7.90 | 7.20 | 5.30 | 6.60 | 3.60 | 2.40 | 1.80 | 2.30 | 3.80 | 0.90 |
| Mean |  |  | 17.29 | 4.26 | 3.81 | 7.76 | 6.93 | 5.58 | 6.76 | 2.98 | 2.31 | 1.66 | 2.29 | 3.46 | 0.95 |
| SD |  |  | 0.64 | 0.27 | 0.22 | 0.40 | 0.25 | 0.36 | 0.33 | 0.35 | 0.26 | 0.07 | 0.16 | 0.33 | 0.05 |
| Min |  |  | 16.70 | 3.80 | 3.60 | 7.10 | 6.60 | 4.90 | 6.30 | 2.50 | 1.90 | 1.60 | 2.00 | 2.90 | 0.90 |
| Max |  |  | 18.70 | 4.60 | 4.50 | 8.70 | 7.60 | 6.20 | 7.80 | 3.60 | 2.70 | 2.00 | 2.50 | 4.20 | 1.20 |

**Table 2.** Morphometric characters and summary statistics of the females of type series of *Epipedobates currulao* sp. nov. Measurements (in mm) of adult females; values are given as mean, SD, and range. See Methods section for explanation of abbreviations. *Holotype.

| Museum Code | Collector Code | Sex | SVL | FAL | HaL | TL | FL | HW | HL | NED | EW | END | IND | IOD | TD |
| --- | --- | --- | --- | --- | --- | --- | --- | --- | --- | --- | --- | --- | --- | --- | --- |
| ANDES:A:5255* | MBC754 | Female | 18.50 | 4.60 | 3.90 | 8.10 | 6.90 | 5.70 | 6.80 | 3.00 | 2.30 | 1.70 | 2.30 | 3.70 | 0.90 |
| ANDES:A:5257 | AJC07733 | Female | 19.50 | 4.50 | 4.50 | 8.70 | 7.20 | 6.00 | 6.70 | 3.20 | 2.60 | 2.00 | 2.40 | 4.20 | 1.10 |
| ANDES:A:5262 | AJC07737 | Female | 18.30 | 4.40 | 3.90 | 7.70 | 6.70 | 6.10 | 7.00 | 3.20 | 2.40 | 1.80 | 2.00 | 3.60 | 1.10 |
| ANDES:A:5264 | AJC07739 | Female | 17.50 | 4.40 | 3.90 | 8.20 | 7.20 | 5.70 | 7.00 | 2.90 | 2.40 | 1.90 | 2.40 | 3.60 | 0.90 |
| MVZ:Herp:295433 | AJC07740 | Female | 19.10 | 4.30 | 4.00 | 8.00 | 7.20 | 6.20 | 7.10 | 3.10 | 2.30 | 1.90 | 2.40 | 3.40 | 1.20 |
| MVZ:Herp:295435 | AJC07743 | Female | 18.90 | 4.60 | 4.20 | 8.60 | 7.60 | 5.80 | 7.10 | 2.90 | 2.20 | 1.70 | 2.10 | 3.40 | 1.10 |
| MVZ:Herp:295432 | RDT0934 | Female | 19.10 | 4.30 | 4.00 | 8.00 | 7.00 | 5.90 | 7.80 | 2.90 | 2.40 | 1.60 | 2.40 | 3.50 | 0.90 |
| ANDES:A:5258 | AJC07734 | NA | 18.60 | 4.40 | 3.60 | 7.30 | 6.60 | 6.00 | 6.50 | 3.40 | 2.30 | 1.90 | 2.30 | 3.90 | 1.00 |
| Mean |  |  | 18.70 | 4.44 | 4.06 | 8.19 | 7.11 | 5.91 | 7.07 | 3.03 | 2.37 | 1.80 | 2.29 | 3.63 | 1.03 |
| SD |  |  | 0.66 | 0.13 | 0.22 | 0.35 | 0.29 | 0.20 | 0.35 | 0.14 | 0.13 | 0.14 | 0.17 | 0.28 | 0.13 |
| Min |  |  | 17.50 | 4.30 | 3.90 | 7.70 | 6.70 | 5.70 | 6.70 | 2.90 | 2.20 | 1.60 | 2.00 | 3.40 | 0.90 |
| Max |  |  | 19.50 | 4.60 | 4.50 | 8.70 | 7.60 | 6.20 | 7.80 | 3.20 | 2.60 | 2.00 | 2.40 | 4.20 | 1.20 |

**Table 3.** Characteristics of morphology, coloration (in life), and calls of *E. currulao* sp. nov. compared to other species of *Epipedobates* found in Colombia. For comparison of morphological structures, we used specimens deposited at the Instituto de Ciencias Naturales of the Universidad Nacional de Colombia. For *E. narinensis* we reviewed the holotype (ICN-A:53344) and type series (ICN-A:53336–53340, 53342–53346). For *E.* aff. *espinosai,* we reviewed one specimen available at ICN (ICN-A:32504); additional body size data were provided by López-Hervas et al. (2024). The table is presented in parts across the next several pages for readability.

| Species | Snout-Vent Length (mm)<br>(mean ± SD) | Finger Length<br>h II>III | Fingers III and V<br>reduced | Finger IV swelling<br>in adult males | Webbing: Toe II<br>Preaxial | Webbing: Toe III<br>Preaxial | Webbing: Toe IV<br>Preaxial | Webbing: Toe IV<br>Postaxial |
| --- | --- | --- | --- | --- | --- | --- | --- | --- |
| *E. currulao* | Females: 18.47 ± 0.91, N = 14<br>Males: 16.98 ± 0.71, N = 16<br>Adults: 18.09 ± 0.50, N = 8 | Yes | Yes | Yes | Absent | Absent | Present, extends beyond the basal subarticular tubercle on toe IV | Absent |
| *E. narinensis* | Females: 17.08 ± 0.77, N = 3<br>Males: 16.82 ± 0.26, N = 5 | Yes | Yes | Yes | Present, reduced | Absent | Present, extends to or just below the basal subarticular tubercle on toe IV | Absent |
| *E. boulengeri*-Gorgona | Females: 20.13 ± 0.20, N = 3<br>Males: 19.15 ± 0.75, N = 4<br>Adults: 19.88 ± 1.81, N = 2 | Yes | No | Yes | Absent | Absent | Present, extends beyond the basal subarticular tubercle on toe IV | Absent |
| *E. boulengeri*-Nariño | Females: 17.57 ± 0.36, N = 8 | Yes | No | Yes | Absent | Absent | Present, extends beyond the basal subarticular tubercle on toe IV | Absent |
| *E. aff. espinosai* | Females: 17.68 ± 1.54, N = 11<br>Males: 16.32 ± 1.63, N = 7<br>Adults: 17.41 ± 1.58, N = 9 | Yes | No | Yes | Absent | Absent | Present, extends beyond the basal subarticular tubercle on toe IV | Absent |

| Species | Relative Finger Lengths | Relative Toe Lengths | Metatarsal Fold | Dorsal skin texture | Dorsal color | Oblique Lateral Stripe color | Oblique Lateral Stripe morphology | Ventrolateral stripe |
| --- | --- | --- | --- | --- | --- | --- | --- | --- |
| *E. curulao* | IV>II>III>V | IV>III>V>II>I | Absent | Finely granulated | Uniformly brown dorsal coloration, lacking markings | Present; white, yellow, or orange-yellow turning copper as it reaches the eye | Extends to eye or close to eye; complete in most but sometimes interrupted or diffuse | Present but poorly defined in some; white to blueish-white |
| *E. narinensis* | IV>II>III>V | IV>III>V>II>I | Absent | Finely granulated | Uniformly dark olive-green, lacking markings | Present; yellow-green | Extends only to mid body; diffuse and poorly defined | Present but indistinct in some; light green |
| *E. boulengeri*-Gorgona | IV>II>III>V | IV>III>V>II>I | Absent | Finely granulated | Brown or reddish brown, with dark spots | Present; creamy yellow | Extends to eye or close to eye; complete and clearly defined | Present and clearly defined in some; creamy white |
| *E. boulengeri*-Nariño | IV>II>III>V | IV>III>V>II>I | Absent | Finely granulated | Reddish brown with dark spots | Present; pale orange-gold; thinner than Gorgona population | Extends to upper eyelid; complete but slightly diffuse | Present and clearly defined in some; white |
| *E. aff. espinosai* | IV>II>III>V | IV>III>V>II>I | Absent | Finely granulated | Dark brick red to dark brown; some with dark spots | Present; white close to groin then turning copper as it reaches mid body | Extends to scapula or eye; complete in some but broken or diffuse in others | Present and clearly defined in some; white |

| Species | Ventral background coloration | Ventral pattern | Lateral shape of snout | Dorsal shape of snout | Tympanum/Eye (%) | HW/SVL (%) | HW/HL (%) | Blotch on anteriodorsal side of thigh | Upper arm color |
| --- | --- | --- | --- | --- | --- | --- | --- | --- | --- |
| *E. curculao* | Varies from pale blue turquoise to white; some with diffuse yellow on sides | Typically, heavily mottled with irregular black blotches | Protruding | Slightly rounded | Female s: 37.5 – 66.0<br>Males: 33.3 – 55.3<br>Adults: 43.1 – 61.6 | Female s: 27.7 – 33.3<br>Males: 29.1 – 36.3<br>Adults: 27.9 – 32.8 | Females: 75.6 – 89.6<br>Males: 73.1 – 103.7<br>Adults: 68.4 – 92.3 | Present and clearly defined: yellow to orange-yellow blotch | Bright yellow or whitish yellow flash marks |
| *E. narinensis* | Yellowish to greenish, bright in some but mostly dull | Speckled or slightly mottled with black | Protruding | Slightly rounded | Female s: 47.6 – 58.9<br>Males: 55.3 – 58.1 | Female s: 31.9 – 32.2<br>Males: 29.8 – 33.7 | Females: 82.1 – 86.3<br>Males: 79.3 – 87.6 | Absent or diffuse yellow-green coloration | Mostly absent, but a few with diffuse yellow coloration |
| *E. boulengeri*-Gorgona | Pale blue turquoise to nearly white | Irregular black blotches and vermiculations | Protruding | Slightly rounded | Female s: 50.0 – 57.1<br>Males: 44.8 – 52.0<br>Adults: 55.5 – 59.4 | Female s: 28.7 – 32.0<br>Males: 30.0 – 33.1<br>Adults: 29.3 – 32.7 | Females: 90.0 – 103.2<br>Males: 89.1 – 93.9<br>Adults: 80.4 – 89.7 | Present: yellow-white; much less yellow than in *E. curculao* | Mostly absent, but a few with diffuse whitish-yellow coloration; much less yellow than in *E. curculao* |
| *E. boulengeri*-Nariño | Bluish white | Mottled or with isolated black spots | Protruding | Slightly rounded | Female s: 45.0 – 64.0 | Female s: 25.7 – 32.2 | Females: 76.4 – 86.6 | Absent, or, less common: diffuse whitish-yellow (present in 2/8 individuals) | Mostly absent, but a few with diffuse cream coloration |
| *E. aff. espinosai* | Pale blue turquoise to nearly white or cream | Irregular black blotches and vermiculations; rarely without | Protruding | Slightly rounded | NA | NA | NA | Absent, or, less common: white or whitish-blue (present in 8/28) | Mostly absent, but a few with diffuse copper or cream |
|  |  | mottling<br>or with<br>yellow<br>markings |  |  |  |  |  | individual<br>s) | colorati<br>on |

| **Species** | **Peak frequency**<br>(mean ± SD, in<br>kHz)<br>(Range) | **Number of pulses**<br>(mean ± SD)<br>(Range) | **Call duration (s)**<br>(mean ± SD, in s)<br>(Range) | **Number of calls in**<br>**call series**<br>**(Range)** |
| --- | --- | --- | --- | --- |
| *E. currulao* (N =<br>15 frogs) | 5.23 ± 0.11<br>(4.98 – 5.47) | 73.94 ± 18.78<br>(22.0 – 122) | 2.21 ± 0.54<br>(0.67 – 3.88) | 1 |
| *E. narinensis* (N =<br>6) | 5.64 ± 0.21<br>(5.29 – 6.17) | 8.98 ± 2.45<br>(6.0 – 14.6) | 0.28 ± 0.070<br>(0.18 – 0.40) | 5–14 |
| *E. boulengeri* -<br>Gorgona (N = 6) | 4.76 ± 0.11<br>(4.56 – 4.90) | 14.48 ± 2.19<br>(9.5 – 20.0) | 0.30 ± 0.036<br>(0.21 – 0.37) | 2–3 |
| *E. boulengeri* -<br>Nariño (N = 5) | 5.51 ± 0.11<br>(5.23 – 5.68) | 13.09 ± 1.75<br>(10.6 – 17.0) | 0.28 ± 0.042<br>(0.23 – 0.37) | 3–6 |
| *E. aff. espinosai* | NA | NA | NA | NA |

**Table 4.** Summary statistics of spectral and temporal parameters of the advertisement calls of *Epipedobates currulao* sp. nov.

| Parameters | Call Duration (s) | Number of pulses | Pulse rate | Intercall interval (s) | Pulse duration (s) | Interpulse interval (s) | Low Frequency (kHz) | High Frequency (kHz) | Peak Frequency (kHz) | Bandwidth 90% (kHz) |
| --- | --- | --- | --- | --- | --- | --- | --- | --- | --- | --- |
| Mean | 2.213 | 73.938 | 34.894 | 62.675 | 0.017 | 0.013 | 4.845 | 5.586 | 5.237 | 0.354 |
| SD | 0.541 | 18.786 | 2.077 | 60.762 | 0.004 | 0.009 | 0.222 | 0.210 | 0.115 | 0.087 |
| Min | 0.674 | 22.000 | 26.000 | 15.120 | 0.009 | 0.001 | 4.236 | 5.221 | 4.981 | 0.215 |
| Max | 3.888 | 122.000 | 39.000 | 315.012 | 0.028 | 0.068 | 5.287 | 6.093 | 5.469 | 0.675 |

### Species comparison

At the type locality the new species occurs in sympatry with *Andinobates minutus* (Dendrobatidae, Dendrobatinae) but differs in body size and coloration (Fig. 3). *Andinobates minutus* has orange rather than yellow blotches on the anteriodorsal side of the thigh and dorsal side of the arms and its supralabial band is dark orange rather than white as in *E. currulao*. Ventrally in *A. minutus* the proportion of black and turquoise blue is similar. In contrast, in *E. currulao* the coloration is mainly turquoise blue with some black spots (Fig. 2C). In addition, adults of *A. minutus* are much smaller (mean = 12.00 mm, SD = 0.91 mm, N = 5) than adults of *E. currulao* (17.99 mm, SD = 0.95, N = 16), but *A. minutus* can be confused with *E. currulao* juveniles. The coloration of juvenile *E. currulao* is very similar to *E. currulao* adults, so color can still be used to differentiate the two species.

**Figure 3.**
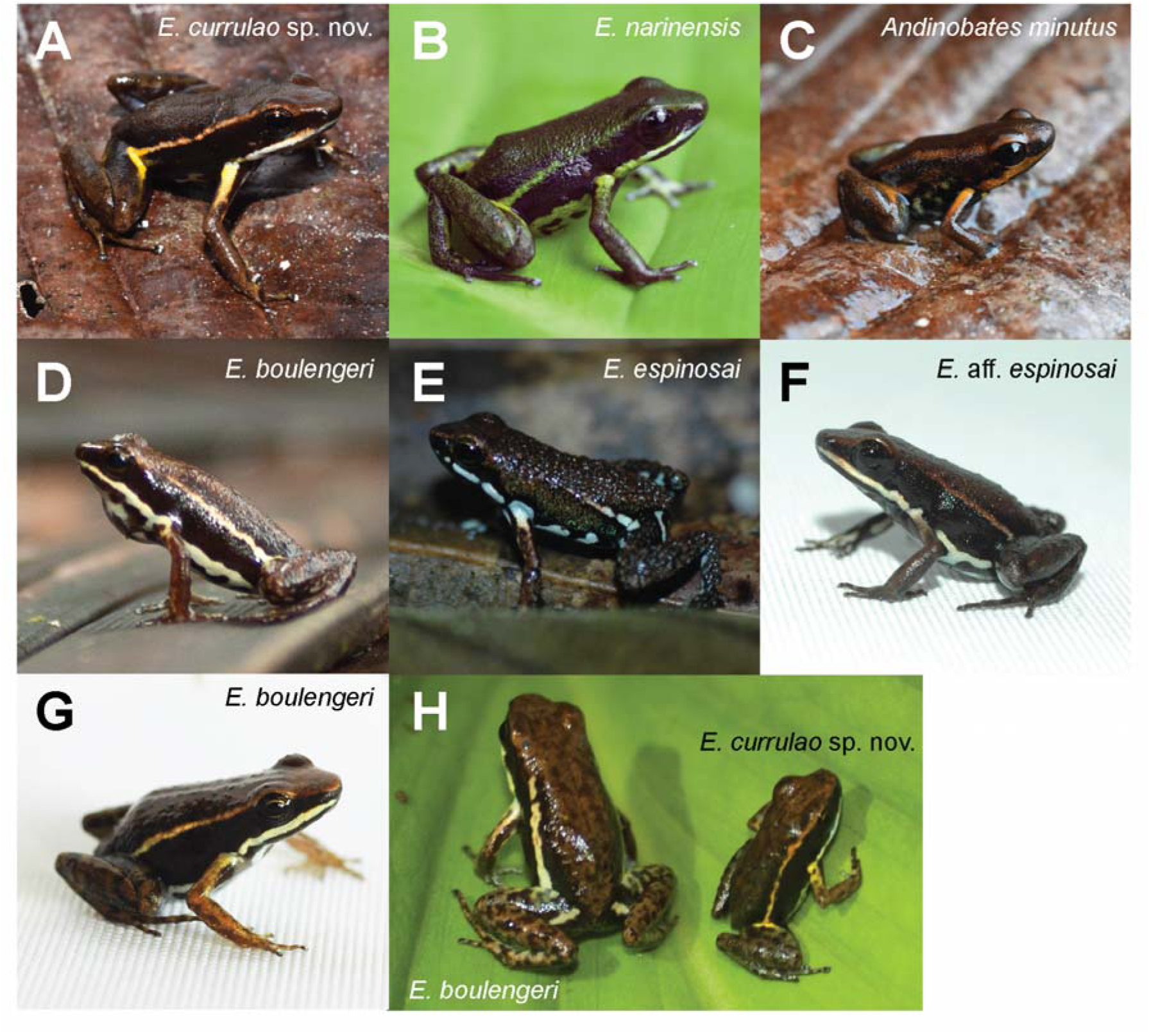
Images in life of *Epipedobates currulao* sp. nov. in comparison with close congeners and sympatric species. **A.** *E. currulao* sp. nov. from the type locality of Ladrilleros, Valle de Cauca, Colombia (ANDES:A:5261; SVL = 20.0 mm; adult male; paratype); **B.** *E. narinensis* from Biotopo, Nariño, Colombia (ANDES:A:3704; 16.39 mm; adult male); **C.** *Andinobates minutus* from Ladrilleros, Valle de Cauca, Colombia (ANDES:A:5266; 13 mm; sex not determined); **D.** *E. boulengeri* from Isla Gorgona, Cauca, Colombia (individual not captured); **E.** *E. espinosai* from Río Palenque, Santo Domingo de los Tsáchilas, Ecuador (individual not captured); **F.** *E.* aff. *espinosai* from La Nutria, Nariño, Colombia (ANDES:A:2476; 17.77 mm; sex not determined); **G.** *E. boulengeri* from Maragrícola, Nariño, Colombia (ANDES:A:2472; 18.96 mm; adult female); **H.** A side-by-side image of *E. boulengeri* from Isla Gorgona (ANDES:A:3695; 20.65 mm; adult female) and *E. currulao* sp. nov. from Pianguita, Valle de Cauca, Colombia (ANDES:A:3690; 16.42 mm; adult female) demonstrating the large size difference between the two species. All images were taken by RDT except for A and C, which were taken by JCR. Photos are not to scale.

Previously, *E. currulao* has been confused with *E. boulengeri.* The two species can be differentiated by the color (in life) of the blotch on the anteriodorsal side of the thigh and the color of the blotch on the dorsal side of the arm close to the axilla, which are both yellow to orange in *E. currulao* and white to whitish yellow in *E. boulengeri*. Some individuals of *E. currulao* have diffuse yellow coloration along the lateral edge of the venter; this coloration has not been identified in any *E. boulengeri* population to date. Compared to the Isla Gorgona *E. boulengeri* population, *E. currulao* has a much thinner oblique lateral stripe; however, this stripe is similar in morphology to the mainland population of *E. boulengeri.* The mainland population of *E. boulengeri* (based on images from Maragrícola) can be differentiated from *E. currulao* by the size, shape, and color of the blotch on the anteriodorsal side of the thigh, which is large, yellow or orange, and clearly defined in *E. currulao* but absent or diffuse and transparent or whitish copper in *E. boulengeri* (Maragrícola). In addition, *E. currulao* has advertisement calls with one call per series while *E. boulengeri* has 2–3 calls (Isla Gorgona) or 3–6 calls per series (Maragrícola) (Table 3).

Individuals of *E. narinensis* and *E. currulao* can be differentiated by the dorsal color, which is olive-green in *E. narinensis* and dark brown in *E. currulao*, the length of the oblique lateral line, which extends to the eye in *E. currulao* but only extends to the mid-body in *E. narinensis*, the shape of the blotch on the anteriodorsal side of the thigh, which is clearly defined in *E. currulao* but diffuse in *E. narinensis*, and the background color of the venter, which is yellowish to greenish in *E. narinensis* and pale blue to turquoise or white in *E. currulao*. The structure of the advertisement call of the species also differs, where calls in *E. currulao* occur one per series but 5–14 per series for *E. narinensis* (Table 3).

Individuals of *E.* aff. *espinosai* and *E. currulao* can be differentiated by the length of fingers III and V, which are reduced in *E. currulao* but not in *E.* aff. *espinosai* (see Morphology in the Systematics section below). The oblique lateral line most often extends to the eye in *E. currulao* but tends to end at the scapula in *E.* aff. *espinosai*. The shape of the blotch on the anteriodorsal side of the thigh is clearly defined and yellow or orange in *E. currulao* but absent or small and whitish-blue in *E.* aff. *espinosai*. The blotch on the dorsal side of the arm near the axilla is bright yellow or whitish yellow while the blotch is mostly absent or with diffuse copper or cream coloration in *E.* aff. *espinosai*.

A summary of the morphological characters of *E. currulao* sp. nov. compared to other *Epipedobates* species in Colombia is shown in Table 3. Please see Suppl. material 4 for morphological measurements of individuals and Suppl. material 5 for more extensive notes on color variation within and among species.

### Description

#### Coloration of holotype in life (**Fig. 2A–D**)

Dorsal surfaces dark brown with an oblique lateral stripe extending from the posterior region of the eye to the groin, with metallic orange-yellow coloration in anterior region becoming yellow towards the groin. Groin dark brownish black with a distinct yellow blotch continuing to the anteriodorsal surface of the thigh. Flanks black. Forelimb and hindlimb background dark brown with irregular dark brown spots. Anterior region of the upper arm with yellow blotch similar in color to thigh blotch. Supralabial stripe creamy white, extending from nares to axilla. Ventral surfaces turquoise blue with irregular black spots. Iris copper. Two elongated black spots in the gular region (Fig. 2C).

#### Coloration of holotype in preservative (**Fig. 2E–K**; after two years of preservation in 70% ethanol)

Dorsally black to dark brown, hind limbs dark brown. Lighter brown forelimbs with some dark brown spots. Dorsal blotches yellow in life on forelimbs and thighs become white in preservation. Ventrally pale white with irregular black to dark brown spots. Groin black. Dark gray oblique lateral line extending from the posterior region of the eye to the groin. Light gray supralabial stripe running from the tip of the face to the axilla.

#### Coloration variation of type series and other populations in life

All individuals in the type series exhibit a uniform dark brown color on the dorsum (Suppl. material 3). The dorsum, head, thigh, and shanks feature a skin texture covered with small and scattered tubercles. Flanks and concealed surfaces of forelimbs and hindlimbs smooth and solid black. All individuals show an oblique lateral stripe that stretches from the posterior region of the eye to the groin and differs slightly among populations. In the Ladrilleros population (type locality), the oblique lateral stripe displays an orange-yellow color with indistinct edges and stretches from the front part of the groin to the eye and the canthus rostralis, gradually changing from yellow to a more brownish-orange hue. In the Anchicayá population, the oblique lateral stripe takes on a yellowish hue and, in most (6/8) specimens, it tapers or fragments into patches, gradually disappearing before reaching the eye. In some individuals from Pianguita (3/5), the oblique lateral stripe exhibits regular interruptions along its entire length.

The groin region is characterized by a dark brown-black color, with a noticeable yellow blotch that extends to the front inner thigh (Suppl. material 3). Most individuals have a paracloacal spot similar in coloration, but smaller and more elongated, on the posterior dorsal surface of the thigh. In the Anchicayá population, these spots are present in only some individuals. Most individuals have a yellow blotch on the dorsal region of the arm that matches the color of the blotch on the anteriodorsal side of the thigh. All individuals exhibit a creamy white to pale turquoise blue upper labial stripe that is notably lighter than the oblique lateral stripe. In some cases, this stripe has a slight iridescent quality and extends from below the nostril to the axilla. Also, in certain individuals, it continues posteriorly as a vaguely defined ventrolateral stripe. The ventral surfaces of the throat, belly, and thighs exhibit a pale turquoise blue to white color with irregular black spots and patterns resembling worm-like lines (vermiculations). In males, the background coloration may be darkened by a diffuse gray pigment located just anterior to the pectoral region and the vocal sac. A few individuals display a diffuse yellowish coloration towards the outer edges of the belly. Full-page plates of images of four *E. currulao* populations can be seen in the supplementary figures 4–6 of López-Hervas et al. (2024) and plates showing images of the type series described herein can be found in Suppl. material 3.

#### Vocalizations

Calls of *E. currulao* sp. nov. consisted of 22–122 pulses per call (mean = 73.98, SD = 18.77, N = 15), with a call duration of 0.67–3.88 s (mean = 2.21, SD = 0.54 s, N = 15). Pulse rate consisted of 26–39 pulses by second (mean = 34.89, SD = 2.07, N = 15), pulse duration of 0.009–0.028 (mean = 0.017, SD = 0.004 s, N = 15) and interpulse interval of 0.001–0.068 s (mean = 0.013, SD = 0.009 s, N = 15). Intercall interval ranges from 15.12–315.01 s (mean = 62.67, SD = 60.76 s, N = 15) (Table 4). The peak frequency ranged between 4.98 and 5.46 kHz (mean = 5.23, SD = 0.11 kHz), and the frequency interquartile bandwidth between 0.21 and 0.67 kHz (mean = 0.35, SD = 0.87 kHz) (Table 4). Calls are not frequency modulated. The amplitude of the first and last pulses is reduced by 7– 10% compared to the rest of the pulses.

#### Etymology

The specific epithet *currulao* is a noun in apposition of masculine gender. It refers to the musical genre that originated on the southern Pacific coast of Colombia and Ecuador, where *E. currulao* occurs and also contributes to the local soundscape. Currulao, also known as bambuco viejo, is an Afro-Colombian sounded practice that inspires dancing and transmits the happiness and cultural tradition of this region. It is a symbol of resilience in the face of racial and regional oppression (Abadía 1973; Aristizabal 2002; Birenbaum Quintero 2006, 2019). We named this species in honor of, and as an homage to, this musical genre that represents the culture of the southern Colombian Pacific because: “la música, como la vida, no se pueden dejar perder”, which translates to “music, like life, cannot be allowed to be lost” (Cruz Hoyos 2016).

### Distribution

*Epipedobates currulao* sp. nov. occurs in the Department of Valle del Cauca in the Pacific lowlands of southwestern Colombia. These frogs inhabit lowland forests from 0–260 m. The type locality is Ladrilleros, Buenaventura, Valle del Cauca, Colombia. We also observed the species in areas close to the type locality including Corregimiento Pianguita and Corregimiento Juan Chaco (La Barra beach) in the municipality of Buenaventura. The distribution towards the western flank of the western mountain range is in the Vereda El Danubio, the upper basin of the Anchicayá River, Dagua, Valle del Cauca. If we assume that the coloration traits of the new species are consistent for all populations of this species, records from iNaturalist would extend the distribution of *E. currulao* 194 km (132 km) straight line south to the municipality of Timbiquí, Cauca (see iNaturalist observations No. 135253843 and 85214439 and Fig. 1). Because individuals of this species were previously assigned to *E. boulengeri*, we recommend further exploration and inspection of museum specimens to better understand the geographic distribution of this species.

### Ecology

*Epipedobates currulao* sp. nov. is a terrestrial species found on the ground during the day in agroforestry areas, on the edges of secondary forests, or in small patches of disturbed secondary forest always near or within marshes and/or slow-flowing streams (Fig. 4). The species is also likely present in primary forest in the region. We observed individuals actively moving among the grass and leaf litter, or actively calling at the edges of water bodies. The type locality (Ladrilleros, Valle de Cauca, Colombia) includes tiny forest fragments among human dwellings. Usually, these areas are contaminated with garbage or agricultural residues (Fig. 4A–F). In Anchicayá, populations of *E. currulao* are usually found along roadsides and forest edges, whenever small streams and leaf litter are present (Fig. 4G–J). We observed greater vocal activity during the morning and in the late afternoon than during warmer or sunnier parts of the day.

**Figure 4:**
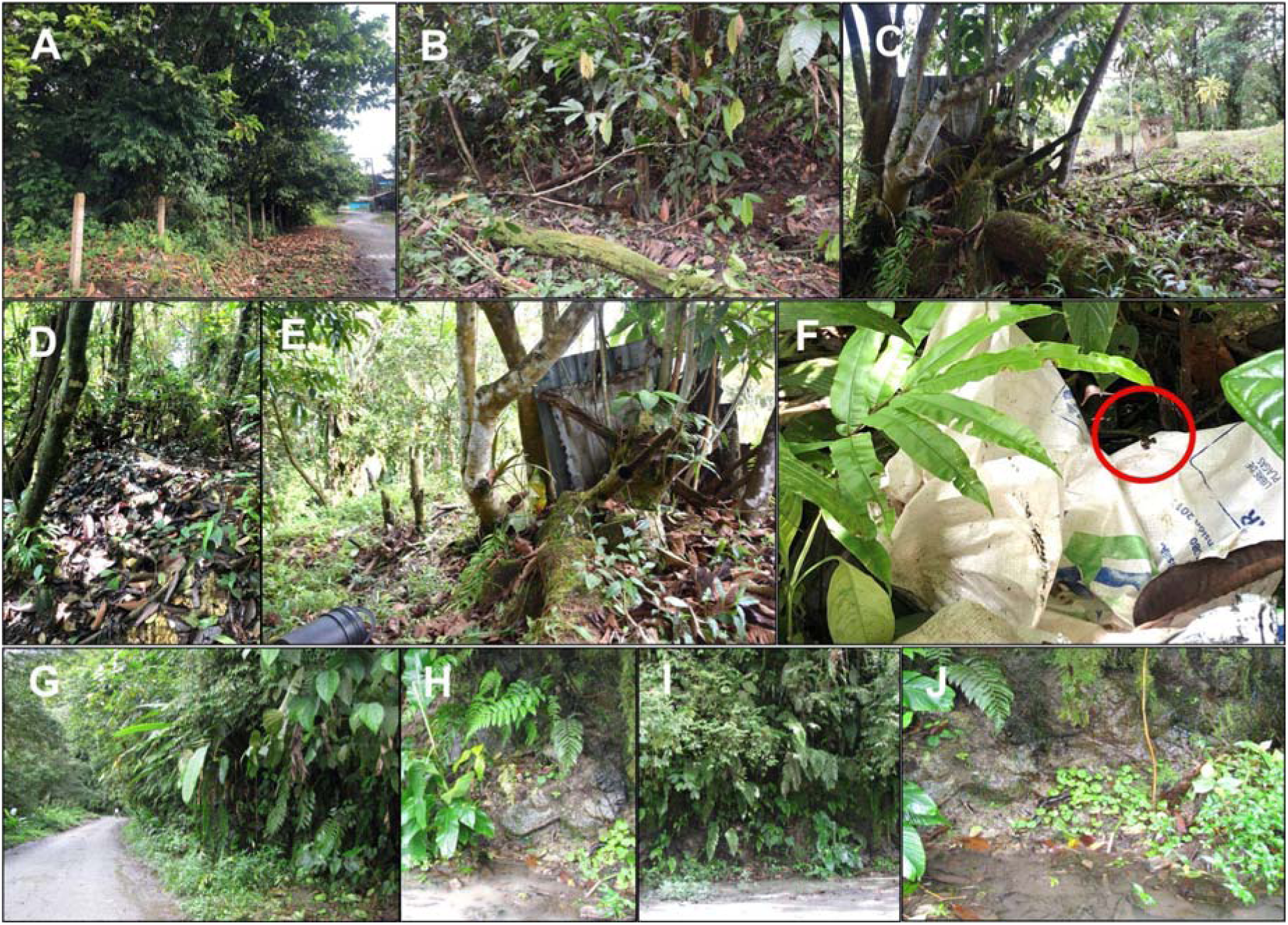
Habitat structure of *Epipedobates currulao* sp. nov. at two localities. **A–C** Images of the type locality at Ladrilleros, Buenaventura, Valle del Cauca, Colombia. Usually, this species is found on roadsides near streams formed by rainfall. **D–F** At the type locality, we observed frogs in habitats contaminated with garbage or agricultural waste. Note the frog in F (red circle). **G–J** Habitat characteristics in Anchicayá, Dagua, Valle del Cauca, Colombia. Images B, C and E were taken by RDT, all others by MBC.

### Conservation status

The populations observed in the four localities (Ladrilleros, Pianguita, La Barra, and Anchicayá) are probably affected by housing construction, garbage, or agricultural waste as well as forest fragmentation and loss of habitat. We do not know if these populations have adapted to human disturbance or if they are remnants of the original distribution of the species, but congeners in Ecuador are abundant in highly modified habitats such as cacao and banana populations (though they are notably absent from African palm plantations; RDT pers. obs.). Although we consider that *E. currulao* sp. nov. is moderately abundant at the type locality, as three of us (RDT, MBC, JCR) captured 17 individuals in an approximate area of 30 m2 in four hours, we do not know their abundance in less disturbed habitats. Improving our understanding of its conservation status will require monitoring and explorations in potential localities in Cauca and Valle del Cauca, especially in protected areas close to the localities reported in this study.

Despite the lack of certainty regarding the distribution of *E. currulao* sp. nov., its extent of occurrence is likely less than 20,000 km^2^. We know the species occurs in at least four localities (Ladrilleros, Anchicayá, Pianguita, and La Barra), with one more pending genetic validation (Timbiquí). The forests in the range of this species have been, and probably will continue to be, subject to strong deforestation pressures that reduce the quantity and quality of available habitat and increase its fragmentation. More data will be necessary to determine the *E. currulao* sp. nov. categorization, but here we provide some recommendations. Under the precautionary attitude, we would recommend that it be categorized as Vulnerable (VU: B1a, biii; IUCN, 2019) based on B1, an Extent of occurrence (EOO) <20.000 km^2^ (approx. 3600 km^2^ from Timbiquí to Ladrilleros) and (a) Severely fragmented OR Number of locations ≤ 10 and (b) Continuing decline observed, estimated, inferred or projected in any of: (iii) area, extent and/or quality of habitat. Under the evidentiary attitude, we would recommend that *E. currulao* sp. nov. be categorized as Near Threatened, recognizing that more studies may reveal additional populations and expand its known range and population size. In addition, we observed this species within the Farallones de Cali National Natural Park, so there is at least one known population (Anchicayá) within a protected area. The distribution of *E. currulao* sp. nov. is also very close to the Parque Nacional Natural Uramba Bahía Málaga. We are not aware of the presence of *E. currulao* sp. nov. within the park, but this conservation area protects more than 47,000 ha of marine and coastal areas, so it is very likely that *E. currulao* sp. nov. is found within the park. More research about this species’ distribution, ecological requirements, and population dynamics will help confidently assign it to a threat category.

## Systematics

### Molecular and phylogenetic analyses

We estimated a phylogeny using existing data (López-Hervas et al. 2024) and 10 new sequences from five *E. currulao* specimens from the type locality. The resulting phylogeny contains strong support for the monophyly of *E. currulao* and recovers known phylogenetic relationships among species in this genus (Fig. 5 and Suppl. material 6; López-Hervas et al. 2024). The sister species to *E. currulao* is *E. narinensis*, which occurs in the south-central part of the Pacific coast of Colombia. These two species constitute a clade of Epipedobates that is highly divergent and sister to a clade that contains all other species of the genus.

**Figure 5.**
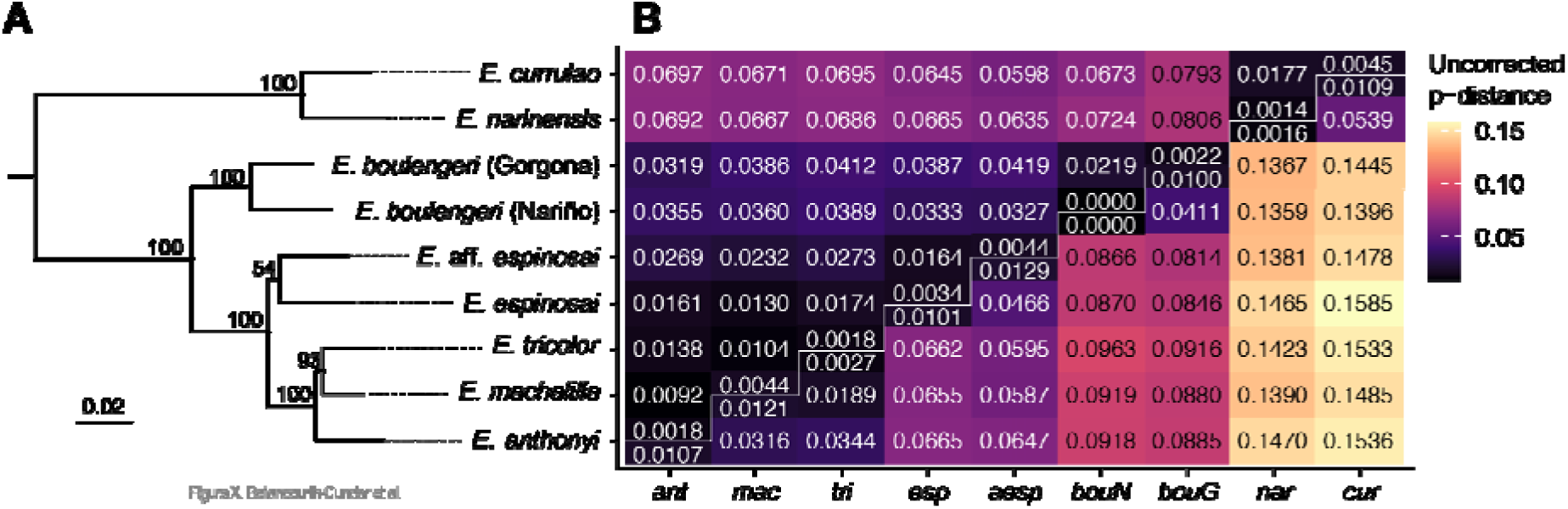
Phylogenetic position and genetic distances of *E. currulao* sp. nov. and other species of *Epipedobates*. **A** Species-level maximum-likelihood phylogeny of *Epipedobates* estimated in IQ-TREE2 using two nuclear genes (*BMP2, H3*) and three mitochondrial gene fragments (*CYTB, 12S–16S, CR*) (see Fig. Supp. material S2 for full phylogeny). Node support values from 10,000 ultrafast bootstrap replicates indicated strong support of *E. currulao* sp. nov. as the sister species of *E. narinensis*. **B** Matrix of mean pairwise p-distances within and among species of *Epipedobates*. The lower triangle distances are based on *CYTB* and the upper triangle distances are from *12S–16S*. Abbreviations: cur, *E. currulao* sp. nov; nar, *E. narinensis*; bouN, *E. boulengeri* (Nariño, Colombia); bouG, *E. boulengeri* (Isla Gorgona, Cauca, Colombia); aesp, *E.* aff. *espinosai*; esp, *E. espinosai*; tri, *E. tricolor*; mac, *E. machalilla*; ant, *E. anthonyi*. See Supp. material 6 for full phylogeny.

**Figure 6.**
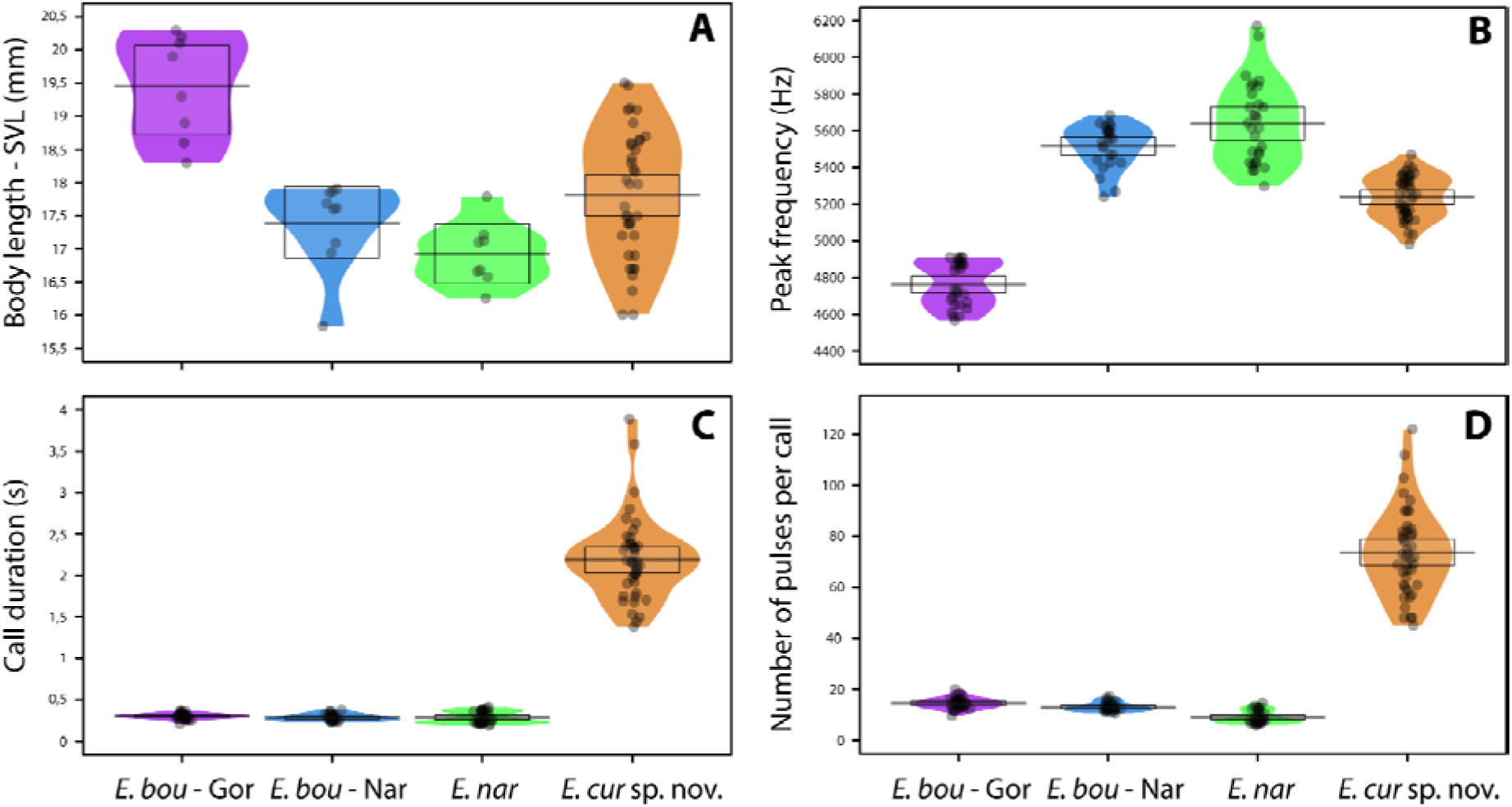
Comparisons of SVL and advertisement call traits between *E. currulao* sp. nov. and other species of *Epipedobates found in Colombia.* **A** Snout-vent length (SVL) **B** Peak frequency **C** Call duration and **D** Number of pulses per call. Abbreviations include: E. bou - Gor, *E. boulengeri* - Gorgona; E. bou - Nar, *E. boulengeri* - Nariño; E. nar*, E. narinensis;* and E. cur sp. nov., *E. currulao* sp. nov. Black points represent raw data, the horizontal bar shows the mean, the shaded diagram is a smoothed density curve showing the full data distribution, and the rectangle represents the uncertainty around the mean using a 95% Bayesian highest density interval.

The average p-distances among clades of Epipedobates species show that *E. currulao* is genetically most similar to *E. narinensis* (1.77% for 12S–16S and 5.39% for CYTB), as expected given the molecular phylogeny.

### Morphology

In *E. currulao* finger II is longer than finger III, and finger IV is swollen in males. In our revision of the morphology, we noted that fingers III and V are reduced in length in *E. currulao* (finger III/SVL: mean ± SD = 16.1% ± 1.59%; finger V/SVL: 15.1% ± 1.42%, N = 37 frogs) (Fig. 2H–I) and *E. narinensis* (finger III/SVL: 16.9% ± 0.85%; finger III/SVL: 16.2% ± 1.08%, N = 8) relative to *E. boulengeri*-Gorgona (finger III/SVL: 19.2% ± 0.67%; finger V/SVL: 18.2% ±1.12%, N = 9), *E. boulengeri*-Nariño (finger III/SVL: 17.5% ± 0.99%; finger V/SVL: 17.7% ± 2.14%, N = 9) or *E.* aff. *espinosai* (finger III/SVL: 19.3%; finger V/SVL: 16.3%, N = 1) (Table 3, Suppl. material 4).

There are significant differences in body size between *Epipedobates currulao* sp. nov., *E. narinensis*, and *E. boulengeri* (Gorgona and Nariño) (Kruskal-Wallis test: H3 = 19.058, P < 0.001; Fig. 6A). *Epipedobates boulengeri* - Gorgona exhibits significantly larger mean body size than the other populations analyzed. A Wilcoxon test showed that *E. currulao* sp. nov. is larger than *E. narinensis* (P = 0.05, N = 33) and smaller than *E. boulengeri* - Gorgona (P < 0.006, N = 32). We found no significant differences in body size between *E. currulao* sp. nov. and the populations of Nariño assigned to *E. boulengeri* (P = 0.462, N = 33). We found sexual dimorphism in body size (SVL) for *E. currulao* sp. nov. (ANOVA: F1,46 = 6.269, P < 0.01), where mean SVL in females was larger (mean = 18.24, SD = 1.09 mm, N = 28, Table 2) than the mean in males (mean = 17.45, SD = 1.07 mm, N = 20, Table 1).

### Vocalizations

*Epipedobates currulao* sp. nov., *E. narinensis*, and *E. boulengeri* (from Gorgona and Nariño) display noteworthy distinctions in call parameters. We found significant differences in the peak frequency of advertisement calls among all species and populations tested (Kruskal-Wallis test: H_3_ = 109.25, P < 0.001; Fig. 6B). Wilcoxon test showed that *Epipedobates currulao* sp. nov. calls with a lower peak frequency than *E. narinensis* (P < 0.001, N = 81) and *E. boulengeri*-Nariño (P < 0.001, N = 76), but its peak frequency is higher than for *E. boulengeri*-Gorgona (P < 0.001, N = 79). We found statistically significant differences in call duration (Kruskal-Wallis test: H_3_ = 95.302, P < 0.001; Fig. 6C). *Epipedobates currulao* sp. nov. exhibits longer calls than *E. narinensis* (P < 0.001, N = 81), *E. boulengeri*-Nariño (P < 0.001, N = 76) and *E. boulengeri*-Gorgona (P < 0.001, N = 79). Similar results were found for the number of pulses per call (Kruskal-Wallis test: H_3_ = 107.52, P < 0.001, Fig. 6D). *Epipedobates currulao* sp. nov. has calls with more pulses than *E. narinensis* (P < 0.001, N = 81), *E. boulengeri*-Nariño (P < 0.001, N = 76), and *E. boulengeri*-Gorgona (P < 0.001, N = 79).

Advertisement calls are composed of groups of calls, one call for *E. currulao* sp. nov., 2–3 for *E. boulengeri* - Gorgona, 3–6 for *E. boulengeri* - Nariño and 5–14 for *E. narinensis* (Fig. 7). Additionally, we recorded two males close to a female with vocalizations different from their advertisement call (Fig. 7E). These vocalizations, most likely a courtship call, ranged from 0.21–0.31 s in length (mean = 0.25, SD = 0.03 s, N = 2), and included 7–9 pulses per call (mean 8.25 ± SD 0.83). The peak frequency ranged between 4.78–5.17 kHz (mean = 4.96, SD = 0.13 kHz).

**Figure 7.**
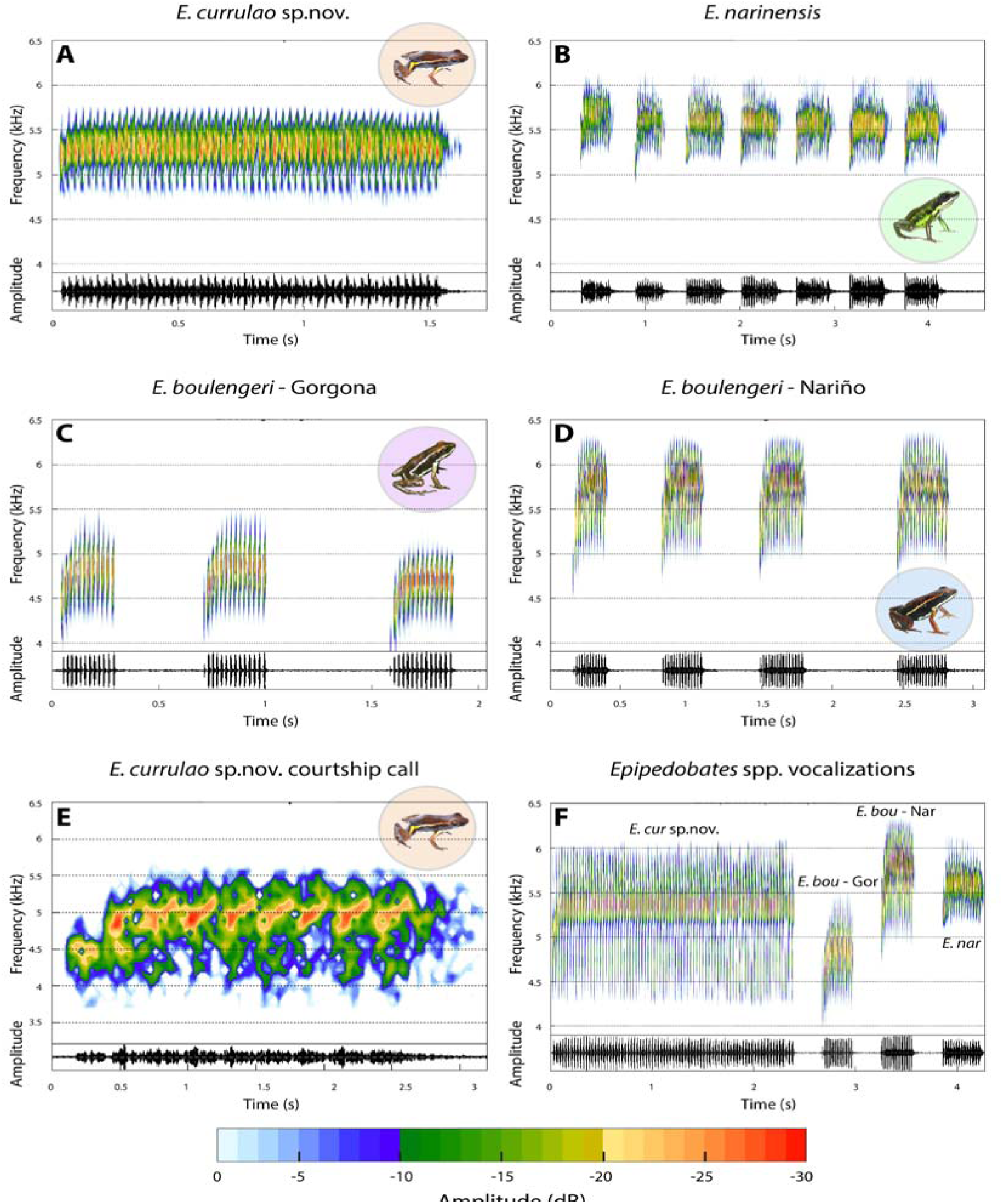
Vocalizations in *E. currulao* sp. nov. and other species of *Epipedobates*. Spectrograms (above) and oscillograms (below) showing general aspects of the structure of one spontaneous advertisement call for one male of **A** *E. currulao* sp. nov. of Ladrilleros, Valle del Cauca, **B** *E. narinensis* of Biotopo, Nariño, **C** *E. boulengeri*-Gorgona (Type locality) and **D** *E. boulengeri*-Nariño of Maragrícola, Nariño. **E** Spectrograms and oscillograms showing general aspects of the structure of one courtship call of *E. currulao* sp. nov. of Ladrilleros, Valle del Cauca. **F** Comparison of one advertisement call between *E. currulao* sp. nov. and other species of *Epipedobates* found in Colombia. *E. bou* - Gor: *E. boulengeri* - Gorgona, *E. bou* - Nar: *E. boulengeri* - Nariño, E. nar: *E. narinensis* and *E. cur* sp. nov.: *E. currulao* sp. nov. Fast Fourier Transform-FFT = 256, overlap = 90%.

A principal component analysis (PCA) of call parameters indicated that the first three principal components explain 88% of the variation in calls (Fig. 8A). PC1 explained 45% of the variation and was positively associated with call duration, number of pulses, and intercall interval. PC1 was also negatively related to high frequency and 90% bandwidth (Suppl. material S4). PC2 explained an additional 33% of the variation in call parameters. It was positively related to low frequency and peak frequency. PC3 explained 10% of the variation and was positively associated with frequency interquartile bandwidth 90% (Fig. 8A, Suppl. material 7). In summary, PC1 was mainly associated with temporal characteristics of the advertisement call and PC2 with spectral characteristics. The linear discriminant analysis (LDA) (Fig. 8B) function using five PCs showed that *E. currulao* can be easily differentiated from other species based on their advertisement calls. LD1 was positively associated with PC1 (temporal characteristics) (LD coef. = 1.71), and LD2 was negatively associated with PC2 (spectral characteristics) (LD coef. = -1.47). This means *E. currulao* is distinguished by having longer calls and a greater number of pulses (Figs 8B, 6CD). With respect to low and peak frequency, *E. currulao* calls are characterized by intermediate values compared to the other species analyzed (peak frequency: 4.84 ± 0.22 kHz, N = 15). *Epipedobates boulengeri-*Gorgona has the lowest peak frequency (4.76 ± 0.11 kHz, N = 5) and *E. narinensis* has the highest peak frequency (5.64 ± 0.21 kHz, N = 6) (Figs 8B and 6B).

**Figure 8.**
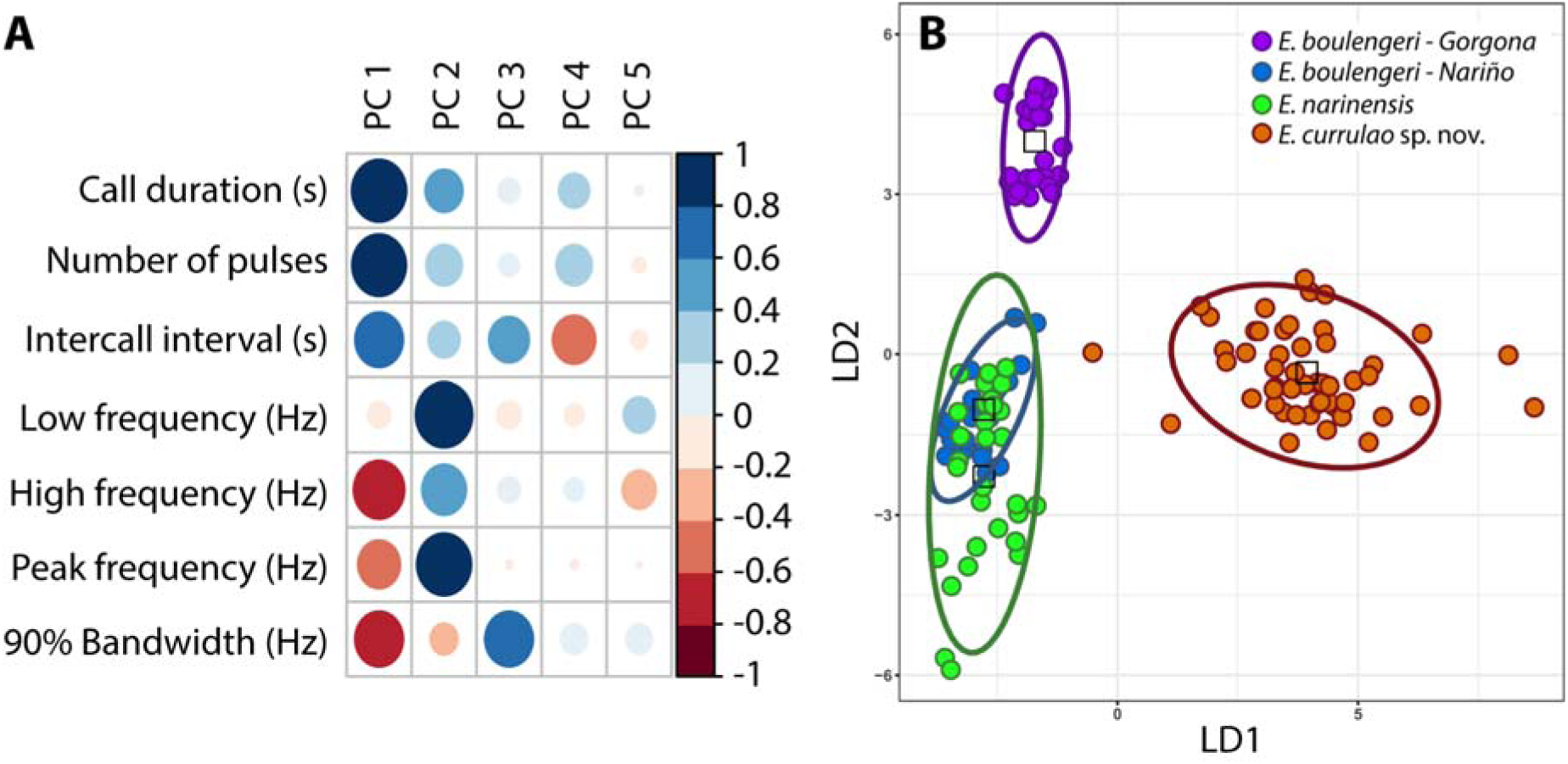
Differences in the acoustic parameters of *E. currulao* sp. nov. compared to other species of *Epipedobates found in Colombia*. **A** The relative contribution of each acoustic variable to the first five principal components obtained in the Principal Component Analysis. The absolute value of each contribution is represented according to the size of the circle, while blue and red colors show positive and negative contributions, respectively. **B** A Linear Discriminant Analysis using PCs as input indicates *E. currulao* can be differentiated from other species based on their advertisement calls, particularly regarding call duration and number of pulses (PC1 loads positively on LD1). Dots indicate the individuals, color indicates the species, and ellipses indicate the 95% confidence intervals for LDA data points.

## DISCUSSION

We describe *Epipedobates currulao* sp. nov. as a new species found along the Pacific chocoan rainforest of Colombia, which was previously considered to be part of the *E. boulengeri* species complex (López-Hervas et al. 2024). We describe this species as new based on prior work (López-Hervas et al. 2024) demonstrating its divergence from its sister species, *E. narinensis*, and its unique coloration. Here we provide additional evidence with characterization of its distinct advertisement call.

*Epipedobates currulao* is named in honor of the Afro-Colombian currulao musical genre and traditions that arose from generations of multicultural practices in the Pacific region, starting with African slaves brought to work in the gold mines, and continuing today with infusion of new ideas and interpretations (Birenbaum Quintero 2019; Guevara Calderón and Godoy Acosta 2015). As part of the cultural heritage of Black communities in the Pacific region, currulao music, also known as bambuco viejo, helped bolster arguments that led to the recognition of the land rights and cultural identity of Afro-Colombians through Law 70 of 1993 (Birenbaum Quintero 2019).

More information will be necessary to evaluate the conservation status of *E. currulao*, but we propose that it may be considered as either Vulnerable or Near Threatened under IUCN categorization given its relatively small range. Although *E. currulao* is not known to be sympatric with other species of *Epipedobates*, regions in southwestern Colombia (especially east of Guapi) have not been extensively surveyed and may contain sites with sympatry between *E. currulao*, *E. boulengeri*, and/or *E.* aff. *espinosai*. *Epipedobates currulao* does occur in sympatry and can be confused with *Andinobates minutus* unless color patterns are carefully distinguished (see Differential diagnosis section above). The other species of the *Epipedobates* genus are found along the western coast and Andean foothills in Ecuador and Peru. Although one species (*Epipedobates maculatus*) is thought to occur in western Panama (Grant et al. 2017; Jungfer 2017), its assignment to this genus has been brought into question (López-Hervas et al. 2024).

We find that the advertisement call of *E. currulao* is unique compared to other *Epipedobates* distributed in Colombia. Previously, Lötters et al. (2003) reported advertisement call differences between *Epipedobates boulengeri* recorded at Anchicayá (now *E. currulao*) and at Lita (now *E.* aff. *espinosai*), suggesting at the time that the populations may belong to a species complex. In our recordings, we found that *E. currulao* makes only one call per series, although Lötters et al. (2003) describe call series with up to three notes (notes are calls according to our definition); however, they recorded only one male.Visually, when looking at the Lötters et al. (2003) recording, it appears that the call is divided into groups of notes. However, when comparing the inter-note times (0.020-0.025 s) of the call, we note that these are similar to our measurements for inter-pulse times (0.001– 0.068 s; mean = 0.013, SD = 0.009 s, N = 15). Thus, we consider that these “notes” comprise a single call, and that the call series of *E. currulao* contains one relatively long call with many pulses.

We provide data on the courtship call of *E. currulao*, which to our knowledge is the first courtship call reported for *Epipedobates*. Like courtship calls of other neotropical poison frogs, it is lower frequency than the advertisement call of *E. currulao* (Caldeira Costa et al. 2006; González-Santoro et al. 2021; Marques Correia da Rocha et al. 2018; Moss et al. 2023). It was also shorter and had fewer pulses (7.9 vs 22–122 per call).

The density of *Epipedobates* varies across species and sites in Colombia. During our fieldwork, we noticed that the populations of *E. boulengeri* at Isla Gorgona and Maragrícola are quite abundant. For example, more than five individuals can be heard calling at the same time, so it was very difficult to record a single male. For *E. narinensis* at Biotopo natural reserve, individuals were calling less densely but were still abundant, with 8–10 m distance between calling males. But in other nearby locations we have heard only two or three individuals in a 3–4-hour walk. The individuals of *E. currulao* sp. nov. are more spread out and males were observed to call at approximately 4 to 5 m distance from each other at the type locality (Ladrilleros) and nearby at La Barra. At Anchicayá, males call every 3–4 m and are associated with small streams at the edge of the unpaved road. We do not know the reason for differences in density, as the pattern does not necessarily correspond to differences in habitat quality. While Isla Gorgona is a national park, *E. boulengeri* appears to be more abundant in the disturbed areas adjacent to field station buildings. The Maragrícola site where we sampled *E. boulengeri* in Nariño is an experimental cacao plantation. Ladrilleros and La Barra are disturbed sites similar to where *E. boulengeri* was abundant at Isla Gorgona, yet *E. currulao* did not occur at such high densities. It is plausible that the *E. currulao* is more vulnerable to habitat transformation than *E. boulengeri*, but additional research is necessary to understand what drives differences in density across sites and species.

Unfortunately, we did not collect larval stages of *E. currulao.* Larval morphology of Dendrobatoidea has contributed to understanding of the phylogenetic relationships within the superfamily (Anganoy-Criollo and Cepeda-Quilindo 2017). It would be interesting to collect tadpoles of the new species for comparison with other Epipedobates species. Comparisons of larval stages made by Anganoy-Criollo and Cepeda-Quilindo (2017) detected some differences between the populations of Ecuador and southwest of Colombia in body shape, tips of marginal papillae, and pattern of tail coloration. However, the specimens from Colombia used in their study likely correspond to *E. boulengeri* - Nariño (*E. boulengeri* sensu stricto) and we are not certain of the identification of the specimens from Ecuador, which could correspond to *E.* aff. *espinosai*, *E. boulengeri* sensu stricto, or *E. espinosai*, which all occur in the region (López-Hervas et al. 2024).

## ACKNOWLEDGEMENTS

RDT and MBC thank Cristian Flórez Pai, Luis Alfredo Esteves, Fray Arriaga, Sandra V. Flechas, Daniel Nastacuaz and Nataly Portillo for assistance in fieldwork. Adolfo Amézquita facilitated the permitting process in 2014 and 2016 and Mireya Osorio R. in 2022. We also thank ANDES, QCAZ, and MVZ museum personnel including Angela Sánchez G. and Carol Spencer for their assistance in accessioning specimens. We thank Valeria Ramírez-Castañeda for assistance with PCR. We thank Mauricio Rivera for access to the specimens of the Instituto de Ciencias Naturales at Universidad Nacional de Colombia, Bogotá, and Nataly Casas for her help in taking photographs of museum specimens. We thank Dr. Stefan Lötters and Dr. Andrés F. Jaramillo-Martínez and Dr. Fedor Konstantinov for their assistance in reviewing the manuscript.

## Additional information Conflict of interest

The authors have declared that no competing interests exist.

## Ethical statement

No ethical statement was reported.

## Funding

Financial support for this project to RDT was provided by NSF (IOS 2319711), start-up funding from University of California Berkeley, and grants from the Society of Systematic Biologists, North Carolina Herpetological Society, Society for the Study of Reptiles and Amphibians, Chicago Herpetological Society, Texas Herpetological Society, the EEB Program at University of Texas at Austin, National Science Foundation Graduate Research Fellowship Program Graduate Research Opportunities Worldwide, National Geographic Society (Young Explorer Grant #9468-14). Support was also provided by NSF to DCC and RDT (DEB 1556967).

## Author contributions

Funding acquisition: RDT, DCC. Conceptualization: MBC, RDT. Investigation: MBC, RDT, JCR. Methodology: MBC, RDT, JCR, AJC. Formal analysis: MBC. Writing – original draft: MBC, RDT. Writing – review and editing: MBC, RDT, DCC, JCR, AJC.

## Data availability

All of the data that support the findings of this study are available in the main text or supplementary material and were uploaded to public databases.

## Notes

### Competing Interest Statement

The authors have declared no competing interest.

### Summary of Updates

We made a few minor corrections to the text and revealed the new species name.

## REFERENCES

Abadía G (1973) La música folklórica colombiana. Universidad Nacional de Colombia, Dirección de Divulgación Cultural, Bogotá D.C.- Colombia, 158 pp.

Anganoy-Criollo M, Cepeda-Quilindo B (2017) Redescription of the tadpoles of *Epipedobates narinensis* and *E. boulengeri* (Anura: Dendrobatidae). Phyllomedusa 16(2): 155–182. 10.11606/issn.2316-9079.v16i2p155-182

Aristizabal M (2002) El festival del Currulao en Tumaco: Dinámicas culturales y construcción de identidad étnica en el litoral pacífico colombiano. Master Thesis, Universidad del Valle, Valle del Cauca, Colombia.

Barbour T (1909) Corrections regarding the names of two recently described Amphibia Salientia. Proceedings of the Biological Society of Washington 22: 89.

Bioacoustics Research Program (2014) Raven Pro: Interactive Sound Analysis Software (Version 1.5) [Computer software] (1.5). The Cornell Lab of Ornithology.

Birenbaum Quintero M (2006) “La música pacífica” al Pacífico violento: Música, multiculturalismo y marginalización en el Pacífico negro colombiano. Trans. Revista Transcultural de Música 10: 1–32. https://www.redalyc.org/pdf/822/82201002.pdf

Birenbaum Quintero M (2019) Rites, Rights & Rhythms A Genealogy of Musical Meaning in Colombia’s Black Pacific. Oxford University Press, Canada, 342 pp. https://www.google.com/books/edition/Rites_Rights_and_Rhythms/YtB0DwAAQBAJ

Boulenger GA (1899) Descriptions of new reptiles and batrachians collected by Mr. P. O. Simons in the Andes of Ecuador. Annals and Magazine of Natural History Series 7, 4: 454–457.

Brown J, Twomey E, Amézquita A, Barbosa De Souza M, Caldwell J, Lötters S, Von May R, Melo-Sampaio P, Mejía-Vargas D, Perez-Peña P, Pepper M, Poelman E, Sanchez-Rodriguez M, & Summers K (2011) A taxonomic revision of the Neotropical poison frog genus *Ranitomeya* (Amphibia: Dendrobatidae). Zootaxa 120(3083): 1–120. 10.11646/zootaxa.3083.1.1

Caldeira Costa R, Gomes Facure K, Giaretta A (2006) Courtship, vocalization, and tadpole description of *Epipedobates flavopictus* (Anura: Dendrobatidae) in southern Goiás, Brazil. Biota Neotropica 6(1). 10.1590/S1676-06032006000100006

Chao K-H, Barton K, Palmer S, Lanfear R (2021) sangeranalyseR: Simple and Interactive Processing of Sanger Sequencing Data in R. Genome Biology and Evolution 13(3): 1–7. 10.1093/gbe/evab028

Cisneros-Heredia D, Yánez-Muñoz M (2010) A new poison frog of the genus *Epipedobates* (Dendrobatoidea: Dendrobatidae) from the north-western Andes of Ecuador. Avances 2(3): B83–86. http://www.cisneros-heredia.org/pdfs/2010_Epipedobatesdarwinwallacei.pdf

Clough M, Summers K (2000) Phylogenetic systematics and biogeography of the poison frogs: evidence from mitochondrial DNA sequences. Biological Journal of the Linnean Society 70(3): 515–540. 10.1006/bijl.1999.0418

Coloma L (1995) Ecuadorian Frogs of the Genus *Colostethus* (Anura: Dendrobatidae). Miscellaneous Publication. Museum of Natural History, University of Kansas 87: 1–72. 10.5962/bhl.title.16171

Cruz Hoyos S (2016) “Samuelito”, el hombre que hizo del currulao una danza universal. https://www.Elpais.Com.Co/Entretenimiento/Cultura/Samuelito-El-Hombre-Que-Hizo-Del-Currulao-Una-Danza-Universal.Html.

Edgar RC (2004) MUSCLE: Multiple sequence alignment with high accuracy and high throughput. Nucleic Acids Research 32(5): 1792–1797. 10.1093/nar/gkh340

Erdtmann L, Amézquita A (2009) Differential evolution of advertisement call traits in dart-poison frogs (Anura: Dendrobatidae). Ethology 115(9): 801–811. 10.1111/j.1439-0310.2009.01673.x

Funkhouser JW (1956) New frogs from Ecuador and southwestern Colombia. Zoologica. New York 41: 73–80.

Globally Unique Identifiers Task Group (2011) GUID and Life Sciences Identifiers Applicability Statements. Biodiversity Information Standards (TDWG). http://www.tdwg.org/standards/150

Goebel AM, Donnelly JM, Atz ME (1999) PCR Primers and Amplification Methods for 12S Ribosomal DNA, the Control Region, Cytochrome Oxidase I, and Cytochromebin Bufonids and other Frogs, and an overview of PCR Primers which have amplified DNA in amphibians auccessfully. Molecular Phylogenetics and Evolution 11(1): 163–199. 10.1006/MPEV.1998.0538

González-Santoro M, Hernández-Restrepo J, Palacios-Rodríguez P (2021) Aggressive behaviour, courtship and mating call description of the neotropical poison frog *Phyllobates aurotaenia* (Anura: Dendrobatidae). Herpetology Notes 14: 1145–1149. https://www.biotaxa.org/hn/article/view/60982

Grant T, Frost DR, Caldwell JP, Gagliardo R, Haddad CFB, Kok PJR, Means DB, Noonan BP, Schargel WE, Wheeler WC (2006) Phylogenetic systematics of dart-poison frogs and their relatives (Amphibia: Athesphatanura: Dendrobatidae). Bulletin of the American Museum of Natural History 299: 1–262. dx.doi.org/10.1206/0003-0090(2006)299[1:PSODFA]2.0.CO;2

Grant T, Rada M, Anganoy-Criollo M, Batista A, Dias PH, Jeckel AM, Machado DJ, Rueda-Almonacid JV (2017) Phylogenetic Systematics of Dart-Poison Frogs and Their Relatives Revisited (Anura: Dendrobatoidea). South American Journal of Herpetology 12(s1): S1–S90. 10.2994/SAJH-D-17-00017.1

Guevara Calderón A, Godoy Acosta C (2015) El Currulao: una propuesta de exploración, interpretación y creación, a través de la batería y la guitarra eléctrica. Master Thesis, Universidad del Bosque, Bogotá D.C. - Colombia.

Jungfer K-H (2017) On Warszewicz’s trail: The identity of *Hyla molitor* O. SCHMIDT, 1857. Salamandra 53(1): 18–24. https://www.researchgate.net/publication/316542564

Kalyaanamoorthy S, Minh BQ, Wong TKF, Von Haeseler A, Jermiin LS (2017) modelfinder: fast model selection for accurate phylogenetic estimates. 14(6). 10.1038/nmeth.4285

Kampstra P (2008) Beanplot: A Boxplot Alternative for Visual Comparison of Distributions. Journal of Statistical Software 28: 1–9. 10.18637/jss.v028.c01

Köhler J, Jansen M, Rodriguez A, Kok PJR, Toledo LF, Emmrich M, Glaw F, Haddad CFB, Rödel M-O, Vences M (2017) The use of bioacoustics in anuran taxonomy: Theory, terminology, methods and recommendations for best practice. Zootaxa 4251(1): 1–124. 10.11646/zootaxa.4251.1.1

Larsson A (2014) AliView: a fast and lightweight alignment viewer and editor for large datasets. Bioinformatics 30(22): 3276–3278. 10.1093/BIOINFORMATICS/BTU531

Lê S, Josse J, Husson F (2008) FactoMineR: An R Package for multivariate analysis. Journal of Statistical Software 25(1): 1–18. 10.18637/JSS.V025.I01

López-Hervas, K., Santos, J. C., Ron, S. R., Betancourth-Cundar, M., Cannatella, D. C., & Tarvin, R. D. (2024). Deep divergences among inconspicuously colored clades of *Epipedobates* poison frogs. Molecular Phylogenetics and Evolution 195: 108065. 10.1016/j.ympev.2024.108065

Lötters S, Reichle S, Jungfer K-H (2003) Advertisement calls of Neotropical poison frogs (Amphibia: Dendrobatidae) of the genera *Colostethus*, *Dendrobates* and *Epipedobates*, with notes on dendrobatid call classification. Journal of Natural History 37(15): 1899–1911. 10.1080/00222930110089157

Lötters S, Jungfer KH, Henkel FW, Schmidt W (2007) Poison Frogs. Biology, Species Captive Maintenance. Frankfurt am Main: Edition Chimaira.

Lynch JD, Suárez-Mayorga AM (2004) Catálogo de anfibios en el Chocó Biogeográfico. pp. 654–668. In: J. O. Rangel (ed.). Colombia Diversidad Biótica IV. El Chocó Biogeográfico. Universidad Nacional de Colombia, Bogotá, D.C.

Marques Correia da Rocha S, Pimentel Lima A, Kaefer IL (2018) Reproductive Behavior of the Amazonian Nurse-Frog *Allobates paleovarzensis* (Dendrobatoidea, Aromobatidae). South American Journal of Herpetology 13(3): 260–270. 10.2994/SAJH-D-17-00076.1

Minh BQ, Nguyen MAT, Von Haeseler A (2013) Ultrafast approximation for phylogenetic bootstrap. Molecular Biology and Evolution 30(5): 1188–1195. 10.1093/MOLBEV/MST024

Minh BQ, Schmidt HA, Chernomor O, Schrempf D, Woodhams MD, Von Haeseler A, Lanfear R, Teeling E (2020) IQ-TREE 2: New models and efficient methods for phylogenetic inference in the genomic era. Molecular Biology and Evolution 37(5): 1530–1534. 10.1093/MOLBEV/MSAA015

Moss JB, Tumulty JP, Fischer EK (2023) Evolution of acoustic signals associated with cooperative parental behavior in a poison frog. Proceedings of the National Academy of Sciences of the United States of America 120(17): 1–7. 10.1073/PNAS.2218956120/-/DCSUPPLEMENTAL

Mueses-Cisneros, J. J., Cepeda-Quilindo, B., & Moreno-Quintero, V. (2008). Una nueva especie de Epipedobates (Anura: Dendrobatidae) del suroccidente de Colombia. Papéis Avulsos de Zoologia (São Paulo) 48(1): 1–10. 10.1590/S0031-10492008000100001

Myers CW, Daly JW (1976) Preliminary evaluation of skin toxins and vocalizations in taxonomic and evolutionary studies of poison-dart frogs (Dendrobatidae). Bulletin of the American Museum of Natural History 157(3): 173–262.

Myers N, Mittermeier RA, Mittermeier CG, Da Fonseca GAB, Kent J (2000) Biodiversity hotspots for conservation priorities. Nature 403: 853–858. 10.1038/35002501

Noble GK (1921) Five new species of Salientia from South America. American Museum Novitates 29: 1–7. http://hdl.handle.net/2246/4615

Paradis E, Schliep K (2019) ape 5.0: an environment for modern phylogenetics and evolutionary analyses in R. Bioinformatics 35(3): 526–528. 10.1093/BIOINFORMATICS/BTY633

Peters WCH (1873) Über eine neue Schildrötenart, Cinosternon Effeldtii und einige andere neue oder weniger bekannte Amphibien. Monatsberichte der Königlichen Preussische Akademie des Wissenschaften zu Berlin 603–618.

Phillips N (2017) yarrr: a companion to the e-book “YaRrr!: the pirates guide to R.

Powers RP, Jetz W (2019) Global habitat loss and extinction risk of terrestrial vertebrates under future land-use-change scenarios. Nature Climate Change 9(4): 323–329. 10.1038/s41558-019-0406-z

R Core Team (2023) R: A Language and Environment for Statistical Computing. R Foundation for Statistical Computing. https://www.R-project.org/.

Santos JC, Baquero M, Barrio-Amorós C, Coloma LA, Erdtmann LK, Lima AP, Cannatella DC (2014). Aposematism increases acoustic diversification and speciation in poison frogs. Proceedings of the Royal Society B 281: 20141761. 10.1098/rspb.2014.1761

Santos JC, Cannatella DC (2011) Phenotypic integration emerges from aposematism and scale in poison frogs. Proceedings of the National Academy of Sciences of the United States of America 108(15): 6175–6180. 10.1073/pnas.1010952108

Santos JC, Coloma LA, Cannatella DC (2003) Multiple, recurring origins of aposematism and diet specialization in poison frogs. Proceedings of the National Academy of Sciences of the United States of America 100(22): 12792–12797. 10.1073/pnas.2133521100

Santos JC, Coloma LA, Summers K, Caldwell JP, Ree R, Cannatella DC (2009) Amazonian amphibian diversity is primarily derived from late Miocene Andean lineages. PLoS Biology 7 e1000056(3): 0448–0461. 10.1371/journal.pbio.1000056

Silverstone PA (1976) A revision of the poison-arrow frogs of the genus Phyllobates Bibron in Sagra Natural History Museum of Los Angeles County Science Bulletin 27.

Sueur J, Aubin T, Simonis C (2008) Seewave, a free modular tool for sound analysis and synthesis. Bioacoustics 18: 213–226. http://cran.r-project.org/src/contrib/Descriptions/

Tarvin RD, Powell EA, Santos JC, Ron SR, Cannatella DC (2017) The birth of aposematism: High phenotypic divergence and low genetic diversity in a young clade of poison frogs. Molecular Phylogenetics and Evolution 109: 283–295. 10.1016/j.ympev.2016.12.035

Vargas-S F, Bolaños-L ME (1999) Anfibios y reptiles en hábitats perturbados de selva lluviosa Tropical en el Bajo Anchicayá, Pacífico Colombiano. Revista Academia Ciencias Exactas, Físicas y Naturales 23 (Suplemento Especial): 499–511.

Venables W, Ripley B (2002) Modern Applied Statistics with S (Fourth Edition). Springer. New York, XII, 498 pp. https://www.stats.ox.ac.uk/pub/MASS4/.

Vences M, Kosuch J, Boistel R, Haddad CFB, La Marca E, Lötters S, Veith M (2003) Convergent evolution of aposematic coloration in Neotropical poison frogs: a molecular phylogenetic perspective. Organisms Diversity & Evolution 3: 215–226. http://www.urbanfischer.de/journals/ode

Warren R, Vanderwal J, Price J, Welbergen JA, Atkinson I, Ramirez-Villegas J, Osborn TJ, Jarvis A, Shoo LP, Williams SE, Lowe J (2013) Quantifying the benefit of early climate change mitigation in avoiding biodiversity loss. Nature Climate Change 3(7): 678–682. 10.1038/nclimate1887

Watters JL, Cummings ST, Flanagan RL, Siler CD (2016) Review of morphometric measurements used in anuran species descriptions and recommendations for a standardized approach. Zootaxa 4072(4): 477–495. 10.11646/zootaxa.4072.4.6

