## Supplementary material for "Honoring the Afro-Colombian musical culture with the naming of *Epipedobates currulao* sp. nov. (Anura, Dendrobatidae), a frog from the Pacific rainforests": S7_Results_PCA_LDA.docx

Supplementary material 7: Results of the Principal Component Analysis and Linear Discriminant Analysis for advertisement call characteristics of *E. currulao* sp. nov. and other species of the genus.

Results of variance explained by each component in the PCA.

| Principal Component | Eigenvalue | Variance (%) | Cumulative Variance (%) |
| --- | --- | --- | --- |
| PC1 | 3.14748299 | 44.9640427 | 44.96404 |
| PC2 | 2.30803443 | 32.9719205 | 77.93596 |
| PC3 | 0.72402832 | 10.3432617 | 88.27922 |
| PC4 | 0.52921898 | 7.5602711 | 95.8395 |
| PC5 | 0.22668337 | 3.2383338 | 99.07783 |
| PC6 | 0.0338963 | 0.4842328 | 99.56206 |
| PC7 | 0.03065562 | 0.4379374 | 100 |

Results of correlation between advertisement calls traits and PCs

| **Call trait** | **PC1** | **PC2** | **PC3** | **PC4** | **PC5** |
| --- | --- | --- | --- | --- | --- |
| Call Duration (s) | 0.8568665 | 0.4004343 | 0.1361066 | 0.26293974 | 0.01937671 |
| Number of Pulses | 0.8395691 | 0.3681253 | 0.1116902 | 0.36111733 | -0.04890828 |
| Intercall Interval (s) | 0.6607398 | 0.2815869 | 0.4578115 | -0.51844854 | -0.07480869 |
| Low Frequency (Hz) | -0.146105 | 0.9258641 | -0.1622284 | -0.10385429 | 0.27785258 |
| High Frequency (Hz) | -0.7411362 | 0.5561055 | 0.1401215 | 0.10850789 | -0.32255493 |
| Peak Frequency (Hz) | -0.508342 | 0.8494198 | -0.0143045 | -0.01779896 | -0.00682724 |
| 90% Bandwidth (Hz) | -0.6654084 | -0.2118671 | 0.6612719 | 0.19496302 | 0.19242972 |

Coefficients of linear discriminants in the LDA

| **PC** | **LD1** | **LD2** | **LD3** |
| --- | --- | --- | --- |
| PC1 | 1.7152917 | 0.422672 | 0.00769169 |
| PC2 | 0.6461926 | -1.47006052 | 0.05081234 |
| PC3 | 0.3081834 | -0.163502 | -0.02211733 |
| PC4 | 0.7684822 | 0.03986448 | 0.1620413 |
| PC5 | 0.3880722 | -0.49436924 | -2.18789095 |
| PC6 | 0.2842764 | 0.29889094 | -1.29748641 |
| PC7 | -0.5724359 | 1.6277654 | -1.52118054 |

Proportion of trace:

LD1: 0.6593

LD2: 0.3228

LD3: 0.0179
