## Supplementary material for "Honoring the Afro-Colombian musical culture with the naming of *Epipedobates currulao* sp. nov. (Anura, Dendrobatidae), a frog from the Pacific rainforests": S8_traduccion_2024-07-25.docx

Está traducción se hizo inicialmente por Google Translate en el 25 de julio del 2024. Será corregida para la versión final del manuscrito.

**Honrando la cultura musical afrocolombiana con el nombre de *Epipedobates currulao* sp. nov. (Anura: Dendrobatidae), una rana de las selvas tropicales del Pacífico.**

Mileidy Betancourth-Cundar^1,2^, Juan Camilo Ríos-Orjuela^1,3^, Andrew J. Crawford^1,4^, David C. Cannatella^5^, Rebecca D. Tarvin^6^

*^1^ Departamento de Ciencias Biológicas, Universidad de los Andes, Bogotá, 111711, Colombia*

*^2^ Department of Biology, Stanford University, Palo Alto, CA, 94305, USA*

*^3^ Grupo de Morfología y Ecología Evolutiva, Instituto de Ciencias Naturales, Universidad Nacional de Colombia, Sede Bogotá, 7495, Colombia*

*^4^ Museo de Historia Natural C.J. Marinkelle, Universidad de los Andes, Bogotá, 111711, Colombia*

*^5^ Department of Integrative Biology and Biodiversity Center, University of Texas, Austin, TX 78712, USA*

*^6^ Museum of Vertebrate Zoology and Department of Integrative Biology, University of California, Berkeley, Berkeley, CA 94720, USA*

**Resumen**

El número de especies de anfibios descritas cada año continúa aumentando, especialmente en las regiones tropicales, lo que implica que la biodiversidad de anfibios sigue siendo subestimada. Describimos una nueva especie de rana venenosa de las tierras bajas del Pacífico del suroccidente de Colombia: *Epipedobates currulao* sp. nov., nombrada así por el género de música y danza del Pacífico conocido como *bambuco viejo* o *currulao*. Las ranas de esta especie habitan en bosques de tierras bajas desde el nivel del mar hasta los 260 m. Este taxón se diferencia de sus congéneres por tener una combinación de manchas amarillas brillantes en la región dorsal anterior del muslo y los brazos, una coloración dorsal homogénea marrón oscuro y cantos de advertencia más largos y en consecuencia con mayor número de pulsos. También describimos la llamada de cortejo de *E. currulao*, con menor frecuencia pico y duración que la llamada de advertencia. Los análisis filogenéticos confirman la monofilia de la especie y su posición como hermana de *Epipedobates narinensis*, la cual se distribuye en el suroccidente de Colombia. Entre las especies de *Epipedobates*, la nueva especie ha sido previamente asignada a *E. boulengeri*, pero las dos especies son alopátricas y representan dos clados filogenéticamente divergentes (1.77% divergentes para 12S–16S y 5.39% para CYTB). Estas especies se pueden distinguir fenotípicamente por la presencia de una mancha amarilla brillante en la región dorsal anterior del muslo y en la parte superior del brazo en E. currulao, que son más blancas que amarillas o están ausentes en E. boulengeri. Además, los cantos de advertencia son distintos, *E. currulao* tiene una única y larga llamada en una serie de llamadas, mientras que E. boulengeri tiene de 2 a 6 llamadas por serie, siendo cada llamada mucho más corta. *Epipedobates* currulao es la especie distribuida más al norte del género *Epipedobates*, el cual se extiende hacia el sur a lo largo del flanco occidental de la cordillera de los Andes. Esta región conocida como el Chocó biogeográfico, ha sido fuertemente transformada por agricultura en Ecuador y está experimentando una transformación generalizada de sus bosques en Colombia, lo cual pone en peligro a *E. currulao* y toda su biodiversidad en un futuro cercano.

**Palabras clave**: bioacústica, Chocó, música, megadiverso, ranas venenosas, especie nueva, taxonomía alfa, códigos de barras de AND

**INTRODUCCIÓN**

*Epipedobates* es un taxón de ranas venenosas neotropicales (Anura: Dendrobatidae) con nueve especies putativas (Grant et al. (2006, 2017) y López-Hervas et al. (2024)): *Epipedobates* aff. *espinosai* (ver López-Hervas et al. 2024), *E. anthonyi* (Noble 1921), *E. boulengeri* (Barbour 1909), *E.* sp. 1 (ver López-Hervas et al. 2024), *E. espinosai* (Funkhouser 1956), *E. machalilla* (Coloma 1995), *E. maculatus* (Peters 1873), *E. narinensis* (Mueses-Cisneros et al. 2008) y *E. tricolor* (Boulenger 1899). Las ranas del género *Epipedobates* habitan en bosques tropicales secos y húmedos desde el nivel del mar hasta los 1800 m en las tierras bajas y estribaciones del lado occidental de los Andes de Colombia, Ecuador y el norte de Perú (Grant et al. 2006, 2017). Una evaluación reciente de la diversidad genética y fenotípica en *Epipedobates* boulengeri encontró varias líneas evolutivas distintas con fenotipos similares, es decir, especies crípticas (López-Hervas et al. 2024). Por lo tanto, no es sorprendente que estudios filogenéticos anteriores que muestrearon diferentes subconjuntos de linajes que ahora sabemos corresponden a diferentes especies no lograran resolver algunas relaciones filogenéticas en *Epipedobates* (Clough & Summers 2000; Grant et al. 2017; Santos et al. 2014; Tarvin et al. 2017; Vences et al. 2003).

Estudios previos de *Epipedobates boulengeri* demostraron alta diversidad genética y variación interpoblacional en rasgos acústicos y morfología larval, indicando que los linajes asignados a *Epipedobates boulengeri* probablemente representaban un complejo de especies (López-Hervas et al. 2024; Lötters et al. 2003; Santos et al. 2009; Tarvin et al. 2017). La evaluación filogenética más reciente (López-Hervas et al. 2024) concluyó que *Epipedobates boulengeri* contenía representantes de cuatro linajes genéticamente distintos: *E. boulengeri* (*sensu stricto*) distribuido en el suroeste de Colombia y el borde noroeste de Ecuador; *E. espinosai* distribuido en el centro-noroeste de Ecuador; *E.* sp. 1 distribuido en el suroeste de Colombia; y *E.* aff. *espinosai* distribuido en el noroeste de Ecuador y hacia el extremo sur de las estribaciones andinas de Colombia (López-Hervas et al. 2024). La evidencia fenotípica apoya esta diversidad críptica; por ejemplo, Lötters et al. (2003) encontraron diferencias en el número de notas y la longitud de la llamada de advertencia entre especímenes referidos como *E. boulengeri* de Colombia (Anchicayá, Valle del Cauca, correspondiente a E. sp. 1) y *E. boulengeri* de Ecuador (Lita, Imbabura, correspondiente a E. aff. espinosai). Además, Anganoy-Criollo y Cepeda-Quilindo (2017) describieron la morfología externa de los renacuajos de *E. boulengeri* de Colombia (Tumaco, Nariño, posiblemente representando a *E. boulengeri* sensu stricto) y de Ecuador (provincias de Carchi, Esmeraldas, Imbabura y Santo Domingo, identificación incierta, posiblemente correspondiendo a *E*. aff. *espinosai*, *E. boulengeri* sensu stricto o *E. espinosai*). Aunque solo examinaron renacuajos de la mitad sur del área de distribución de *E. boulengeri* (es decir, excluyendo el linaje descrito como *E.* sp. 1 por López-Hervas et al. 2024), encontraron que las poblaciones ecuatorianas diferían de las de Colombia en la forma del cuerpo, las puntas de las papilas marginales y el patrón de coloración de la cola. Así, la evidencia previa apuntaba fuertemente a *E. boulengeri* como un complejo de especies.

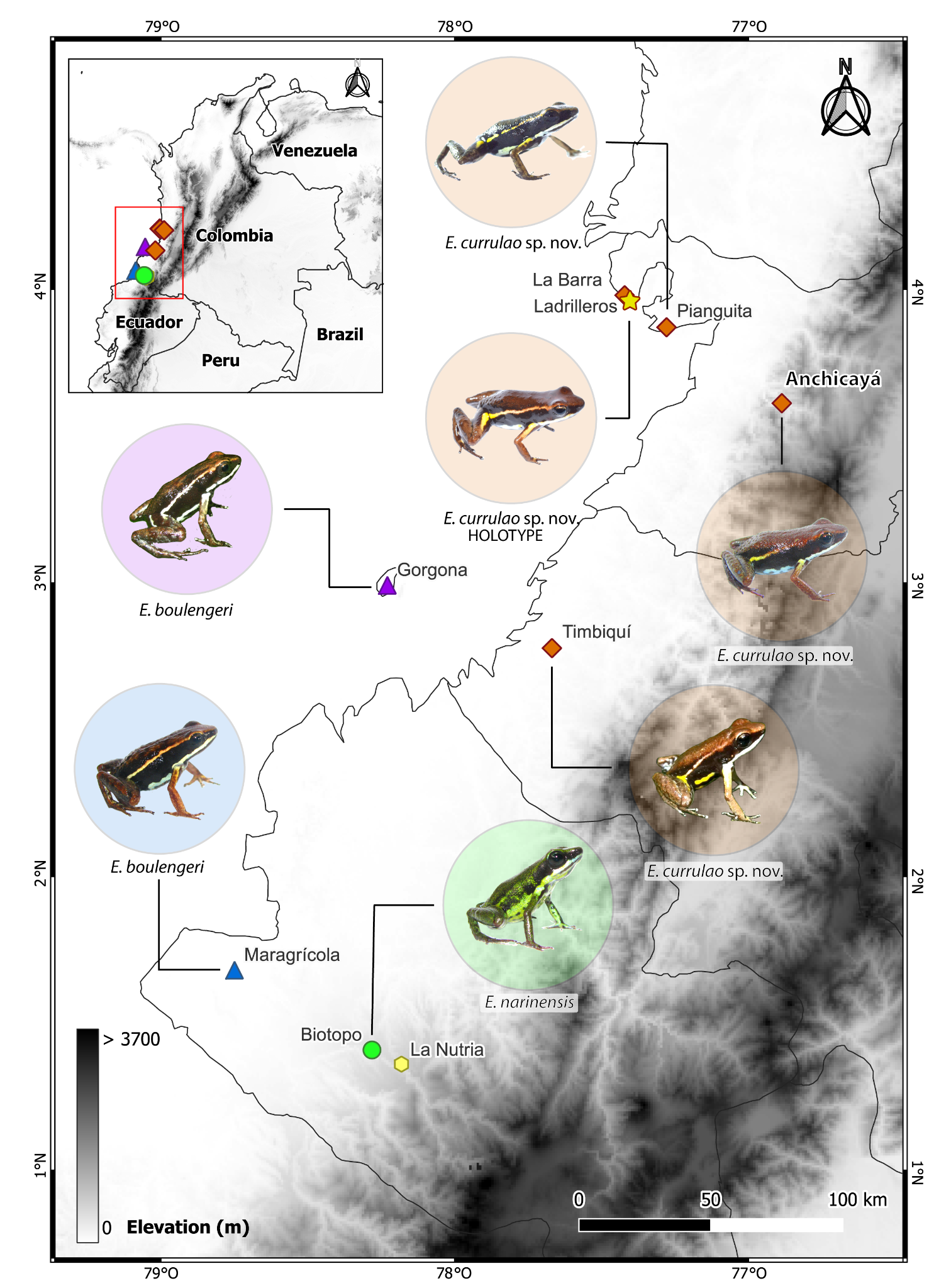

**Figura 1. Mapa del área de investigación**. La mitad sur de las tierras bajas del Pacífico de Colombia, parte de la región biogeográfica del Chocó, alberga al menos tres especies de *Epipedobates*. La fotografía de la rana de Timbiquí (Fundación ProAves y la Reserva Ranita Terribilis) se obtuvo de iNaturalist (observación No. 135253843); otras imágenes fueron tomadas por los autores.

En este estudio, describimos una nueva especie de *Epipedobates* de las tierras bajas del Pacífico colombiano correspondiente a *E.* sp. 1 según lo asignado por López-Hervas et al. (2024). Nuestros análisis acústicos, morfológicos y moleculares son consistentes con el trabajo de López-Hervas et al. (2024), que identificó a esta rana como una especie distinta y previamente no reconocida. Nombramos a la nueva especie *Epipedobates currulao* sp. nov. y proporcionamos una descripción detallada de sus llamadas, morfología, comportamiento e historia natural. Como muchas otras especies de anfibios (Myers et al. 2000; Powers & Jetz 2019; Warren et al. 2013), *Epipedobates currulao* sp. nov. está amenazada por la pérdida de hábitat, la contaminación y el cambio climático. Al identificar y describir esta nueva especie, podemos comprender mejor la diversidad, evolución, distribución geográfica, requisitos de hábitat y necesidades de conservación de las especies de *Epipedobates* en Colombia. Nuestros hallazgos estimularán más investigaciones y esfuerzos de conservación para proteger la biodiversidad anfibia única e irremplazable de la región biogeográfica del Chocó.

**MÉTODOS**

**Declaración de Ética**

Los procedimientos de recolección científica con animales vivos siguieron protocolos aprobados por el IACUC de la Universidad de Texas en Austin (AUC-2012-00032), la Universidad de California, Berkeley (AUP-2019-08-12457) y el CICUAL de la Universidad de los Andes (POE 18–003). La investigación y recolección de muestras en campo fueron autorizadas por la Autoridad Nacional de Licencias Ambientales (ANLA) de Colombia bajo el permiso marco resolución No. 1177 otorgado a la Universidad de los Andes. Se recolectaron especímenes de *E. boulengeri* de la localidad tipo bajo Res. 061-2016 de Parques Nacionales Naturales de Colombia. Las muestras fueron exportadas bajo los siguientes permisos: CITES No. CO39282, CO41443, CO46948 y ANLA No. 00561.

**Colecta científica**

Recolectamos datos genéticos, morfológicos y acústicos, así como especímenes completos, de diferentes especies y poblaciones de *Epipedobates* en Colombia (Fig. 1; ver también López-Hervas et al., 2024) durante 2014, 2016 y 2022. Animales fueron sacrificados por una sobredosis de lidocaína. El sexo de la mayoría de las muestras se determinó en el momento de la recolección en función de si los individuos llamaban o no y se confirmó mediante examen directo de las gónadas después de la eutanasia. Antes de la fijación, se tomaron muestras de hígado y músculo para análisis genéticos moleculares y se almacenaron en etanol al 95%. En algunos casos, se eliminó la piel para cuantificar el contenido de alcaloides defensivos (ver Suppl. material 1). Los especímenes voucher se fijaron en formalina tamponada neutra al 10% y se transfirieron directamente a etanol al 70% para su almacenamiento a largo plazo en la colección de anfibios del Museo de Historia Natural C.J. Marinkelle de la Universidad de los Andes en Bogotá, Colombia (ANDES:A) y la Colección de Herpetología del Museo de Zoología de Vertebrados de la Universidad de California, Berkeley en Berkeley, California, EE. UU. (MVZ:Herp).

**Morfología**

Siguiendo descripciones previas de nuevas especies de *Epipedobates* (Cisneros-Heredia & Yánez-Muñoz, 2010; Mueses-Cisneros et al., 2008), tomamos 13 medidas morfométricas estandarizadas después de la fijación (Watters et al., 2016): Hocico-respiradero-longitud (SVL); longitud del antebrazo (FAL), tomada desde la punta del codo flexionado y el borde proximal del tubérculo palmar; longitud de la mano (HaL), medida desde el borde proximal del tubérculo palmar medial grande hasta la punta del dedo más largo (cuarto); longitud de la tibia (TL), tomada desde la superficie exterior de la rodilla flexionada hasta la flexión del talón; longitud del pie (FL), tomada desde el borde proximal del tubérculo metatarsiano externo hasta el final del cuarto dedo; ancho de la cabeza (HW), tomado entre los ángulos de las mandíbulas; longitud de la cabeza (HL), distancia desde la punta del hocico hasta el ángulo de la mandíbula; distancia desde el centro de una de las fosas nasales hasta el borde anterior del ojo (NED); ancho de los ojos (EW); distancia ojo-nares (FIN); distancia interna (IND); distancia interocular (IOD); y diámetro horizontal del tímpano (TD). Todas las mediciones se tomaron bajo un microscopio de disección utilizando un calibrador digital con una resolución de 0,01 mm. Realizamos una prueba de Kruskal-Wallis para comparar el tamaño corporal (hocico-respiradero-longitud) entre especies y una comparación pareada post-hoc utilizando una prueba de suma de rangos de Wilcoxon. También realizamos un ANOVA para evaluar el dimorfismo sexual en el tamaño corporal, comprobando previamente sus supuestos.

También revisamos y resumimos la variación fenotípica a nivel de especie como se describe en López-Hervas et al. (2024) para proporcionar una evaluación a nivel de especie de la variación del patrón de color que pueda usarse para distinguir visualmente las especies.

**Análisis moleculares y filogenéticos.**

El ADN se extrajo del tejido hepático y muscular utilizando el kit Qiagen DNeasy Blood & Tissue (Valencia, CA) siguiendo el protocolo del fabricante. Secuenciamos dos fragmentos de genes mitocondriales de 5 individuos de *Epipedobates currulao* sp. nov. recolectado de la localidad tipo: un fragmento que incluye partes de los genes de ARNr mitocondrial *12S* y *16S* y el gen intermedio del ARNt de valina (*12S-16S*; 692 pb) y el citocromo b (*CYTB*; 659 pb). Utilizamos los mismos cebadores y protocolos descritos en López-Hervas et al. (2024). La región *12S-16S* se secuenció utilizando el cebador directo 12Sa (5'-AACTGGGATTAGATACCCCACTAT-3') y el cebador inverso 16SH-H (5'-TACCTTTTGCATCATGGTCTAGC-3') con condiciones de PCR: 2 min a 94 °C para la desnaturalización inicial seguida por 35 ciclos de 30 s a 94°C, 30 s a 46.5°C y 1 min a 72°C y un tiempo de extensión final de 7 min a 72°C (Santos y Cannatella, 2011). La región *CYTB* se secuenció utilizando el cebador directo CytbDen3-L (5'-AAYATYTCCRYATGATGRAAYTTYGG-3') y el cebador inverso CytbDen1-H (5'-GCRAANAGRAAGTATCATTCNGGYT-3') utilizando las mismas condiciones de PCR descritas *para 12S-16S*. Los productos de PCR se secuenciaron en ambas direcciones en el Laboratorio de Genómica Funcional QB3 de la Universidad de California, Berkeley (RRID:SCR_022170). Los cromatogramas se recortaron y se obtuvieron cóntigos de consenso utilizando sangeranalyseR (Chao et al., 2021) en R v.4.3.1 (R Core Team, 2023). Se depositaron nuevas secuencias en GenBank (números de acceso OR789875 – OR789884).

Alineamos las cinco nuevas secuencias usando MUSCLE v3.8.31 (Edgar, 2004) en AliView (Larsson, 2014) con datos existentes de *12S–16S* y *CYTB* para otros 108 individuos de Epipedobates, incluido un grupo externo (*Silverstoneia nubicola*) y 4 individuos asignados a *Epipedobates* sp. 1 de López-Hervas et al. (2024), que corresponden aquí a *Epipedobates currulao* sp. nov. Utilizando la alineación resultante (Suppl. material 2), calculamos las distancias p medias por pares (no corregidas) entre especies para cada gen con el script R proporcionado por López-Hervas et al. (2024) con la función 'dist.dna' en ape v5.7.1 (Paradis & Schliep, 2019), con el modelo configurado como "raw" y pairwise.deletion establecida en TRUE. Siguiendo a López-Hervas et al. (2024), excluimos a los individuos del grupo *E*. aff. *espinosai* localidades de Bilsa y La Tortuga, las cuales presentan evidencia de introgresión con *E. machalilla.*

Complementamos nuestro conjunto de datos moleculares con tres marcadores nucleares (longitud total de 2227 pb), histona H3 (*H3*), proteína morfogenética ósea 2 (BMP2) y canal de potasio 1.3 activado por voltaje (*K_V_1.3*), así como la región de control del ADNmt (*CR*; 1031 pb) de secuencias publicadas previamente de los mismos 108 individuos mencionados anteriormente (Suppl. material 1). También agregamos 28 secuencias de otros 23 individuos de *Epipedobates* que se publicaron previamente (Grant et al. 2006, 2017; Santos & Cannatella, 2011; Santos et al., 2003, 2009) a la alineación (ver Suppl. material 1 para los números de GenBank). Estas secuencias primero se recortaron en las regiones genéticas presentes en la alineación y luego se alinearon usando MUSCLE en AliView. Se estimó una filogenia de máxima verosimilitud (ML, por sus siglas en inglés) utilizando esta alineación ampliada en IQ-TREE v2.3.3 (Kalyaanamoorthy et al., 2017; Minh et al., 2020) y los parámetros descritos en López-Hervas et al. (2024). Brevemente, dividimos los genes codificadores de proteínas por gen y por posición de codón. Excluimos la primera y segunda posición de los codones de *BMP* y *H3*, que fueron invariantes en todas las muestras. Seleccionamos el modelo de evolución y particiones de datos bajo el Criterio de Información Bayesiano usando la opción -m TESTMERGE, parámetros de frecuencia base que incluyen iguales y estimados (-mfreq F, FO) y parámetros de heterogeneidad de tasas que incluyen iguales, gamma e invariantes (-mrate E , G, I). Realizamos el análisis tres veces y verificamos la coherencia entre las ejecuciones. Para evaluar el soporte de las ramas, realizamos 10 000 réplicas de arranque ultrarrápido (Minh et al., 2013) y las trazamos en el árbol de probabilidad óptima (Suppl. material 2).

**Grabación de llamadas y análisis bioacústicos.**

Registramos los llamados publicitarios de doce ranas macho de *Epipedobates* *currulao* sp. nov. de tres localidades del Valle del Cauca en condiciones naturales: Anchicayá (cuatro individuos), Pianguita (seis) y de la localidad tipo, Ladrilleros (dos). También registramos cinco machos de *E. boulengeri* de la localidad tipo (Isla Gorgona, Cauca, Colombia), cinco machos de *E. boulengeri* de Maragrícola (Tumaco, Nariño, Colombia) y seis machos de *E. narinensis* de su localidad tipo (Biotopo Reserva Natural, Nariño, Colombia). Todas las grabaciones fueron depositadas en Fonozoo (FZ-SOUND-CODE 14657-14685). Las grabaciones se realizaron durante la mañana (7 a 13 h) utilizando un micrófono unidireccional (Sennheiser K6/ME66) conectado a una grabadora digital (Marantz PMD660/Zoom H4n Pro) y colocado entre 50 y 150 cm delante de un hombre que llamaba. Inmediatamente después del registro, medimos la temperatura del sustrato utilizando un termómetro infrarrojo (Oakton modelo 35629). Digitalizamos las grabaciones con una resolución mínima de 16 bits y una frecuencia de muestreo de 44,1 kHz en RAVEN PRO 1.6 (Programa de Investigación Bioacústica, 2014). Para la terminología y los procedimientos para medir los rasgos de llamada seguimos a Köhler et al. (2017). Los espectrogramas y oscilogramas se graficaron utilizando el paquete Seewave R (Sueur et al., 2008) con una ventana FFT utilizando el algoritmo de Blackman, una longitud de ventana de 256 muestras y una superposición del 90%.

Definimos una llamada como el sonido producido durante una única contracción muscular abdominal (Erdtmann & Amézquita, 2009; Köhler et al., 2017). En *Epipedobates*, tales contracciones producen un único "zumbido" compuesto por varios pulsos (Brown et al., 2011; Erdtmann & Amézquita, 2009; Myers & Daly, 1976). Analizamos cinco llamadas consecutivas por individuo midiendo su frecuencia máxima (la frecuencia con mayor amplitud), baja frecuencia (la frecuencia más baja de la llamada), alta frecuencia (la frecuencia superior de la llamada) y 90% de ancho de banda (-10 dB). umbral que contiene el rango de frecuencia que abarca el 90% de la energía sonora en cada llamada). Las frecuencias altas y bajas de las llamadas se midieron a 20 dB (re 20 mPA) por debajo de la intensidad máxima de la frecuencia máxima. Con este valor todavía se podía distinguir claramente la energía de la señal del ruido de fondo. También medimos las siguientes variables temporales: duración de la llamada, número de pulsos por llamada e intervalo entre llamadas. Para *E. currulao* sp. nov., también medimos la frecuencia del pulso, la duración del pulso y el intervalo entre pulsos. Los parámetros de llamada para las llamadas de cada individuo se promediaron para comparar estadísticamente entre individuos. Para evitar la redundancia entre las variables acústicas, utilizamos un Análisis de Componentes Principales (PCA) para reducir la cantidad de parámetros implementados en la función 'PCA' del paquete R 'FactoMineR' (Lê et al., 2008). Con el fin de identificar si *E. currulao* sp. nov. podrían diferenciarse en función de las llamadas publicitarias, ejecutamos un LDA en las primeras 5 PC usando la función lda' del paquete MASS R (Venables & Ripley, 2002). Utilizamos una prueba de Kruskal-Wallis para examinar los rasgos de llamada relevantes (frecuencia máxima, duración de la llamada y número de pulsos por llamada) que diferencian a *E. currulao* sp. nov. de otras especies y una comparación pareada post-hoc utilizando una prueba de suma de rangos de Wilcoxon. Representamos visualmente estos rasgos utilizando el paquete R “yarrr” de la función “pirateplot” (Phillips, 2017). Los gráficos piratas muestran todos los datos brutos, su distribución, la media y los intervalos de mayor densidad (IDH) del 95% de la media de cada estimación (Kampstra, 2008). Utilizamos 999 iteraciones para calcular el IDH.

**Repositorios, siglas institucionales y abreviaturas institucionales**

A lo largo de nuestro artículo utilizamos identificadores únicos globalmente (GUID, Globally Unique Identifiers Task Group, 2011) para referirnos a los especímenes de muestra en QCAZ (QCAZ:A:XXXX, Museo de Zoología de la Pontificia Universidad Católica del Ecuador, Quito, Ecuador), la MVZ (MVZ:Herp:XXXX, colecciones herpetológicas del Museo de Zoología de Vertebrados, Universidad de California, Berkeley, EE. UU.), y la colección de anfibios del Museo de Historia Natural C.J. Marinkelle de la Universidad de los Andes, Bogotá, Colombia (ANDES:A:XXXX), en un esfuerzo por facilitar la futura legibilidad mecánica de este texto. Las abreviaturas de las series de campo son las siguientes: RDT (Rebecca D. Tarvin), AJC (Andrew J. Crawford), MBC (Mileidy Betancourth-Cundar).

**RESULTADOS**

**Taxonomía**

**Nombre**

*Epipedobates currulao* sp. nov.

*Epipedobates boulengeri*: Silverstone (1976), Lötters et al. (2003), Vargas-S & Bolaños-L (1999), Castro-Herrera & Vargas-Salinas (2008), Lynch & Suárez-Mayorga (2004), Lötters et al. (2007)

*Epipedobates* sp. 1: López-Hervas et al. (2024)

Nombre común en inglés propuesto: Currulao Nurse Frog

Nombre común español propuesto: Rana nodriza de currulao

<http://zoobank.org/> [el número de registro se agregará una vez que se acepte el documento]

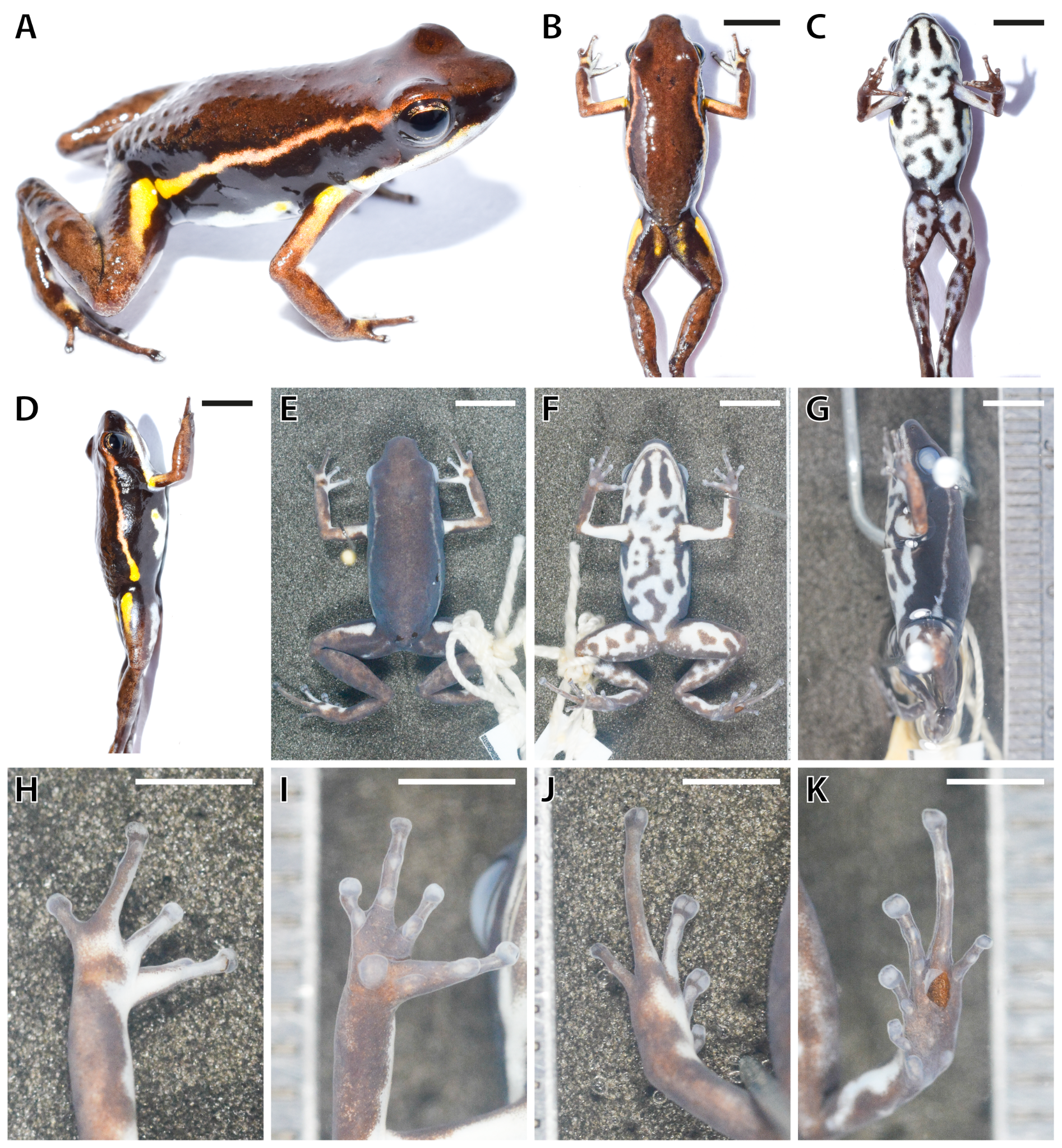

**Figura 2**. Imágenes en vida y en conservante del holotipo de Epipedobates currulao sp. nov. Un espécimen completo en la vida; B Vista dorsal en vida; C Vista ventral en la vida; D Vista lateral de la vida; E Vista dorsal en conservante (70% etanol); F Vista ventral en conservante; G Vista lateral en conservante; H Mano dorsal en conservante; I mano ventral en conservante; J Pie dorsal en conservante; K Pie ventral en conservante. Las barras de escala representan 5 mm (B a G) o 2,5 mm (H a K).

**Material**

**Holotipo**

COLOMBIA • ♀; Ladrilleros, Buenaventura, Valle del Cauca; 3.945221, −77.364993; 28 msnm; 6 Aug. 2022; Rebecca D. Tarvin, Mileidy Betancourth-Cundar, Juan Camilo Ríos-Orjuela leg.; ANDES:A:5255.

**Topoparatipos**

COLOMBIA • 4 ♀♀, 6 ♂♂, 1 ND; mismos datos que para el holotipo; ANDES:A:5254, 5256–5265 • 3 ♀♀, 2 ♂♂; mismos datos que para el holotipo; Genbank: OR789880–84 and OR789875–79; MVZ:Herp:295432–295436. 1 ND; Ladrilleros, Buenaventura, Valle del Cauca; 3.950882, −77.358293; 53 msnm; 26 Nov. 2014; Rebecca D. Tarvin y Fray Arriaga leg.; Genbank: OR489012, OR179791, OR734704, OR179836, and OR179880; ANDES:A:2464.

**Otros materiales**

**COLOMBIA**

• *Epipedobates currulao* 2 ♀♀, 5 ND; Localidad tipa, Ladrilleros, Buenaventura, Valle del Cauca; 3.950882, −77.358293; 53 msnm; 26 Nov. 2014; Rebecca D. Tarvin y Fray Arriaga leg.; GenBank: OR489011, OR179790, OR734703, OR179835, OR179875; ANDES:A:2458–2463, 2465. La piel se eliminó antes de la preservación para evaluar el contenido de alcaloides de estos individuos. • *E. currulao* 1 ♀; La Barra, Buenaventura, Valle del Cauca; 3.985064, –77.376723; 15 msnm; 26 Nov. 2014; Rebecca D. Tarvin y Fray Arriaga leg.; GenBank: OR489010, OR179789, OR734702, OR179834, OR179851; ANDES:A:2455. • *E. currulao* 5 ♂♂; Pianguita, Buenaventura, Valle del Cauca; 3.841954, –77.198718; 17 msnm; 12 Sep. 2016; Rebecca D. Tarvin, Mileidy Betancourth-Cundar, Sandra V. Flechas leg.; ANDES:A:3690–94. La piel se eliminó antes de la preservación para evaluar el contenido de alcaloides de estos individuos con la excepción de ANDES:A:3691. • *E. currulao* 5 ♀♀, 3 ♂♂, Danubio, Dagua, Valle del Cauca; 3.611528, –76.885194; 705 msnm; 5 Nov. 2016; Mileidy Betancourth-Cundar, Adolfo Amézquita, y Ivan Beltrán leg.; GenBank: OR489013, OR179792, OR734705, OR179837; ANDES:A:3713–20. La piel se eliminó antes de la preservación para evaluar el contenido de alcaloides de estos individuos.

• *E. boulengeri* 3 ♀♀, 4 ♂♂, 1 ND; Isla Gorgona, Guapi, Cauca; 2.96465, –78.173685; 20 msnm; 12 Sep. 2016; Rebecca D. Tarvin, Mileidy Betancourth-Cundar y Sandra V. Flechas leg.; GenBank: OR OR488992, OR179771, OR734681, OR179812; ANDES:A:3695-3702. • 1 ND; Isla Gorgona, Guapi, Cauca; 08 Sep. 2005; ANDES:A:0560; Colector desconocido. • *E. boulengeri* 3 ♀♀, 4 ♂♂, 1 ND; Isla Gorgona, Guapi, Cauca; 2.96465, –78.173685; 20 msnm; 12 Sep. 2016; Rebecca D. Tarvin, Mileidy Betancourth-Cundar, Sandra V. Flechas leg.; GenBank: OR488992, OR179771, OR734681, OR179812; ANDES:A:3695-3702. • *E. boulengeri* 8 ♀♀; Maragrícola, Tumaco, Nariño; 1.68084, –78.74924; 7 msnm; 09 Dec. 2014; Rebecca D. Tarvin, Mileidy Betancourth-Cundar, Cristian Flórez leg.; GenBank: OR488990, OR179769, OR734679, OR179810, OR179862, OR488991, OR179770, OR734680, OR179811, OR179863; ANDES:A:2468–75. • *E. boulengeri* 1 ND; La Nutria, El Diviso, Barbacoas, Nariño; 1.36083, –78.18076; 762 msnm; 10 Dec. 2014; Rebecca D. Tarvin, Mileidy Betancourth-Cundar, Cristian Flórez leg.; GenBank: OR488977, OR179756, OR734663, OR179794, OR179840; ANDES:A:2476.

• *E. narinensis* 3 ♀♀, 5 ♂♂, 1 ND; Reserva Natural Biotopo, Berlín, Barbacoas, Nariño; 1.408999, –78.281246; 518 msnm; 25 Sep. 2016; Rebecca D. Tarvin, Mileidy Betancourth-Cundar y Cristian Flórez leg.; GenBank: OR489008, OR179787, OR734700, OR179832, OR489009, OR179788, OR734701, OR179833; ANDES:A:3703–3711. • *E. narinensis* 10 ♂♂; Reserva Natural Biotopo, Berlín, Barbacoas, Nariño; 1.411263, –78.285099; 600 msnm; 22 Jul. 2006; Viviana Moreno-Quintero, Jonh Jairo Mueses-Cisneros, Luisa Mercedes Bravo, Carol Narváez, y Bienvenido Cortés leg.; ICN-A:53344 (holotype), ICN-A:53336–53340, 53342–53346 (paratipos).

• *Andinobates minutus* 4 ND; La Barra, Buenaventura, Valle del Cauca; 3.985064, –77.376723; 15 msnm; 25 Nov. 2014; Rebecca D. Tarvin and Fray Arriaga leg.; ANDES:A 2451–54. • *A. minutus* 1 ND; Ladrilleros, Buenaventura, Valle del Cauca; 3.945221, −77.364993; 28 msnm; 6 Aug. 2022; Rebecca D. Tarvin, Mileidy Betancourth-Cundar, Juan Camilo Ríos-Orjuela leg.; ANDES:A:5266.

**ECUADOR**

• *Epipedobates* aff. *espinosai* 1 ND; Lita, Carchi; 12 Aug. 1992; M. Bueno leg.; ICN-A:32504.

**Diagnóstico**

*Epipedobates currulao* es una pequeña rana dendrobátida (SVL media = 17,99 mm y SD = 0,95 mm, N = 16 ranas; Tablas 1, 2) con coloración dorsal uniformemente marrón, lados negros, una franja lateral oblicua de blanco a amarillo, una mancha amarilla brillante en el lado anterodorsal del muslo y en la parte superior del brazo, y un vientre de color azul pálido o turquesa con moteado negro (Fig. 2, Suppl. material 3). Llamadas de *E. currulao* sp. nov son largos con una duración de llamada de 0,67 a 3,88 s (media = 2,21, DE = 0,54 s, N = 15) y 22 a 122 pulsos por llamada (media = 73,98, DE = 18,77, N = 15). Ocurren en series de llamadas de una sola llamada (Tablas 3, 4).

**Tabla 1**. Caracteres morfométricos y estadísticas resumidas de los machos de la serie tipo de *Epipedobates currulao* sp. nov. Medidas (en mm) de hembras y machos adultos; los valores se dan como media, DE y rango. Consulte la sección Métodos para obtener una explicación de las abreviaturas. *Holotipo.

| Código del museo | Código del colector | Sexo | SVL | FAL | HaL | TL | FL | HW | HL | NED | EW | END | IND | IOD | TD |
| --- | --- | --- | --- | --- | --- | --- | --- | --- | --- | --- | --- | --- | --- | --- | --- |
| ANDES:A:5254 | MBC753 | Macho | 17.20 | 4.30 | 3.70 | 7.70 | 7.00 | 5.4 | 6.3 | 2.5 | 2.10 | 1.60 | 2.10 | 2.90 | 0.90 |
| ANDES:A:5256 | AJC07731 | Macho | 17.20 | 4.10 | 3.90 | 7.80 | 6.90 | 5.80 | 6.90 | 2.90 | 2.10 | 1.60 | 2.20 | 3.30 | 1.00 |
| ANDES:A:5260 | AJC07735 | Macho | 17.40 | 4.40 | 3.70 | 7.90 | 7.00 | 5.60 | 6.40 | 3.00 | 2.50 | 1.60 | 2.50 | 3.80 | 1.00 |
| ANDES:A:5261 | AJC07736 | Macho | 18.70 | 4.50 | 4.30 | 8.50 | 7.30 | 5.90 | 7.00 | 3.30 | 2.70 | 1.70 | 2.50 | 3.80 | 1.00 |
| ANDES:A:5263 | AJC07738 | Macho | 17.50 | 4.60 | 3.80 | 7.70 | 6.60 | 6.00 | 7.30 | 3.00 | 2.40 | 1.60 | 2.30 | 3.50 | 0.90 |
| MVZ:Herp:295434 | AJC07742 | Macho | 16.70 | 4.00 | 3.60 | 7.50 | 6.70 | 4.90 | 6.70 | 2.80 | 1.90 | 1.70 | 2.30 | 3.20 | 0.90 |
| MVZ:Herp:295436 | AJC07744 | Macho | 16.70 | 3.80 | 3.70 | 7.10 | 6.70 | 5.70 | 6.90 | 2.70 | 2.40 | 1.70 | 2.10 | 3.40 | 1.00 |
| ANDES:A:5259 | RDT0935 | Macho | 16.90 | 4.40 | 3.80 | 7.90 | 7.20 | 5.30 | 6.60 | 3.60 | 2.40 | 1.80 | 2.30 | 3.80 | 0.90 |
| Promedio | | | 17.29 | 4.26 | 3.81 | 7.76 | 6.93 | 5.58 | 6.76 | 2.98 | 2.31 | 1.66 | 2.29 | 3.46 | 0.95 |
| DE | | | 0.64 | 0.27 | 0.22 | 0.40 | 0.25 | 0.36 | 0.33 | 0.35 | 0.26 | 0.07 | 0.16 | 0.33 | 0.05 |
| Min | | | 16.70 | 3.80 | 3.60 | 7.10 | 6.60 | 4.90 | 6.30 | 2.50 | 1.90 | 1.60 | 2.00 | 2.90 | 0.90 |
| Max | | | 18.70 | 4.60 | 4.50 | 8.70 | 7.60 | 6.20 | 7.80 | 3.60 | 2.70 | 2.00 | 2.50 | 4.20 | 1.20 |

**Tabla 2**. Caracteres morfométricos y estadísticas resumidas de los hembras de la serie tipo de *Epipedobates currulao* sp. nov. Medidas (en mm) de hembras y machos adultos; los valores se dan como media, DE y rango. Consulte la sección Métodos para obtener una explicación de las abreviaturas. *Holotipo.

| Código del museum | Código del colector | Sexo | SVL | FAL | HaL | TL | FL | HW | HL | NED | EW | END | IND | IOD | TD |
| --- | --- | --- | --- | --- | --- | --- | --- | --- | --- | --- | --- | --- | --- | --- | --- |
| ANDES:A:5255* | MBC754 | Hembra | 18.50 | 4.60 | 3.90 | 8.10 | 6.90 | 5.70 | 6.80 | 3.00 | 2.30 | 1.70 | 2.30 | 3.70 | 0.90 |
| ANDES:A:5257 | AJC07733 | Hembra | 19.50 | 4.50 | 4.50 | 8.70 | 7.20 | 6.00 | 6.70 | 3.20 | 2.60 | 2.00 | 2.40 | 4.20 | 1.10 |
| ANDES:A:5262 | AJC07737 | Hembra | 18.30 | 4.40 | 3.90 | 7.70 | 6.70 | 6.10 | 7.00 | 3.20 | 2.40 | 1.80 | 2.00 | 3.60 | 1.10 |
| ANDES:A:5264 | AJC07739 | Hembra | 17.50 | 4.40 | 3.90 | 8.20 | 7.20 | 5.70 | 7.00 | 2.90 | 2.40 | 1.90 | 2.40 | 3.60 | 0.90 |
| MVZ:Herp:295433 | AJC07740 | Hembra | 19.10 | 4.30 | 4.00 | 8.00 | 7.20 | 6.20 | 7.10 | 3.10 | 2.30 | 1.90 | 2.40 | 3.40 | 1.20 |
| MVZ:Herp:295435 | AJC07743 | Hembra | 18.90 | 4.60 | 4.20 | 8.60 | 7.60 | 5.80 | 7.10 | 2.90 | 2.20 | 1.70 | 2.10 | 3.40 | 1.10 |
| MVZ:Herp:295432 | RDT0934 | Hembra | 19.10 | 4.30 | 4.00 | 8.00 | 7.00 | 5.90 | 7.80 | 2.90 | 2.40 | 1.60 | 2.40 | 3.50 | 0.90 |
| ANDES:A:5258 | AJC07734 | ND | 18.60 | 4.40 | 3.60 | 7.30 | 6.60 | 6.00 | 6.50 | 3.40 | 2.30 | 1.90 | 2.30 | 3.90 | 1.00 |
| Promedio | | | 18.70 | 4.44 | 4.06 | 8.19 | 7.11 | 5.91 | 7.07 | 3.03 | 2.37 | 1.80 | 2.29 | 3.63 | 1.03 |
| DE | | | 0.66 | 0.13 | 0.22 | 0.35 | 0.29 | 0.20 | 0.35 | 0.14 | 0.13 | 0.14 | 0.17 | 0.28 | 0.13 |
| Min | | | 17.50 | 4.30 | 3.90 | 7.70 | 6.70 | 5.70 | 6.70 | 2.90 | 2.20 | 1.60 | 2.00 | 3.40 | 0.90 |
| Max | | | 19.50 | 4.60 | 4.50 | 8.70 | 7.60 | 6.20 | 7.80 | 3.20 | 2.60 | 2.00 | 2.40 | 4.20 | 1.20 |

**Comparación de especies**

En la localidad tipo, la nueva especie ocurre en simpatría con *Andinobates minutus* (Dendrobatidae: Dendrobatinae) pero difiere en el tamaño corporal y la coloración (Fig. 3). *Andinobates minutus* tiene manchas anaranjadas en lugar de amarillas en el lado anterodorsal del muslo y el lado dorsal de los brazos y su banda supralabial es de color naranja oscuro en lugar de blanco como en *E. currulao*. Ventralmente en *A. minutus* la proporción de negro y azul turquesa es similar. En cambio, en *E. currulao* la coloración es principalmente azul turquesa con algunas manchas negras (Fig. 2C). Además, los adultos de *A. minutus* son mucho más pequeños (media = 12,00 mm, DE = 0,91 mm, N = 5) que los adultos de *E. currulao* (17,99 mm, DE = 0,95, N = 16), pero *A. minutus* puede confundirse con los juveniles de *E. currulao*. La coloración de los juveniles de *E. currulao* es muy similar a la de los adultos de *E. currulao*, por lo que aún se puede utilizar el color para diferenciar las dos especies.

Anteriormente, *E. currulao* se había confundido con *E. boulengeri*. Las dos especies se pueden diferenciar por el color (en vida) de la mancha en el lado anterodorsal del muslo y el color de la mancha en el lado dorsal del brazo cerca de la axila, ambas de color amarillo a naranja en *E. currulao* y de color blanco a amarillo blanquecino en *E. boulengeri.* Algunos individuos de *E. currulao* tienen coloración amarilla difusa a lo largo del borde lateral del vientre; esta coloración no ha sido identificada en ninguna población de *E. boulengeri* hasta la fecha. En comparación con la población de Isla Gorgona *E. boulengeri*, *E. currulao* tiene una franja lateral oblicua mucho más delgada; sin embargo, esta franja es similar en morfología a la población continental de *E. boulengeri*. La población continental de *E. boulengeri* (basada en imágenes de Maragrícola) se puede diferenciar de *E. currulao* por el tamaño, forma y color de la mancha en la cara anterodorsal del muslo, que es grande, amarilla o naranja, y claramente definido en *E. currulao* pero ausente o difuso y cobrizo transparente o blanquecino en *E. boulengeri* (Maragrícola). Además, *E. currulao* tiene llamadas publicitarias con una llamada por serie mientras que *E. boulengeri* tiene 2 a 3 llamadas (Isla Gorgona) o 3 a 6 llamadas por serie (Maragrícola) (Tabla 3).

Los individuos de *E. narinensis* y *E. currulao* se pueden diferenciar por el color dorsal, que es verde oliva en *E. narinensis* y marrón oscuro en *E. currulao*, la longitud de la línea lateral oblicua, que se extiende hasta el ojo en *E. currulao* pero solo se extiende hasta la mitad del cuerpo en *E. narinensis*, la forma de la mancha en el lado anterodorsal del muslo, que está claramente definida en *E. currulao* pero difusa en *E. narinensis*, y el color de fondo del vientre, que es de color amarillento a verdoso en *E. narinensis* y de azul pálido a turquesa o blanco en *E. currulao*. La estructura de la llamada publicitaria de la especie también difiere, donde las llamadas en *E. currulao* ocurren una por serie, pero de 5 a 14 por serie para *E. narinensis* (Tabla 3).

Individuos de *E.* aff. *espinosai* y *E. currulao* se pueden diferenciar por la longitud de los dedos III y V, que están reducidos en *E. currulao* pero no en *E*. aff. *espinosai* (ver Morfología en la sección Sistemática a continuación). La línea lateral oblicua se extiende con mayor frecuencia hasta el ojo en *E. currulao* pero tiende a terminar en la escápula en *E*. aff. *espinosai*. La forma de la mancha en el lado anterodorsal del muslo está claramente definida y es amarilla o naranja en *E. currulao* pero ausente o pequeña y de color azul blanquecino en *E.* aff. *espinosai*. La mancha en el lado dorsal del brazo cerca de la axila es de color amarillo brillante o amarillo blanquecino, mientras que la mancha está prácticamente ausente o tiene una coloración cobrizo o crema difusa en *E*. aff. *espinosai*.

Un resumen de los caracteres morfológicos de *E. currulao* sp. nov. en comparación con otras especies de Epipedobates en Colombia se muestra en la Tabla 3. Consulte Suppl. material 4 para medidas morfológicas de individuos y Suppl. Material 5 para notas más extensas sobre la variación de color dentro y entre especies.

**Tabla 3.** Características de morfología, coloración (en vida) y llamados de *E. currulao* sp. nov. en comparación con otras especies de *Epipedobates* encontradas en Colombia. Para la comparación de estructuras morfológicas se utilizaron especímenes depositados en el Instituto de Ciencias Naturales de la Universidad Nacional de Colombia. Para *E. narinensis* revisamos el holotipo (ICN-A:53344) y la serie tipo (ICN-A:53336–53340, 53342–53346). Para *E*. aff. *espinosai*, revisamos un espécimen disponible en el ICN (ICN-A:32504); López-Hervas et al. proporcionaron datos adicionales sobre el tamaño corporal. (2024). La tabla se presenta en partes a lo largo de las siguientes páginas para facilitar su lectura.

| **Especie** | **Longitud hocico-cloaca (mm)** (promedio ± DE) | **Longitud de los dedos II>III** | **Dedos III y V reducidos** | **Inflamación del dedo IV en hombres adultos** | **Correas: Dedo II Preaxial** | **Correas: Dedo III Preaxial** | **Correas: Dedo IV Preaxial** | **Correas: Dedo IV Postaxial** |
| --- | --- | --- | --- | --- | --- | --- | --- | --- |
| *E. currulao* | Hembras:  18,47 ± 0,91, N = 14  Machos: 16,98 ± 0,71, N = 16  Adultos: 18,09 ± 0,50, N = 8 | Sí | Sí | Sí | Ausente | Ausente | Presente, se extiende más allá del tubérculo subarticular basal del dedo IV | Ausente |
| *E. narinensis* | Hembras*:* 17,08 ± 0,77, N = 3  Machos: 16,82 ± 0,26, N = 5 | Sí | Sí | Sí | Presente, reducido | Ausente | Presente, se extiende hasta o justo debajo del tubérculo subarticular basal del dedo IV | Ausente |
| *E. boulengeri-* Gorgona | Hembras*: 20,13* ± 0,20, N = 3  Machos:  19,15 ± 0,75, N = 4  Adultos:  19,88 ± 1,81, N = 2 | Sí | No | Sí | Ausente | Ausente | Presente, se extiende más allá del tubérculo subarticular basal del dedo IV | Ausente |
| *E. boulengeri* - Nariño | Hembras: 17,57 ± 0,36, N = 8 | Sí | No | Sí | Ausente | Ausente | Presente, se extiende más allá del tubérculo subarticular basal del dedo IV | Ausente |
| *E.* aff. *espinosai* | Hembras: 17,68 ± 1,54, N = 11  Machos:  16,32 ± 1,63, N = 7  Adultos:  17,41 ± 1,58, N = 9 | Sí | No | Sí | Ausente | Ausente | Presente, se extiende más allá del tubérculo subarticular basal del dedo IV | Ausente |

| **Especie** | **Longitudes relativas de los dedos** | **Longitudes relativas de los dedos del pie** | **Pliegue metatarsiano** | **Textura de la piel dorsal** | **Color dorsal** | **Color de la raya lateral oblicua** | **Morfología de la raya lateral oblicua** | **Raya ventrolateral** |
| --- | --- | --- | --- | --- | --- | --- | --- | --- |
| *E. currulao* | IV >II>III>V | IV>III>V>II>I | Ausente | Finamente granulado | Coloración dorsal uniformemente marrón, sin marcas. | Presente; blanco, amarillo o amarillo anaranjado que se vuelve cobrizo cuando llega al ojo | Se extiende hasta el ojo o cerca del ojo; completo en la mayoría pero a veces interrumpido o difuso | Presente pero mal definido en algunos; blanco a blanco azulado |
| *E. narinensis* | IV >II>III>V | IV>III>V>II>I | Ausente | Finamente granulado | Verde oliva oscuro uniforme, sin marcas. | Presente; amarillo verde | Se extiende sólo hasta la mitad del cuerpo; difuso y mal definido | Presente pero indistinto en algunos; verde claro |
| *E. boulengeri-* Gorgona | IV >II>III>V | IV>III>V>II>I | Ausente | Finamente granulado | Marrón o marrón rojizo, con manchas oscuras. | Presente; amarillo cremoso | Se extiende hasta el ojo o cerca del ojo; completo y claramente definido | Presente y claramente definido en algunos; Blanco crema |
| *E. boulengeri-* Nariño | IV >II>III>V | IV>III>V>II>I | Ausente | Finamente granulado | Marrón rojizo con manchas oscuras. | Presente; naranja pálido dorado; más delgada que la población de Gorgona | Se extiende hasta el párpado superior; completo pero ligeramente difuso | Presente y claramente definido en algunos; blanco |
| *E.* aff. *espinosai* | IV >II>III>V | IV>III>V>II>I | Ausente | Finamente granulado | De color rojo ladrillo oscuro a marrón oscuro; algunos con manchas oscuras | Presente; blanco cerca de la ingle y luego se vuelve cobrizo cuando llega a la mitad del cuerpo | Se extiende hasta la escápula o el ojo; completo en algunos pero roto o difuso en otros | Presente y claramente definido en algunos; blanco |

| **Especie** | **Coloración de fondo ventral** | **Patrón ventral** | **Forma lateral del hocico** | **Forma dorsal del hocico** | **Tímpano/Ojo (%)** | **HW/SVL (%)** | **HW/HL (%)** | **Mancha en el lado anterodorsal del muslo** | **Color del brazo superior** |
| --- | --- | --- | --- | --- | --- | --- | --- | --- | --- |
| *E. currulao* | Varía del azul turquesa pálido al blanco; algunos con amarillo difuso en los lados | Generalmente muy moteado con manchas negras irregulares. | Saliente | Ligeramente redondeado | Hembras:  37,5 – 66,0  Machos:  33,3 – 55,3  Adultos:  43,1 – 61,6 | Hembras:  27,7 – 33,3  Machos:  29,1 – 36,3  Adultos:  27,9 – 32,8 | Hembras:  75,6 - 89,6  Machos:  73,1 – 103,7  Adultos:  68,4 – 92,3 | Presente y claramente definida: mancha de color amarillo a amarillo anaranjado. | Marcas de destellos de color amarillo brillante o amarillo blanquecino |
| *E. narinensis* | De color amarillento a verdoso, brillante en algunos pero en su mayoría opaco. | Moteado o ligeramente moteado de negro. | Saliente | Ligeramente redondeado | Hembras:  47,6 – 58,9  Machos: 55,3 – 58,1 | Hembras:  31,9 – 32,2 Machos:  29,8 – 33,7 | Hembras:  82,1 – 86,3  Machos:  79,3 – 87,6 | Coloración amarillo verdosa ausente o difusa. | En su mayoría ausentes, pero algunos con coloración amarilla difusa. |
| *E. boulengeri-* Gorgona | Azul turquesa pálido a casi blanco | Manchas negras irregulares y vermiculaciones. | Saliente | Ligeramente redondeado | Hembras:  50,0 – 57,1  Machos: 44,8 – 52,0  Adultos:  55,5 – 59,4 | Hembras: 28,7 – 32,0  Machos:  30,0 – 33,1  Adultos: 29,3 – 32,7 | Hembras:  90,0 – 103,2  Machos:  89,1 – 93,9  Adultos: 80,4 – 89,7 | Presente: amarillo-blanco; mucho menos amarillo que en E. currulao | En su mayoría ausentes, pero algunos con coloración amarilla blanquecina difusa; mucho menos amarillo que en *E. currulao* |
| *E. boulengeri-* Nariño | Blanco azulado | Moteado o con puntos negros aislados | Saliente | Ligeramente redondeado | Hembras:  45,0 – 64,0 | Hembras:  25,7 – 32,2 | Hembras:  76,4 – 86,6 | Ausente o menos común: amarillo blanquecino difuso (presente en 2/8 individuos) | En su mayoría ausentes, pero algunos con coloración crema difusa. |
| *E.* aff. *espinosai* | Azul turquesa pálido a casi blanco o crema | Manchas negras irregulares y vermiculaciones; rara vez sin moteado o con marcas amarillas | Saliente | Ligeramente redondeado | N / D | N / D | N / D | Ausente o menos común: blanco o azul blanquecino (presente en 8/28 individuos) | En su mayoría ausentes, pero algunos con coloración cobrizo o crema difusa |

| **Especie** | **Frecuencia máxima** (media ± DE, en kHz)  (Rango) | **Número de pulsos** (media ± DE)  (Rango) | **Duración de la llamada (s)** (media ± DE, en s)  (Rango) | **Número de llamadas en serie de llamadas**  **(Rango)** |
| --- | --- | --- | --- | --- |
| *E. currulao* (N = 15 ranas) | 5,23 ± 0,11  (4,98 – 5,47) | 73,94 ± 18,78  (22.0 – 122) | 2,21 ± 0,54  (0,67 – 3,88) | 1 |
| *E. narinensis* (N = 6) | 5,64 ± 0,21  (5.29 – 6.17) | 8,98 ± 2,45  (6,0 – 14,6) | 0,28 ± 0,070  (0,18 – 0,40) | 5-14 |
| *E. boulengeri -* Gorgona (N = 6) | 4,76 ±0,11  (4,56 – 4,90) | 14,48 ± 2,19  (9,5 – 20,0) | 0,30 ± 0,036  (0,21 – 0,37) | 2–3 |
| *E. boulengeri -* Nariño (N = 5) | 5,51 ± 0,11  (5,23 – 5,68) | 13,09 ± 1,75  (10,6 – 17,0) | 0,28 ± 0,042  (0,23 – 0,37) | 3–6 |
| *E.* aff. *espinosai* | N / D | N / D | N / D | N / D |

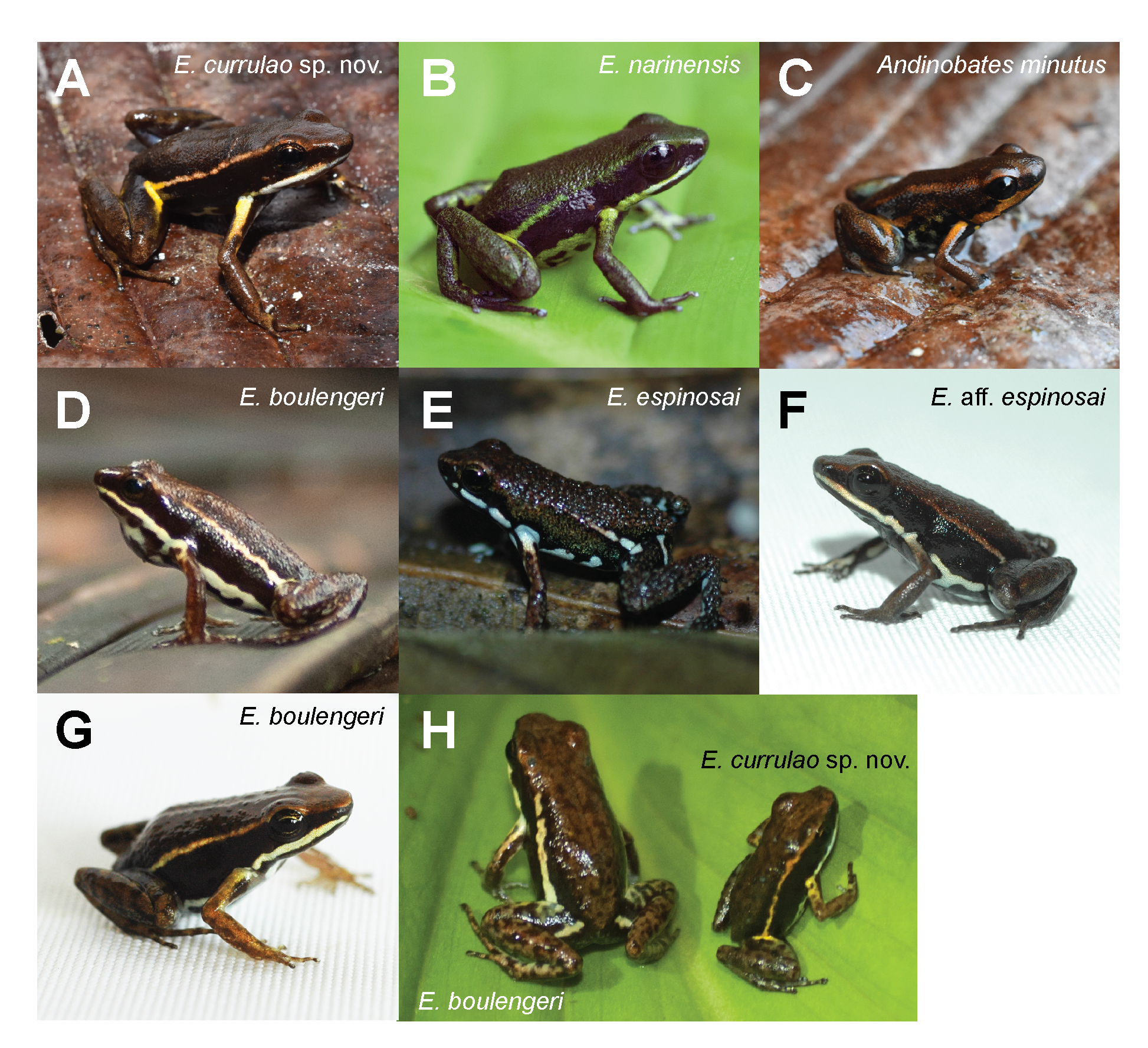

**Figura 3.** **Imágenes en vida de *Epipedobates currulao* sp. nov. en comparación con congéneres cercanos y especies simpátricas.** **A.** *E. currulao* sp. nov. de la localidad tipo de Ladrilleros, Valle de Cauca, Colombia (ANDES:A:5261; SVL = 20.0 mm; macho adulto; paratipo); **B.** *E. narinensis* de Biotopo, Nariño, Colombia (ANDES:A:3704; 16,39 mm; macho adulto); **C.** *Andinobates minutus* de Ladrilleros, Valle de Cauca, Colombia (ANDES:A:5266; 13 mm; sexo no determinado); **D.** *E. boulengeri* de Isla Gorgona, Cauca, Colombia (individuo no capturado); **MI.** *E. espinosai* de Río Palenque, Santo Domingo de los Tsáchilas, Ecuador (individuo no capturado); **F.** *E.* aff. *espinosai* de La Nutria, Nariño, Colombia (ANDES:A:2476; 17,77 mm; sexo no determinado); **G.** *E. boulengeri* de Maragrícola, Nariño, Colombia (ANDES:A:2472; 18,96 mm; hembra adulta); **H.** Una imagen de lado a lado de *E. boulengeri* de Isla Gorgona (ANDES:A:3695; 20,65 mm; hembra adulta) y *E. currulao* sp. nov. de Pianguita, Valle de Cauca, Colombia (ANDES:A:3690; 16,42 mm; hembra adulta) lo que demuestra la gran diferencia de tamaño entre las dos especies. Todas las imágenes fueron tomadas por RDT excepto A y C, que fueron tomadas por JCR. Las fotos no están a escala.

**Descripción**

**Coloración del holotipo en vida (Fig. 2A – D).** Superficies dorsales de color marrón oscuro con una franja lateral oblicua que se extiende desde la región posterior del ojo hasta la ingle, con una coloración amarilla anaranjada metálica en la región anterior que se vuelve amarilla hacia la ingle. Ingle de color negro parduzco oscuro con una clara mancha amarilla que continúa hasta la superficie anterodorsal del muslo. Flancos negros. Fondo de las extremidades anteriores y posteriores de color marrón oscuro con manchas irregulares de color marrón oscuro. Región anterior de la parte superior del brazo con una mancha amarilla de color similar a la mancha del muslo. Franja supralabial de color blanco cremoso que se extiende desde la narina hasta la axila. Superficies ventrales de color azul turquesa con manchas negras irregulares. Iris cobre. Dos puntos negros alargados en la región gular (Fig. 2C).

**Coloración del holotipo en conservante (Fig. 2E – K; después de dos años de conservación en etanol al 70%).** Dorsalmente negro a marrón oscuro, extremidades traseras marrón oscuro. Extremidades anteriores de color marrón más claro con algunas manchas de color marrón oscuro. Las manchas dorsales amarillas en vida en las extremidades anteriores y los muslos se vuelven blancas cuando se conservan. Ventralmente blanco pálido con manchas irregulares de color negro a marrón oscuro. Ingle negra. Línea lateral oblicua de color gris oscuro que se extiende desde la región posterior del ojo hasta la ingle. Franja supralabial de color gris claro que va desde la punta de la cara hasta la axila.

**Variación de coloración de series tipo y otras poblaciones en vida.** Todos los individuos de la serie tipo exhiben un color marrón oscuro uniforme en el dorso (Suppl. material 3). El dorso, la cabeza, el muslo y las patas presentan una textura de piel cubierta de tubérculos pequeños y dispersos. Flancos y superficies ocultas de las extremidades anteriores y posteriores lisas y de color negro sólido. Todos los individuos muestran una franja lateral oblicua que se extiende desde la región posterior del ojo hasta la ingle y difiere ligeramente entre poblaciones. En la población de Ladrilleros (localidad tipo), la franja lateral oblicua presenta un color amarillo anaranjado con bordes indistintos y se extiende desde la parte anterior de la ingle hasta el ojo y el canto rostral, cambiando gradualmente de un tono amarillo a un tono más anaranjado parduzco. En la población de Anchicayá, la franja lateral oblicua adquiere una tonalidad amarillenta y, en la mayoría de los ejemplares (6/8), se estrecha o se fragmenta en parches, desapareciendo gradualmente antes de llegar al ojo. En algunos individuos de Pianguita (3/5), la franja lateral oblicua presenta interrupciones regulares en toda su longitud.

La región de la ingle se caracteriza por un color marrón oscuro-negro, con una mancha amarilla notable que se extiende hasta la parte anterior interna del muslo (Suppl. material 3). La mayoría de los individuos tienen una mancha paracloacal de coloración similar, pero más pequeña y alargada, en la superficie dorsal posterior del muslo. En la población de Anchicayá estas manchas están presentes sólo en algunos individuos. La mayoría de las personas tienen una mancha amarilla en la región dorsal del brazo que coincide con el color de la mancha en el lado anterodorsal del muslo. Todos los individuos exhiben una franja labial superior de color blanco cremoso a azul turquesa pálido que es notablemente más clara que la franja lateral oblicua. En algunos casos, esta franja tiene una ligera cualidad iridiscente y se extiende desde debajo de la fosa nasal hasta la axila. Además, en ciertos individuos, continúa posteriormente como una franja ventrolateral vagamente definida. Las superficies ventrales de la garganta, el vientre y los muslos exhiben un color azul turquesa pálido a blanco con manchas negras irregulares y patrones que se asemejan a líneas parecidas a gusanos (vermiculaciones). En los machos, la coloración de fondo puede oscurecerse por un pigmento gris difuso ubicado justo por delante de la región pectoral y el saco vocal. Algunos individuos muestran una coloración amarillenta difusa hacia los bordes exteriores del vientre. Se pueden ver láminas de página completa de imágenes de cuatro poblaciones de *E. currulao* en las figuras complementarias 4 a 6 de López-Hervas et al. (2024) y las placas que muestran imágenes de las series tipo descritas en este documento se pueden encontrar en Suppl. material 3.

**Vocalizaciones.** Llamadas de *E. currulao* sp. nov consistió en 22 a 122 pulsos por llamada (media = 73,98, DE = 18,77, N = 15), con una duración de llamada de 0,67 a 3,88 s (media = 2,21, DE = 0,54 s, N = 15). La frecuencia del pulso constaba de 26 a 39 pulsos por segundo (media = 34,89, DE = 2,07, N = 15), duración del pulso de 0,009 a 0,028 (media = 0,017, DE = 0,004 s, N = 15) y intervalo entre pulsos de 0,001 a 0,068 s (media = 0,013, DE = 0,009 s, N = 15). El intervalo entre llamadas oscila entre 15,12 y 315,01 s (media = 62,67, DE = 60,76 s, N = 15) (Tabla 4). La frecuencia máxima osciló entre 4,98 y 5,46 kHz (media = 5,23, DE = 0,11 kHz), y la frecuencia ancho de banda intercuartil entre 0,21 y 0,67 kHz (media = 0,35, DE = 0,87 kHz) (Tabla 4). Las llamadas no están moduladas en frecuencia. La amplitud del primer y último pulso se reduce entre un 7 y un 10% en comparación con el resto de pulsos.

**Tabla 4.** Estadísticas resumidas de parámetros espectrales y temporales de las llamadas publicitarias de *Epipedobates currulao* sp. nov.

| **Parámetros** | **Duración de la llamada (s)** | **Número de pulsos** | **La frecuencia del pulso** | **Intervalo(s) de interllamada** | **Duración del pulso (s)** | **Intervalo(s) entre impulsos** | **Baja frecuencia (kHz)** | **Alta frecuencia (kHz)** | **Frecuencia pico (kHz)** | **Ancho de banda 90% (kHz)** |
| --- | --- | --- | --- | --- | --- | --- | --- | --- | --- | --- |
| **Promedio** | 2.213 | 73.938 | 34.894 | 62.675 | 0,017 | 0.013 | 4.845 | 5.586 | 5.237 | 0.354 |
| **DE** | 0.541 | 18.786 | 2.077 | 60.762 | 0.004 | 0.009 | 0.222 | 0.210 | 0,115 | 0,087 |
| **Min** | 0,674 | 22.000 | 26.000 | 15.120 | 0.009 | 0.001 | 4.236 | 5.221 | 4.981 | 0.215 |
| **Max** | 3.888 | 122.000 | 39.000 | 315.012 | 0,028 | 0.068 | 5.287 | 6.093 | 5.469 | 0,675 |

**Etimología**

El epíteto específico currulao es un sustantivo en aposición de género masculino. Se refiere al género musical que se originó en la costa del Pacífico sur de Colombia y Ecuador, donde se presenta *E. currulao* y también contribuye al paisaje sonoro local. El currulao, también conocido como bambuco viejo, es una práctica de sonido afrocolombiano que inspira el baile y transmite la alegría y tradición cultural de esta región. Es un símbolo de resiliencia frente a la opresión racial y regional (Abadía, 1973; Aristizabal, 2002; Birenbaum Quintero, 2006, 2019). Nombramos a esta especie en honor y homenaje a este género musical que representa la cultura del Pacífico sur colombiano porque: “la música, como la vida, no se pueden dejar perder” (Cruz Hoyos, 2016).

**Distribución**

*Epipedobates currulao* sp. nov. ocurre en el Departamento del Valle del Cauca en las tierras bajas del Pacífico del suroeste de Colombia. Estas ranas habitan en bosques de tierras bajas entre 0 y 260 m. La localidad tipo es Ladrilleros, Buenaventura, Valle del Cauca, Colombia. También observamos la especie en áreas cercanas a la localidad tipo incluyendo Corregimiento Pianguita y Corregimiento Juan Chaco (playa La Barra) en el municipio de Buenaventura. La distribución hacia el flanco occidental de la cordillera occidental es en la Vereda El Danubio, cuenca alta del río Anchicayá, Dagua, Valle del Cauca. Si asumimos que los rasgos de coloración de la nueva especie son consistentes para todas las poblaciones de esta especie, los registros de iNaturalist extenderían la distribución de *E. currulao* 194 km (132 km) en línea recta al sur hasta el municipio de Timbiquí, Cauca (ver iNaturalist observaciones N° 135253843 y 85214439 y Fig. 1). Debido a que los individuos de esta especie fueron asignados previamente a *E. boulengeri*, recomendamos una mayor exploración e inspección de especímenes de museo para comprender mejor la distribución geográfica de esta especie.

**Ecología**

*Epipedobates currulao* sp. nov. es una especie terrestre que se encuentra en el suelo durante el día en áreas agroforestales, en los bordes de bosques secundarios o en pequeños parches de bosque secundario perturbado siempre cerca o dentro de pantanos y/o arroyos de flujo lento (Fig. 4). Es probable que la especie también esté presente en los bosques primarios de la región. Observamos individuos moviéndose activamente entre la hierba y la hojarasca, o llamando activamente en los bordes de los cuerpos de agua. La localidad tipo (Ladrilleros, Valle de Cauca, Colombia) incluye pequeños fragmentos de bosque entre viviendas humanas. Por lo general, estas áreas están contaminadas con basura o residuos agrícolas (Fig. 4A-F). En Anchicayá, las poblaciones de *E. currulao* generalmente se encuentran a lo largo de caminos y bordes de bosques, siempre que haya pequeños arroyos y hojarasca (Fig. 4G-J). Observamos una mayor actividad vocal durante la mañana y al final de la tarde que durante las partes más cálidas o soleadas del día.

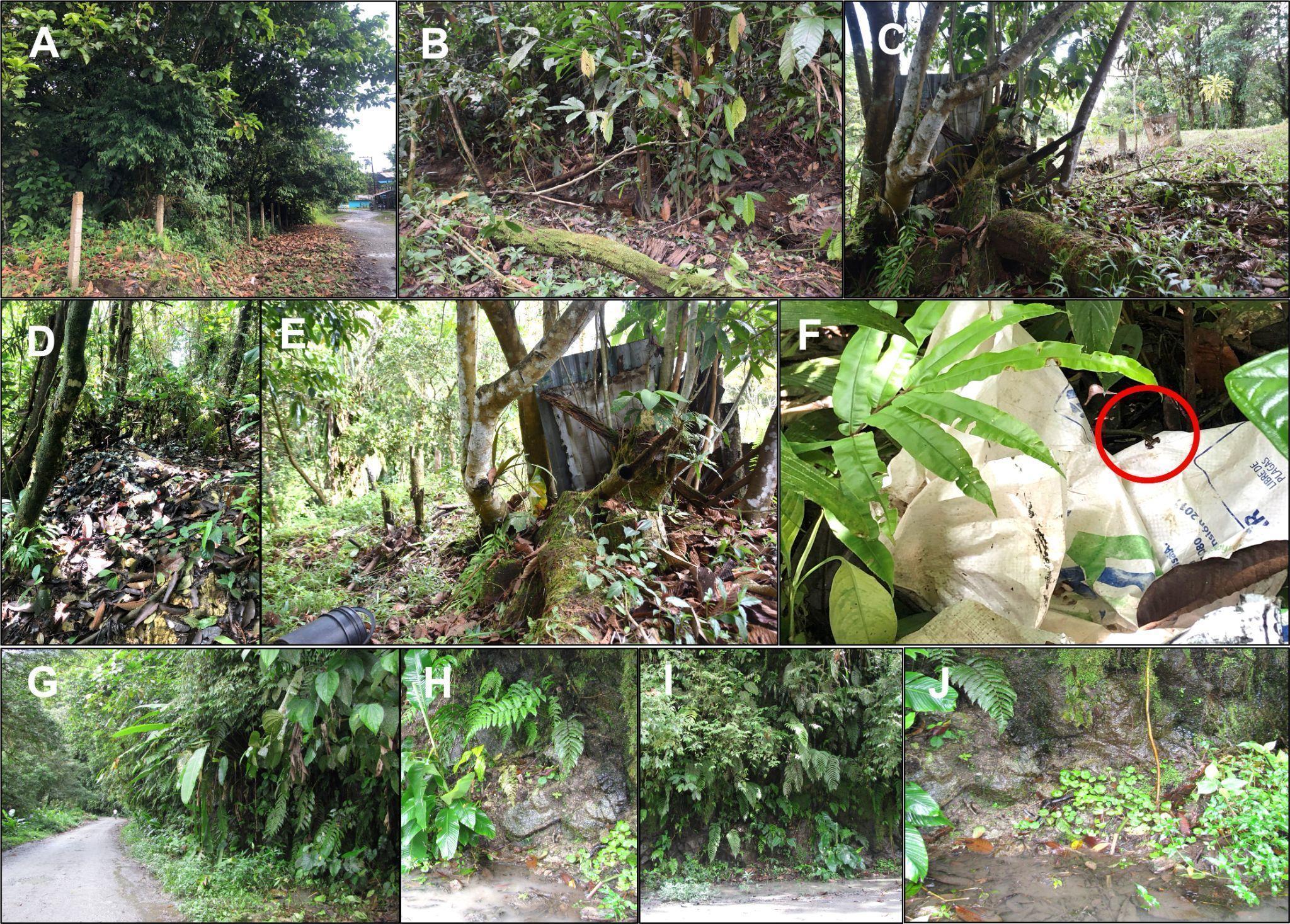

**Figura 4: Estructura del hábitat de *Epipedobates currulao* sp. nov. en dos localidades.** A–C Imágenes de la localidad tipo de Ladrilleros, Buenaventura, Valle del Cauca, Colombia. Por lo general, esta especie se encuentra en los bordes de las carreteras cerca de arroyos formados por las lluvias. D–F En la localidad tipo, observamos ranas en hábitats contaminados con basura o desechos agrícolas. Observe la rana en F (círculo rojo). G–J Características del hábitat en Anchicayá, Dagua, Valle del Cauca, Colombia. Las imágenes B, C y E fueron tomadas por RDT, todas las demás por MBC.

**Estado de conservación**

Las poblaciones observadas en las cuatro localidades (Ladrilleros, Pianguita, La Barra y Anchicayá) probablemente estén afectadas por la construcción de viviendas, basura o desechos agrícolas, así como por la fragmentación de bosques y la pérdida de hábitat. No sabemos si estas poblaciones se han adaptado a la perturbación humana o si son remanentes de la distribución original de la especie, pero los congéneres en Ecuador abundan en hábitats altamente modificados como las poblaciones de cacao y banano (aunque están notablemente ausentes en la palma africana). plantaciones; RDT obs.). Aunque consideramos que *E. currulao* sp. nov. es moderadamente abundante en la localidad tipo, ya que tres de nosotros (RDT, MBC, JCR) capturamos 17 individuos en un área aproximada de 30 m^2^ en cuatro horas, desconocemos su abundancia en hábitats menos perturbados. Mejorar nuestra comprensión de su estado de conservación requerirá monitoreo y exploraciones en localidades potenciales en Cauca y Valle del Cauca, especialmente en áreas protegidas cercanas a las localidades reportadas en este estudio.

A pesar de la falta de certeza sobre la distribución de *E. currulao* sp. de noviembre, es probable que su extensión sea inferior a 20.000 km^2^. Sabemos que la especie se encuentra en al menos cuatro localidades (Ladrilleros, Anchicayá, Pianguita y La Barra), quedando una más pendiente de validación genética (Timbiquí). Los bosques en el área de distribución de esta especie han estado, y probablemente seguirán estando, sujetos a fuertes presiones de deforestación que reducen la cantidad y calidad del hábitat disponible y aumentan su fragmentación. Se necesitarán más datos para determinar la presencia de *E. currulao* sp. nov. categorización, pero aquí ofrecemos algunas recomendaciones. Bajo la actitud precautoria, recomendaríamos categorizarlo como Vulnerable (VU: B1a, biii; UICN, 2019) con base en B1, una Extensión de ocurrencia (EOO) <20.000 km 2 (aprox. 3600 km 2 desde Timbiquí hasta Ladrilleros ) y (a) Severely fragmented OR Number of locations ≤ 10 and (b) Continuing decline observed, estimated, inferred or projected in any of: (iii) area, extent and/or quality of habitat. Under the evidentiary attitude, we would recommend que *E. currulao* sp. nov. ser categorizado como Casi Amenazado, reconociendo que más estudios pueden revelar poblaciones adicionales y ampliar su área de distribución y tamaño de población conocidos. Además, observamos esta especie dentro del Parque Nacional Natural Farallones de Cali, por lo que existe al menos una población conocida (Anchicayá) dentro de un área protegida. La distribución de *E. currulao* sp. nov. También está muy cerca del Parque Nacional Natural Uramba Bahía Málaga. No tenemos conocimiento de la presencia de *E. currulao* sp. nov. dentro del parque, pero esta área de conservación protege más de 47.000 ha de áreas marinas y costeras, por lo que es muy probable que *E. currulao* sp. nov. se encuentra dentro del parque. Más investigaciones sobre la distribución, los requisitos ecológicos y la dinámica poblacional de esta especie ayudarán a asignarla con seguridad a una categoría de amenaza.

**Sistemática**

**Análisis moleculares y filogenéticos**

Estimamos una filogenia utilizando datos existentes (López-Hervas et al., 2024) y 10 secuencias nuevas de cinco especímenes de *E. currulao* de la localidad tipo. La filogenia resultante contiene un fuerte apoyo a la monofilia de E. currulao y recupera relaciones filogenéticas conocidas entre las especies de este género (Fig. 5 y Suppl. material 6; López-Hervas et al., 2024). La especie hermana de *E. currulao* es *E. narinensis*, que se encuentra en la parte centro-sur de la costa del Pacífico de Colombia. Estas dos especies constituyen un clado de Epipedobates que es muy divergente y hermano de un clado que contiene todas las demás especies del género.

Las distancias p promedio entre clados de especies de Epipedobates muestran que E. currulao es genéticamente más similar a *E. narinensis* (1,77% para 12S-16S y 5,39% para CYTB), como se esperaba dada la filogenia molecular.

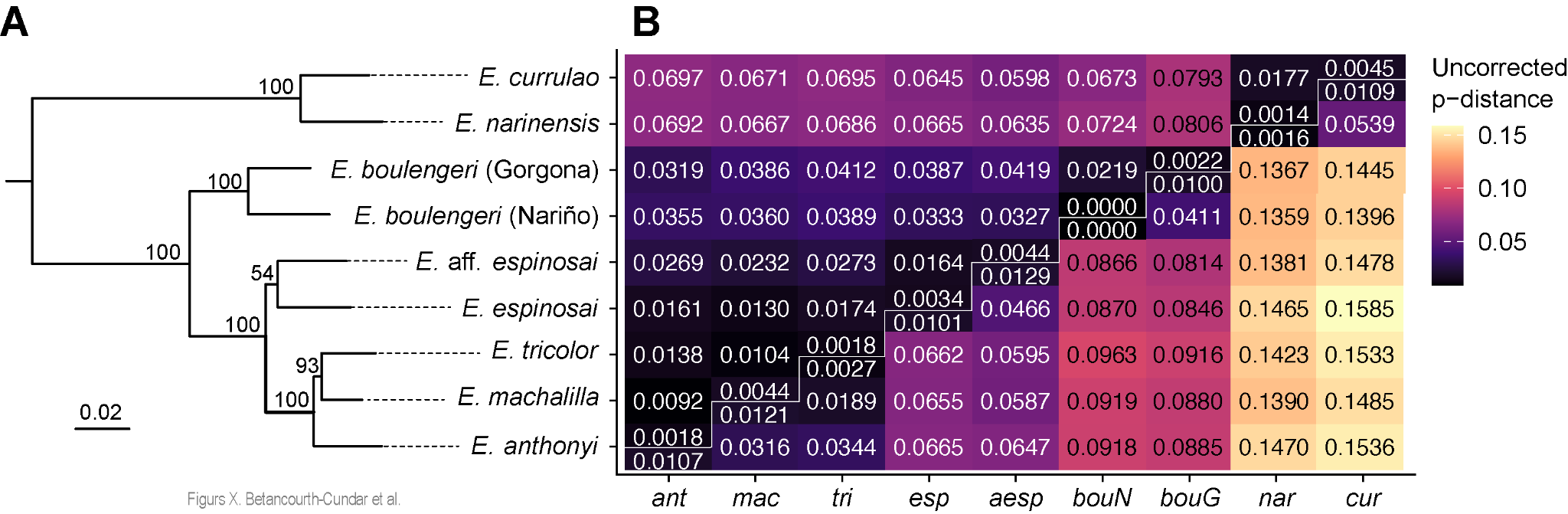

**Figura 5. Posición filogenética y distancias genéticas de *E. currulao* sp. nov. y otras especies de Epipedobates.** Una filogenia de Epipedobates de máxima probabilidad a nivel de especie estimada en IQ-TREE2 utilizando dos genes nucleares (*BMP2*, *H3*) y tres fragmentos de genes mitocondriales (*CYTB, 12S-16S, CR*) (ver Fig. Suppl. material 2 para ver la filogenia completa). Los valores de soporte de nodos de 10.000 réplicas de arranque ultrarrápido indicaron un fuerte soporte de *E. currulao* sp. nov. como especie hermana de *E. narinensis* . B Matriz de distancias p medias por pares dentro y entre especies de Epipedobates. Las distancias del triángulo inferior se basan en *CYTB* y las distancias del triángulo superior son de *12S--16S*. Abreviaturas: cur, *E. currulao* sp. nov; nar, *E. narinensis;* bouN, *E. boulengeri* (Nariño, Colombia); bouG, *E. boulengeri* (Isla Gorgona, Cauca, Colombia); aesp, *E.* aff. *espinosai;* especialmente, *E. espinosai*; tri, *E. tricolor*; mac, *E. machalilla*; ant, *E. anthonyi.* Ver Suppl. material 6 para filogenia completa.

**Morfología**

En *E. currulao* el dedo II es más largo que el dedo III y el dedo IV está hinchado en los machos. En nuestra revisión de la morfología, notamos que los dedos III y V tienen una longitud reducida en *E. currulao* (dedo III/SVL: media ± DE = 16,1% ± 1,59%; dedo V/SVL: 15,1% ± 1,42%, N = 37 ranas) (Fig. 2H–I) y *E. narinensis* (dedo III/SVL: 16,9% ± 0,85%; dedo III/SVL: 16,2% ± 1,08%, N = 8) relativo a *E. boulengeri* -Gorgona (dedo III/SVL: 19,2% ± 0,67%; dedo V/SVL: 18,2% ± 1,12%, N = 9), *E. boulengeri* -Nariño (dedo III/SVL: 17,5% ± 0,99%; dedo V/SVL: 17,7% ± 2,14%, N = 9) o *E.* aff. *espinosai* (dedo III/SVL: 19,3%; dedo V/SVL: 16,3%, N = 1) (Tabla 3, Suppl. material 4).

Existen diferencias significativas en el tamaño corporal entre *Epipedobates currulao* sp. nov., *E. narinensis* y *E. boulengeri* (Gorgona y Nariño) (prueba de Kruskal-Wallis: H3 = 19,058, P < 0,001; Fig. 6A). *Epipedobates boulengeri* - Gorgona exhibe un tamaño corporal promedio significativamente mayor que las otras poblaciones analizadas. Una prueba de Wilcoxon mostró que *E. currulao* sp. nov. es más grande que *E. narinensis* (P = 0.05, N = 33) y más pequeño que *E. boulengeri* - Gorgona (P < 0.006, N = 32). No encontramos diferencias significativas en el tamaño corporal entre *E. currulao* sp. nov. y las poblaciones de Nariño asignadas a *E. boulengeri* (P = 0.462, N = 33). Encontramos dimorfismo sexual en el tamaño corporal (SVL) para *E. currulao* sp. nov. (ANOVA: F1,46 = 6,269, P < 0,01), donde la media del SVL en mujeres fue mayor (media = 18,24, DE = 1,09 mm, N = 28, Tabla 2) que la media en hombres (media = 17,45, DE = 1,07 mm, N = 20, Tabla 1).

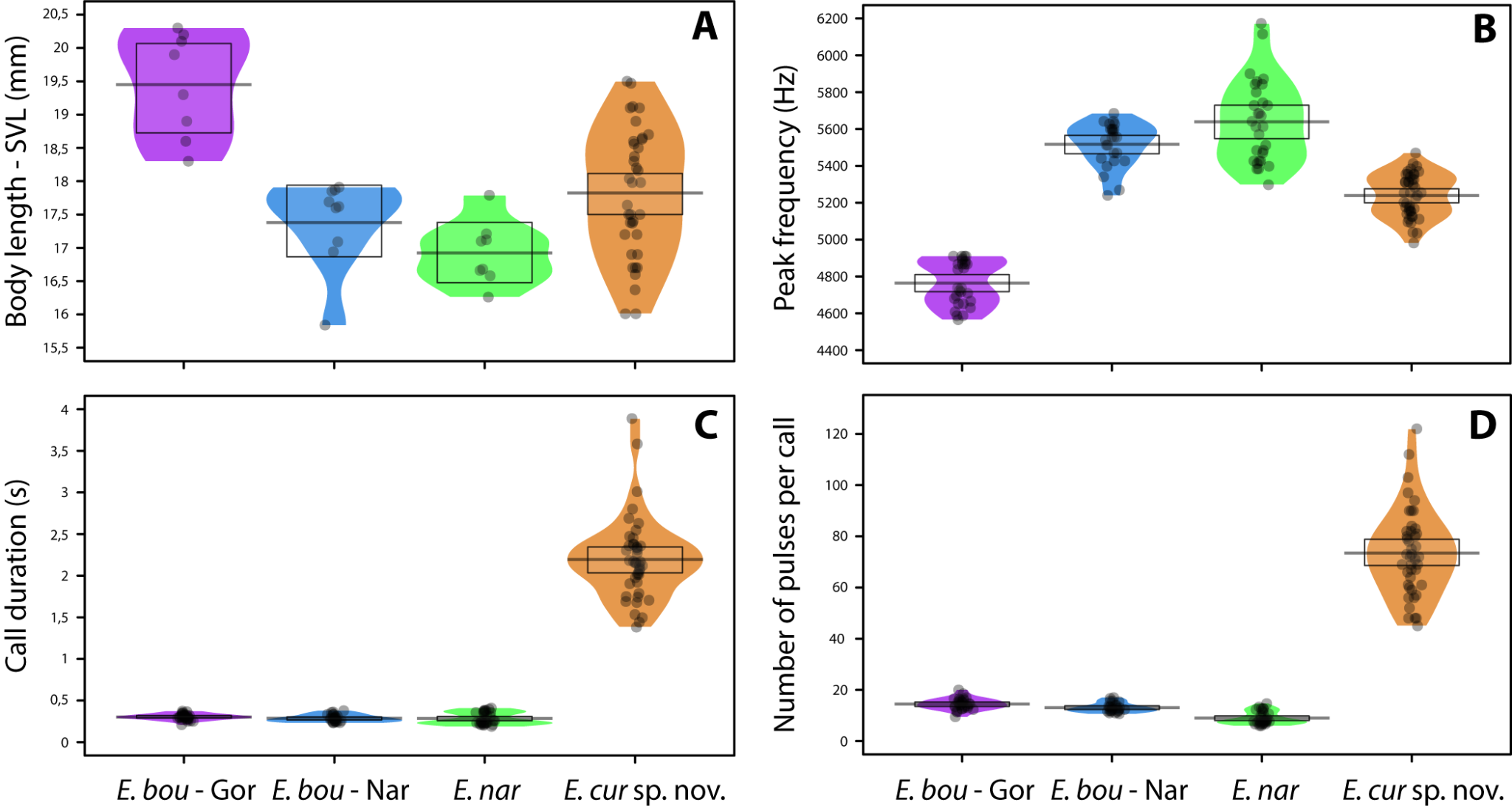

**Figura 6. Comparaciones de SVL y rasgos de llamadas publicitarias entre *E. currulao* sp. nov. y otras especies de Epipedobates encontradas en Colombia.** A Longitud hocico-cloaca (SVL) B Frecuencia máxima C Duración de la llamada y D Número de pulsos por llamada. Las abreviaturas incluyen: E. bou - Gor, *E. boulengeri* - Gorgona; E. bou - Nar, *E. boulengeri* - Nariño; E. nar, *E. narinensis* ; y *E. cur* sp. nov., *E. currulao* sp. nov. Los puntos negros representan datos sin procesar, la barra horizontal muestra la media, el diagrama sombreado es una curva de densidad suavizada que muestra la distribución completa de los datos y el rectángulo representa la incertidumbre alrededor de la media utilizando un intervalo de densidad más alto bayesiano del 95 %.

**Vocalizaciones**

*Epipedobates currulao* sp. nov., *E. narinensis* y *E. boulengeri* (de Gorgona y Nariño) muestran distinciones notables en los parámetros de llamada. Encontramos diferencias significativas en la frecuencia máxima de llamadas publicitarias entre todas las especies y poblaciones analizadas (prueba de Kruskal-Wallis: H _3_ = 109,25, P <0,001; Fig. 6B). La prueba de Wilcoxon demostró que *Epipedobates* *currulao* sp. nov. llamadas con una frecuencia máxima más baja que *E. narinensis* (P < 0.001, N = 81) y *E. boulengeri*-Nariño (P < 0.001, N = 76), pero su frecuencia máxima es mayor que la de *E. boulengeri*-Gorgona (P < 0,001, N = 79). Encontramos diferencias estadísticamente significativas en la duración de la llamada (prueba de Kruskal-Wallis: H_3_ = 95,302, P <0,001; Fig. 6C). *Epipedobates* *currulao* sp. nov. exhibe llamadas más largas que *E. narinensis* (P < 0.001, N = 81), *E. boulengeri* -Nariño (P < 0.001, N = 76) y *E. boulengeri* -Gorgona (P < 0.001, N = 79). Se encontraron resultados similares para el número de pulsos por llamada (prueba de Kruskal-Wallis: H_3_ = 107,52, P <0,001, Fig. 6D). *Epipedobates* *currulao* sp. nov. tiene llamadas con más pulsos que *E. narinensis* (P < 0.001, N = 81), *E. boulengeri*-Nariño (P < 0.001, N = 76) y *E. boulengeri*-Gorgona (P < 0.001, N = 79).

Las convocatorias publicitarias se componen de grupos de convocatorias, una convocatoria para E. currulao sp. nov., 2-3 para *E. boulengeri* - Gorgona, 3-6 para *E. boulengeri* - Nariño y 5-14 para *E. narinensis* (Fig. 7). Además, registramos a dos machos cerca de una hembra con vocalizaciones diferentes a su llamada publicitaria (Fig. 7E). Estas vocalizaciones, muy probablemente una llamada de cortejo, variaron entre 0,21 y 0,31 s de duración (media = 0,25, DE = 0,03 s, N = 2) e incluyeron de 7 a 9 pulsos por llamada (media 8,25 ± DE 0,83). La frecuencia máxima osciló entre 4,78 y 5,17 kHz (media = 4,96, DE = 0,13 kHz).

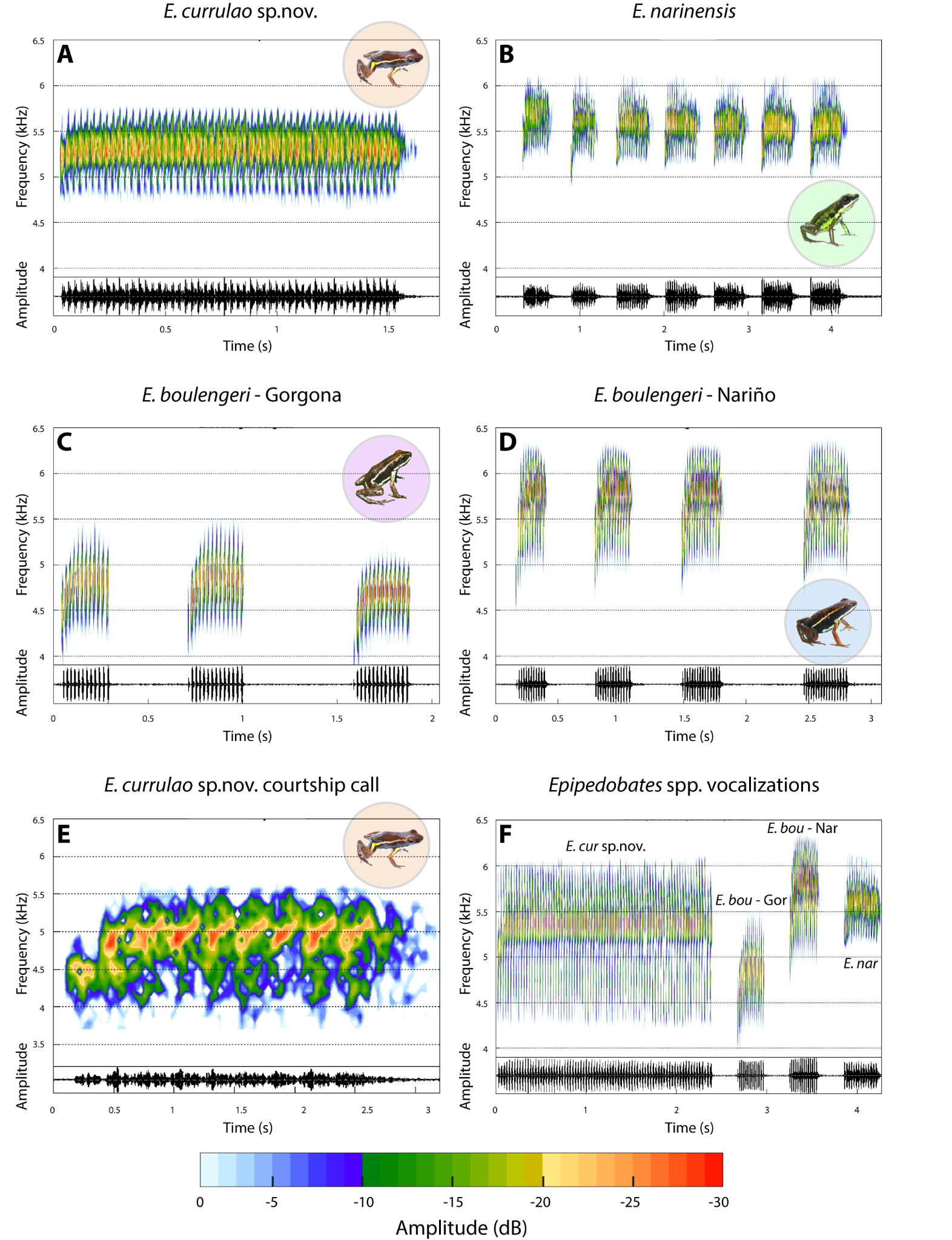

**Figura 7**. **Vocalizaciones en *E. currulao* sp. nov. y otras especies de *Epipedobates* .** Los espectrogramas (arriba) y los oscilogramas (abajo) que muestran aspectos generales de la estructura de un anuncio espontáneo llaman a un macho de **A** *E. currulao* sp. nov. de Ladrilleros, Valle del Cauca, **B.** *E. narinensis* del Biotopo, Nariño, **C** *E. boulengeri*-Gorgona (Localidad tipo) y **D** *E. boulengeri*-Nariño de Maragrícola, Nariño. **E** Espectrogramas y oscilogramas que muestran aspectos generales de la estructura de una llamada de cortejo de *E. currulao* sp. nov. de Ladrilleros, Valle del Cauca. **F** Comparación de una llamada publicitaria entre *E. currulao* sp. nov. y otras especies de *Epipedobates encontradas en Colombia* . *E. bou* - Gor: *E. boulengeri* - Gorgona, *E. bou* - Nar: *E. boulengeri* - Nariño, E. nar: *E. narinensis* y *E. cur* sp. nov.: *E. currulao* sp. nov *.* Transformada rápida de Fourier-FFT = 256, superposición = 90%.

Un análisis de componentes principales (PCA) de los parámetros de las llamadas indicó que los primeros tres componentes principales explican el 88% de la variación en las llamadas (Fig. 8A). PC1 explicó el 45% de la variación y se asoció positivamente con la duración de la llamada, el número de pulsos y el intervalo entre llamadas. PC1 también se relacionó negativamente con la alta frecuencia y el 90% del ancho de banda (Suppl. material S4). PC2 explicó un 33% adicional de la variación en los parámetros de llamada. Se relacionó positivamente con la baja frecuencia y la frecuencia máxima. PC3 explicó el 10% de la variación y se asoció positivamente con el ancho de banda intercuartil de frecuencia del 90% (Fig. 8A, Suppl. material 7). En resumen, PC1 se asoció principalmente con características temporales de la llamada publicitaria y PC2 con características espectrales. La función de análisis discriminante lineal (LDA) (Fig. 8B) utilizando cinco PC mostró que *E. currulao* se puede diferenciar fácilmente de otras especies en función de sus anuncios. LD1 se asoció positivamente con PC1 (características temporales) (coef. LD = 1,71), y LD2 se asoció negativamente con PC2 (características espectrales) (coef. LD = -1,47). Esto significa que *E. currulao* se distingue por tener llamadas más largas y una mayor cantidad de pulsos (Figs. 8B, 6CD). Con respecto a la frecuencia baja y máxima, las llamadas de *E. currulao* se caracterizan por valores intermedios en comparación con las otras especies analizadas (frecuencia máxima: 4,84 ± 0,22 kHz, N = 15). *Epipedobates boulengeri-* Gorgona tiene la frecuencia máxima más baja (4,76 ± 0,11 kHz, N = 5) y *E. narinensis* tiene la frecuencia máxima más alta (5,64 ± 0,21 kHz, N = 6) (Figs. 8B y 6B).

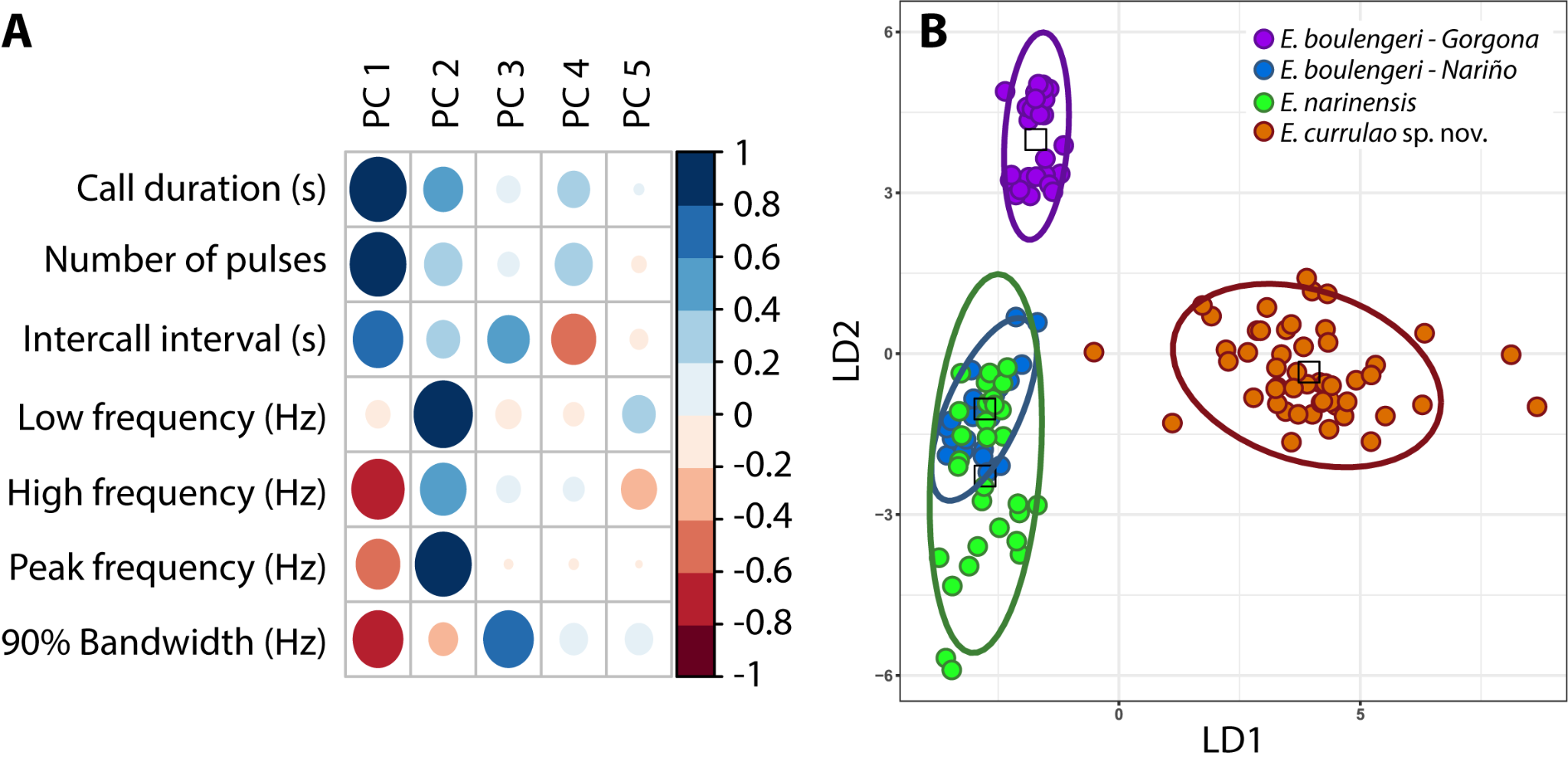

**Figura 8. Diferencias en los parámetros acústicos de *E. currulao* sp. nov. en comparación con otras especies de *Epipedobates encontradas en Colombia*.** **A** La contribución relativa de cada variable acústica a los primeros cinco componentes principales obtenidos en el Análisis de Componentes Principales. El valor absoluto de cada contribución se representa según el tamaño del círculo, mientras que los colores azul y rojo muestran contribuciones positivas y negativas, respectivamente. **B** Un análisis discriminante lineal que utiliza PC como entrada indica que *E. currulao* se puede diferenciar de otras especies en función de sus llamadas publicitarias, particularmente con respecto a la duración de la llamada y la cantidad de pulsos (PC1 se carga positivamente en LD1). Los puntos indican los individuos, el color indica la especie y las elipses indican los intervalos de confianza del 95% para los puntos de datos LDA.

**DISCUSIÓN**

Describimos *Epipedobates currulao* sp. nov. como una nueva especie encontrada a lo largo de la selva chocoana del Pacífico de Colombia, que anteriormente se consideraba parte del complejo de especies de *E. boulengeri* (López-Hervas et al., 2024). Describimos esta especie como nueva basándonos en trabajos anteriores (López-Hervas et al., 2024) que demuestran su divergencia con su especie hermana, *E. narinensis*, y su coloración única. Aquí proporcionamos evidencia adicional con la caracterización de su distintivo llamado publicitario.

*Epipedobates currulao* lleva el nombre en honor al género musical y las tradiciones afrocolombianas del currulao que surgieron de generaciones de prácticas multiculturales en la región del Pacífico, comenzando con los esclavos africanos llevados a trabajar en las minas de oro y continuando hoy con una infusión de nuevas ideas e interpretaciones. (Birenbaum Quintero, 2019; Guevara Calderón & Godoy Acosta, 2015). Como parte del patrimonio cultural de las comunidades negras de la región del Pacífico, la música currulao, también conocida como bambuco viejo, contribuyó a reforzar argumentos que llevaron al reconocimiento de los derechos territoriales y la identidad cultural de los afrocolombianos a través de la Ley 70 de 1993 (Birenbaum Quintero, 2019).

Será necesaria más información para evaluar el estado de conservación de *E. currulao*, pero proponemos que pueda ser considerado como Vulnerable o Casi Amenazado según la categorización de la UICN, dado su rango relativamente pequeño. Aunque no se sabe que *E. currulao* sea simpátrica con otras especies de *Epipedobates*, las regiones del suroeste de Colombia (especialmente al este de Guapi) no han sido estudiadas exhaustivamente y pueden contener sitios con simpatría entre *E. currulao*, *E. boulengeri* y/o *E.* *espinosai*. *Epipedobates currulao* ocurre en simpatría y puede confundirse con *Andinobates minutus* a menos que se distingan cuidadosamente los patrones de color (consulte la sección Diagnóstico diferencial más arriba). Las otras especies del género *Epipedobates* se encuentran a lo largo de la costa occidental y las estribaciones andinas de Ecuador y Perú. Aunque se cree que una especie ( *Epipedobates maculatus* ) se encuentra en el oeste de Panamá (Grant et al., 2017; Jungfer, 2017), se ha cuestionado su asignación a este género (López-Hervas et al., 2024).

Encontramos que la llamada publicitaria de *E. currulao* es única en comparación con otros *Epipedobates* distribuidos en Colombia. Anteriormente, Lötters et al. (2003) reportaron diferencias en llamadas publicitarias entre *Epipedobates boulengeri* registradas en Anchicayá (ahora *E. currulao*) y en Lita (ahora *E.* aff. *espinosai*), sugiriendo en ese momento que las poblaciones pueden pertenecer a un complejo de especies. En nuestras grabaciones encontramos que *E. currulao* realiza sólo una llamada por serie , aunque Lötters et al. (2003) describen series de llamadas con hasta tres notas (las notas son llamadas según nuestra definición); sin embargo, registraron sólo un macho. Visualmente, al observar el estudio de Lötters et al. (2003) al grabar, parece que la llamada se divide en grupos de notas. Sin embargo, al comparar los tiempos entre notas (0,020-0,025 s) de la llamada , observamos que son similares a nuestras mediciones para los tiempos entre pulsos (0,001 – 0,068 s; media = 0,013, SD = 0,009 s, norte = 15). Así , consideramos que estas “notas” comprender una sola llamada, y que la serie de llamadas de *E. currulao* contiene una llamada relativamente larga con muchos pulsos .

Proporcionamos datos sobre el llamado de cortejo de *E. currulao*, que hasta donde sabemos es el primer llamado de cortejo reportado para *Epipedobates*. Al igual que los cantos de cortejo de otras ranas venenosas neotropicales, su frecuencia es menor que el canto publicitario de *E. currulao* (Caldeira Costa et al., 2006; González-Santoro et al., 2021; Marques Correia da Rocha et al., 2018; Moss et al., 2023). También fue más corto y tuvo menos pulsos (7,9 vs 22 – 122 por llamada).

La densidad de *Epipedobates* varía según las especies y sitios de Colombia. Durante nuestro trabajo de campo notamos que las poblaciones de *E. boulengeri* en Isla Gorgona y Maragrícola son bastante abundantes. Por ejemplo, se puede escuchar a más de cinco individuos llamando al mismo tiempo, por lo que fue muy difícil grabar a un solo macho. Para *E. narinensis* en la reserva natural de Biotopo, los individuos cantaban con menos densidad pero aún eran abundantes, con una distancia de 8 a 10 m entre los machos que cantaban. Pero en otros lugares cercanos sólo hemos escuchado a dos o tres individuos en una caminata de 3 a 4 horas. Los individuos de *E. currulao* sp. nov. están más dispersos y se observó que los machos cantaban a aproximadamente 4 a 5 m de distancia entre sí en la localidad tipo (Ladrilleros) y cerca en La Barra. En Anchicayá, los machos cantan cada 3 a 4 m y se asocian con pequeños arroyos al borde del camino de terracería. No sabemos el motivo de las diferencias en densidad, ya que el patrón no corresponde necesariamente a diferencias en la calidad del hábitat. Si bien Isla Gorgona es un parque nacional, *E. boulengeri* parece ser más abundante en las áreas perturbadas adyacentes a los edificios de las estaciones de campo. El sitio Maragrícola donde tomamos muestras de *E. boulengeri* en Nariño es una plantación experimental de cacao. Ladrilleros y La Barra son sitios perturbados similares a donde E. boulengeri era abundante en Isla Gorgona, aunque *E. currulao* no se presentó en densidades tan altas. Es posible que *E. currulao* sea más vulnerable a la transformación del hábitat que E. boulengeri, pero se necesita investigación adicional para comprender qué impulsa las diferencias en densidad entre sitios y especies.

Desafortunadamente, no recolectamos estadios larvarios de *E. currulao.* La morfología larvaria de Dendrobatoidea ha contribuido a la comprensión de las relaciones filogenéticas dentro de la superfamilia (Anganoy-Criollo & Cepeda-Quilindo, 2017). Sería interesante recolectar renacuajos de la nueva especie para compararlos con otras especies de Epipedobates. Las comparaciones de estadios larvarios realizadas por Anganoy-Criollo y Cepeda-Quilindo (2017) detectaron algunas diferencias entre las poblaciones de Ecuador y el suroeste de Colombia en la forma del cuerpo, puntas de las papilas marginales y patrón de coloración de la cola. Sin embargo, los ejemplares de Colombia utilizados en su estudio probablemente correspondan a *E. boulengeri* - Nariño (*E. boulengeri* sensu estricto) y no tenemos certeza de la identificación de los ejemplares de Ecuador, que podrían corresponder a *E.* aff. *espinosai*, *E. boulengeri* sensu estricto o *E. espinosai* , todos los cuales se encuentran en la región (López-Hervas et al., 2024).

**AGRADECIMIENTOS**

RDT y MBC agradecen a Cristian Flórez Pai, Luis Alfredo Esteves, Fray Arriaga, Sandra V. Flechas, Daniel Nastacuaz y Nataly Portillo por su asistencia en el trabajo de campo. Adolfo Amézquita facilitó el proceso de permisos en 2014 y 2016 y Mireya Osorio R. en 2022. También agradecemos al personal del museo de ANDES, QCAZ y MVZ, incluidas Angela Sánchez G. y Carol Spencer, por su ayuda en la adquisición de especímenes. Agradecemos a Valeria Ramírez-Castañeda por la ayuda con la PCR. Agradecemos a Mauricio Rivera por el acceso a los especímenes del Instituto de Ciencias Naturales de la Universidad Nacional de Colombia, Bogotá, y a Nataly Casas por su ayuda en la toma de fotografías de especímenes del museo. Agradecemos al Dr. Stefan Lötters y al Dr. Andrés F. Jaramillo-Martínez por su ayuda en la revisión del manuscrito.

**Información adicional**

**Conflicto de intereses**

Los autores han declarado que no existen intereses contrapuestos.

**Declaración ética**

No se informó ninguna declaración ética.

**Fondos**

El apoyo financiero para este proyecto para RDT fue proporcionado por NSF (IOS 2319711), financiación inicial de la Universidad de California Berkeley y subvenciones de la Sociedad de Biólogos Sistemáticos, la Sociedad Herpetológica de Carolina del Norte, la Sociedad para el Estudio de Reptiles y Anfibios, Chicago. Sociedad Herpetológica, Sociedad Herpetológica de Texas, Programa EEB de la Universidad de Texas en Austin, Programa de becas de investigación para graduados de la Fundación Nacional de Ciencias Oportunidades de investigación para graduados en todo el mundo, Sociedad Geográfica Nacional (Beca para jóvenes exploradores n.° 9468-14). NSF también brindó apoyo a DCC y RDT (DEB 1556967).

**Contribuciones de autor**

Adquisición de financiación: RDT, DCC. Conceptualización: MBC, RDT. Investigación: MBC, RDT, JCR. Metodología: MBC, RDT, JCR, AJC. Análisis formal: MBC. Redacción – borrador original: MBC, RDT. Redacción – revisión y edición: MBC, RDT, DCC, JCR, AJC.

**Autor ORCID**

Mileidy Betancourth-Cundar <https://orcid.org/0000-0003-2368-6028>

Juan Camilo Ríos-Orjuela <https://orcid.org/0000-0001-6976-9131>

Andrew J. Crawford https://orcid.org/0000-0003-3153-6898

David C. Cannatella <https://orcid.org/0000-0001-8675-0520>

Rebecca D. Tarvin <https://orcid.org/0000-0001-5387-7250>

**Disponibilidad de datos**

Todos los datos que respaldan los hallazgos de este estudio están disponibles en el texto principal o en Suppl. material y se cargaron en bases de datos públicas.

**REFERENCIAS**

Abadía G (1973) La música folklórica colombiana. Universidad Nacional de Colombia, Dirección de Divulgación Cultural, Bogota D.C.- Colombia, 158 pp.

Anganoy-Criollo M, Cepeda-Quilindo B (2017) Redescription of the tadpoles of *Epipedobates narinensis* and *E. boulengeri* (Anura: Dendrobatidae). Phyllomedusa, 16(2), 155–182. https://doi.org/10.11606/issn.2316-9079.v16i2p155-182

Aristizabal M (2002) El festival del Currulao en Tumaco: Dinámicas culturales y construcción de identidad étnica en el litoral pacífico colombiano. Master Thesis, Universidad del Valle, Valle del Cauca, Colombia. <https://www.humanas.unal.edu.co/colantropos/files/2914/5615/3098/aristizabal_currulao.pdf>

Barbour T (1909) Corrections regarding the names of two recently described Amphibia Salientia. Proceedings of the Biological Society of Washington, 22, 89.

Bioacoustics Research Program (2014) Raven Pro: Interactive Sound Analysis Software (Version 1.5) [Computer software] (1.5). The Cornell Lab of Ornithology. <http://www.birds.cornell.edu/raven>

Birenbaum Quintero M (2006) “La música pacífica” al Pacífico violento: Música, multiculturalismo y marginalización en el Pacífico negro colombiano. Trans. Revista Transcultural de Música, 10, 1–32. <https://www.redalyc.org/pdf/822/82201002.pdf>

Birenbaum Quintero M (2019) Rites, Rights & Rhythms A Genealogy of Musical Meaning in Colombia’s Black Pacific. Oxford University Press, Canada, 342 pp. <https://www.google.com/books/edition/Rites_Rights_and_Rhythms/YtB0DwAAQBAJ>

Boulenger GA (1899) Descriptions of new reptiles and batrachians collected by Mr. P. O. Simons in the Andes of Ecuador. Annals and Magazine of Natural History, Series 7, 4, 454–457.

Brown J, Twomey E, Amézquita A, Barbosa De Souza M, Caldwell J, Lötters S, Von May R, Melo-Sampaio P, Mejía-Vargas D, Perez-Peña P, Pepper M, Poelman E, Sanchez-Rodriguez M, & Summers K (2011) A taxonomic revision of the Neotropical poison frog genus *Ranitomeya* (Amphibia: Dendrobatidae). Zootaxa, 120(3083), 1–120. <https://doi.org/10.11646/zootaxa.3083.1.1>

Caldeira Costa R, Gomes Facure K, Giaretta A (2006) Courtship, vocalization, and tadpole description of *Epipedobates flavopictus* (Anura: Dendrobatidae) in southern Goiás, Brazil. Biota Neotropica, 6(1). <https://doi.org/10.1590/S1676-06032006000100006>

Chao K-H, Barton K, Palmer S, Lanfear R (2021) sangeranalyseR: Simple and Interactive Processing of Sanger Sequencing Data in R. Genome Biology and Evolution, 13(3), 1–7. <https://doi.org/10.1093/gbe/evab028>

Cisneros-Heredia D, Yánez-Muñoz M (2010) A new poison frog of the genus *Epipedobates* (Dendrobatoidea: Dendrobatidae) from the north-western Andes of Ecuador. Avances, 2(3), B83–86. <http://www.cisneros-heredia.org/pdfs/2010_Epipedobatesdarwinwallacei.pdf>

Clough M, Summers K (2000) Phylogenetic systematics and biogeography of the poison frogs: evidence from mitochondrial DNA sequences. Biological Journal of the Linnean Society, 70(3), 515–540. <https://doi.org/10.1006/bijl.1999.0418>

Coloma L (1995) Ecuadorian Frogs of the Genus *Colostethus* (Anura: Dendrobatidae). Miscellaneous Publication. Museum of Natural History, University of Kansas, 87, 1–72. <https://doi.org/10.5962/bhl.title.16171>

Cruz Hoyos S (2016) “Samuelito”, el hombre que hizo del currulao una danza universal. [Https://www.Elpais.Com.Co/Entretenimiento/Cultura/Samuelito-El-Hombre-Que-Hizo-Del-Currulao-Una-Danza-Universal.Html](https://www.elpais.com.co/Entretenimiento/Cultura/Samuelito-El-Hombre-Que-Hizo-Del-Currulao-Una-Danza-Universal.Html).

Edgar RC (2004) MUSCLE: Multiple sequence alignment with high accuracy and high throughput. Nucleic Acids Research, 32(5), 1792–1797. <https://doi.org/10.1093/nar/gkh340>

Erdtmann L, Amézquita A (2009) Differential evolution of advertisement call traits in dart-poison frogs (Anura: Dendrobatidae). Ethology, 115(9), 801–811. <https://doi.org/10.1111/j.1439-0310.2009.01673.x>

Funkhouser JW (1956) New frogs from Ecuador and southwestern Colombia. Zoologica. New York, 41, 73–80.

Globally Unique Identifiers Task Group (2011) GUID and Life Sciences Identifiers Applicability Statements. Biodiversity Information Standards (TDWG). <http://www.tdwg.org/standards/150>

Goebel AM, Donnelly JM, Atz ME (1999) PCR Primers and Amplification Methods for 12S Ribosomal DNA, the Control Region, Cytochrome Oxidase I, and Cytochromebin Bufonids and other Frogs, and an overview of PCR Primers which have amplified DNA in amphibians auccessfully. Molecular Phylogenetics and Evolution, 11(1), 163–199. <https://doi.org/10.1006/MPEV.1998.0538>

González-Santoro M, Hernández-Restrepo J, Palacios-Rodríguez P (2021) Aggressive behaviour, courtship and mating call description of the neotropical poison frog *Phyllobates aurotaenia* (Anura: Dendrobatidae). Herpetology Notes, 14, 1145–1149. <https://www.biotaxa.org/hn/article/view/60982>

Grant T, Frost DR, Caldwell JP, Gagliardo R, Haddad CFB, Kok PJR, Means DB, Noonan BP, Schargel WE, Wheeler WC (2006) Phylogenetic systematics of dart-poison frogs and their relatives (Amphibia: Athesphatanura: Dendrobatidae). Bulletin of the American Museum of Natural History, 299, 1–262. [https://doi.org/dx.doi.org/10.1206/0003-0090(2006)299[1:PSODFA]2.0.CO;2](https://doi.org/dx.doi.org/10.1206/0003-0090(2006)299%5B1:PSODFA%5D2.0.CO;2)

Grant T, Rada M, Anganoy-Criollo M, Batista A, Dias PH, Jeckel AM, Machado DJ, Rueda-Almonacid JV (2017) Phylogenetic Systematics of Dart-Poison Frogs and Their Relatives Revisited (Anura: Dendrobatoidea). South American Journal of Herpetology, 12(s1), S1–S90. <https://doi.org/10.2994/SAJH-D-17-00017.1>

Guevara Calderón A, Godoy Acosta C (2015) El Currulao: una propuesta de exploración, interpretación y creación, a través de la batería y la guitarra eléctrica. Master Thesis, Universidad del Bosque, Bogotá D.C. - Colombia.

Jungfer K-H (2017) On Warszewicz’s trail: The identity of *Hyla molitor* O. SCHMIDT, 1857. Salamandra, 53(1), 18–24. <https://www.researchgate.net/publication/316542564>

Kalyaanamoorthy S, Minh BQ, Wong TKF, Von Haeseler A, Jermiin LS (2017) modelfinder: fast model selection for accurate phylogenetic estimates. 14(6). <https://doi.org/10.1038/nmeth.4285>

Kampstra P (2008) Beanplot: A Boxplot Alternative for Visual Comparison of Distributions. Journal of Statistical Software, 28:1–9. <https://doi.org/10.18637/jss.v028.c01>

Köhler J, Jansen M, Rodriguez A, Kok PJR, Toledo LF, Emmrich M, Glaw F, Haddad CFB, Rödel M-O, Vences M (2017) The use of bioacoustics in anuran taxonomy: Theory, terminology, methods and recommendations for best practice. Zootaxa, 4251(1), 1–124. <https://doi.org/https://doi.org/10.11646/zootaxa.4251.1.1>

Larsson A (2014) AliView: a fast and lightweight alignment viewer and editor for large datasets. Bioinformatics, 30(22), 3276–3278. <https://doi.org/10.1093/BIOINFORMATICS/BTU531>

Lê S, Josse J, Husson F (2008) FactoMineR: An R Package for multivariate analysis. Journal of Statistical Software, 25(1), 1–18. <https://doi.org/10.18637/JSS.V025.I01>

López-Hervas K, Santos J, Ron S, Betancourth-Cundar M, Cannatella D, Tarvin R (2023) Deep divergences among cryptic clades of *Epipedobates* poison frogs. bioRxiv, 547117. <https://doi.org/10.1101/2023.06.29.547117>

Lötters S, Reichle S, Jungfer K-H (2003) Advertisement calls of Neotropical poison frogs (Amphibia: Dendrobatidae) of the genera *Colostethus*, *Dendrobates* and *Epipedobates*, with notes on dendrobatid call classification. Journal of Natural History, 37(15), 1899–1911. <https://doi.org/10.1080/00222930110089157>

Lötters S, Jungfer KH, Henkel FW, Schmidt W (2007) Poison Frogs. Biology, Species Captive Maintenance. Frankfurt am Main: Edition Chimaira.

Lynch JD, Suárez-Mayorga AM (2004) Catálogo de anfibios en el Chocó Biogeografico. pp. 654- 668. In: J. O. Rangel (ed.). Colombia Diversidad Biotica IV. El Chocó Biogeografico. Universidad Nacional de Colombia, Bogotá, D.C.

Marques Correia da Rocha S, Pimentel Lima A, Kaefer IL (2018) Reproductive Behavior of the Amazonian Nurse-Frog *Allobates paleovarzensis* (Dendrobatoidea, Aromobatidae). South American Journal of Herpetology, 13(3), 260–270. <https://doi.org/10.2994/SAJH-D-17-00076.1>

Minh BQ, Nguyen MAT, Von Haeseler A (2013) Ultrafast approximation for phylogenetic bootstrap. Molecular Biology and Evolution, 30(5), 1188–1195. <https://doi.org/10.1093/MOLBEV/MST024>

Minh BQ, Schmidt HA, Chernomor O, Schrempf D, Woodhams MD, Von Haeseler A, Lanfear R, Teeling E (2020) IQ-TREE 2: New models and efficient methods for phylogenetic inference in the genomic era. Molecular Biology and Evolution, 37(5), 1530–1534. <https://doi.org/10.1093/MOLBEV/MSAA015>

Moss JB, Tumulty JP, Fischer EK (2023) Evolution of acoustic signals associated with cooperative parental behavior in a poison frog. Proceedings of the National Academy of Sciences of the United States of America, 120(17), 1–7. <https://doi.org/10.1073/PNAS.2218956120/-/DCSUPPLEMENTAL>

Mueses-Cisneros JJ, Cepeda-Quilindo B, Moreno-Quintero V (2008) Una nueva especie de *Epipedobates* (Anura: Dendrobatidae) del suroccidente de Colombia. Papéis Avulsos de Zoologia (São Paulo), 48(1), 1–10. <https://doi.org/10.1590/S0031-10492008000100001>

Myers CW, Daly JW (1976) Preliminary evaluation of skin toxins and vocalizations in taxonomic and evolutionary studies of poison-dart frogs (Dendrobatidae). Bulletin of the American Museum of Natural History, 157(3), 173–262.

Myers N, Mittermeier RA, Mittermeier CG, Da Fonseca GAB, Kent J (2000) Biodiversity hotspots for conservation priorities. Nature, 403, 853–858. <https://doi.org/10.1038/35002501>

Noble GK (1921) Five new species of Salientia from South America. American Museum Novitates, 29, 1–7. <http://hdl.handle.net/2246/4615>

Paradis E, Schliep K (2019) ape 5.0: an environment for modern phylogenetics and evolutionary analyses in R. Bioinformatics, 35(3), 526–528. <https://doi.org/10.1093/BIOINFORMATICS/BTY633>

Peters WCH (1873) Über eine neue Schildrötenart, Cinosternon Effeldtii und einige andere neue oder weniger bekannte Amphibien. Monatsberichte der Königlichen Preussische Akademie des Wissenschaften zu Berlin, 603–618.

Phillips N (2017) yarrr: a companion to the e-book “YaRrr!: the pirates guide to R.

Powers RP, Jetz W (2019) Global habitat loss and extinction risk of terrestrial vertebrates under future land-use-change scenarios. Nature Climate Change, 9(4), 323–329. <https://doi.org/10.1038/s41558-019-0406-z>

R Core Team (2023) R: A Language and Environment for Statistical Computing. R Foundation for Statistical Computing. [https://www.R-project.org/](https://www.r-project.org/).

Santos JC, Baquero M, Barrio-Amorós C, Coloma LA, Erdtmann LK, Lima AP, Cannatella DC (2014). Aposematism increases acoustic diversification and speciation in poison frogs. Proceedings of the Royal Society B, 281, 20141761. <https://doi.org/10.1098/rspb.2014.1761>

Santos JC, Cannatella DC (2011) Phenotypic integration emerges from aposematism and scale in poison frogs. Proceedings of the National Academy of Sciences of the United States of America, 108(15), 6175–6180. <https://doi.org/10.1073/pnas.1010952108>

Santos JC, Coloma LA, Cannatella DC (2003) Multiple, recurring origins of aposematism and diet specialization in poison frogs. Proceedings of the National Academy of Sciences of the United States of America, 100(22), 12792–12797. <https://doi.org/10.1073/pnas.2133521100>

Santos JC, Coloma LA, Summers K, Caldwell JP, Ree R, Cannatella DC (2009) Amazonian amphibian diversity is primarily derived from late Miocene Andean lineages. PLoS Biology, 7 e1000056(3), 0448–0461. <https://doi.org/10.1371/journal.pbio.1000056>

Silverstone PA (1976) A revision of the poison-arrow frogs of the genus *Phyllobates* Bibron in Sagra Natural History Museum of Los Angeles County Science Bulletin, 27.

Sueur J, Aubin T, Simonis C (2008) Seewave, a free modular tool for sound analysis and synthesis. Bioacoustics, 18, 213–226. <http://cran.r-project.org/src/contrib/Descriptions/>

Tarvin RD, Powell EA, Santos JC, Ron SR, Cannatella DC (2017) The birth of aposematism: High phenotypic divergence and low genetic diversity in a young clade of poison frogs. Molecular Phylogenetics and Evolution, 109, 283–295. <https://doi.org/10.1016/j.ympev.2016.12.035>

Vargas-S F, Bolaños-L ME (1999) Anfibios y reptiles en hábitats perturbados de selva lluviosa Tropical en el Bajo Anchicayá, Pacifico Colombiano. Revista Academia Ciencias Exactas, Físicas y Naturales 23 (Suplemento Especial), 499–511.

Venables W, Ripley B (2002) Modern Applied Statistics with S (Fourth Edition). Springer. <https://www.stats.ox.ac.uk/pub/MASS4/>.

Vences M, Kosuch J, Boistel R, Haddad CFB, La Marca E, Lötters S, Veith M (2003) Convergent evolution of aposematic coloration in Neotropical poison frogs: a molecular phylogenetic perspective. Organisms Diversity & Evolution, 3, 215–226. <http://www.urbanfischer.de/journals/ode>

Warren R, Vanderwal J, Price J, Welbergen JA, Atkinson I, Ramirez-Villegas J, Osborn TJ, Jarvis A, Shoo LP, Williams SE, Lowe J (2013) Quantifying the benefit of early climate change mitigation in avoiding biodiversity loss. Nature Climate Change, 3(7), 678–682. <https://doi.org/10.1038/nclimate1887>

Watters JL, Cummings ST, Flanagan RL, Siler CD (2016) Review of morphometric measurements used in anuran species descriptions and recommendations for a standardized approach. Zootaxa, 4072(4), 477–495. <https://doi.org/10.11646/zootaxa.4072.4.6>
