## Supplementary material for "Honoring the Afro-Colombian musical culture with the naming of *Epipedobates currulao* sp. nov. (Anura, Dendrobatidae), a frog from the Pacific rainforests": Supplementary Material Information.docx

**Suppl. material 1**: Excel file with details of the specimens examined including their museum numbers, locality information, GenBank accession numbers, and notes regarding species ID and skin removal.

**Suppl. material 2**: A zip file containing the alignments used for distance calculations (AllEpipedobates_cytb_haplotypes.fasta and AllEpipedobates_12S_haplotypes.fasta) and tree inference (Epipedobates-6Genes_updated_2024-07-08.nexus), the partition file for IQ-TREE (Epipedobates-6Genes-120-FINAL-Partitions-Invar-Omitted_2024-07-08.nex), the script used to run IQ-TREE (run_Epipedobates_6Genes_2024-07-08.sh), and the optimal likelihood tree obtained from IQ-TREE with bootstrap values in newick format (revised_2024-07-08.treefile.tre).

**Suppl. material 3**: Lateral, dorsal, and ventral images of the type series of *Epipedobates currulao.*

**Suppl. material 4**: Morphological measurements per individual in mm.

**Suppl. material 5**: Detailed comparisons of coloration across *Epipedobates* species based on Suppl. material 3 and images from López-Hervas et al. (2024).

**Suppl. material 6**: The optimal likelihood tree obtained from IQ-TREE with bootstrap values and specimen information.

**Suppl. material 7**: Results of the Principal Component Analysis and Linear Discriminant Analysis for advertisement call characteristics of *E. currulao* sp. nov. and other species of the genus.

**Suppl. material 8:** A translation of the manuscript into Spanish. (Una traducción del manuscrito al español.)
