## Supplementary figures and images for "Honoring the Afro-Colombian musical culture with the naming of *Epipedobates currulao* sp. nov. (Anura, Dendrobatidae), a frog from the Pacific rainforests"

### S3A_Ladrilleros.png

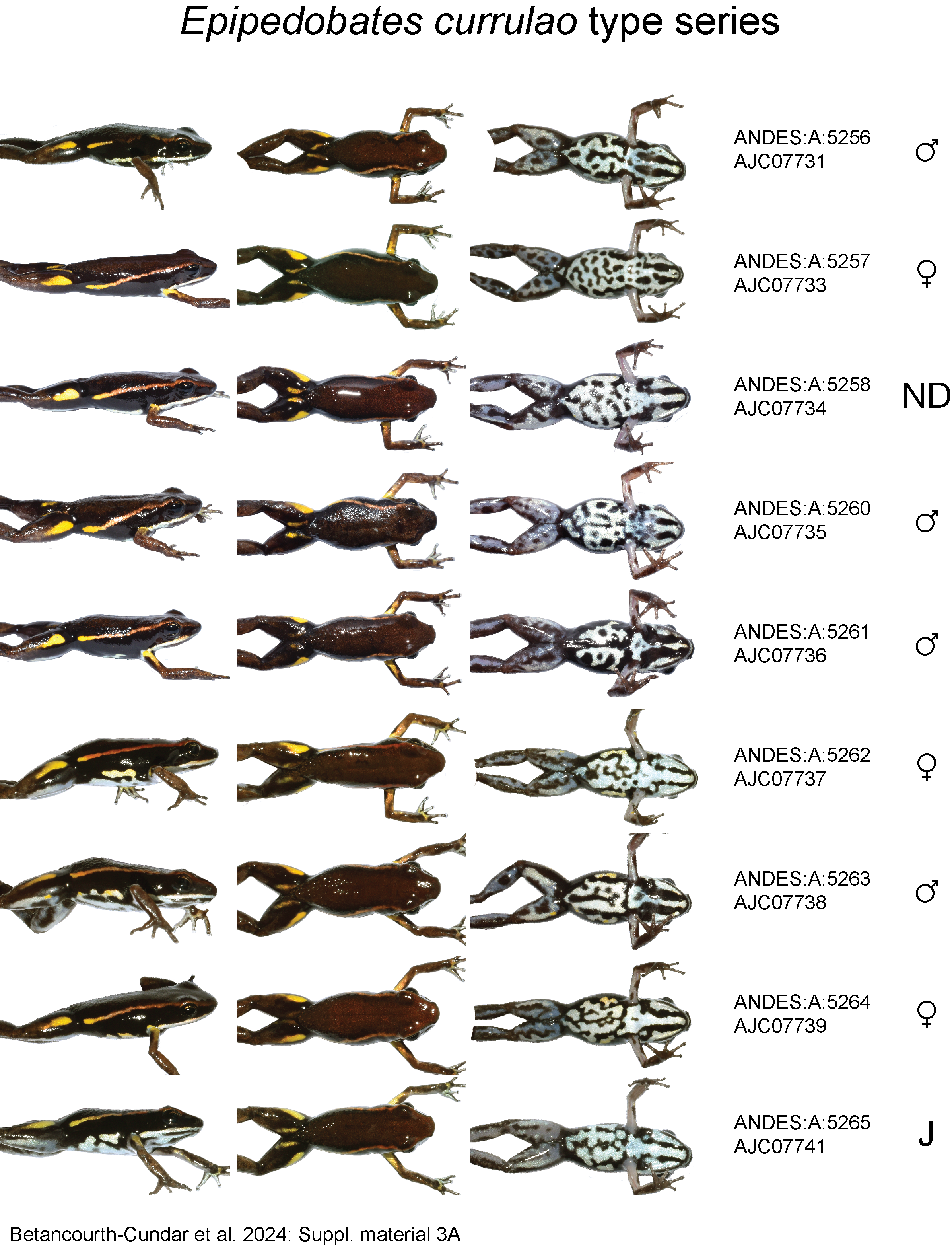

### S3B_Ladrilleros.png

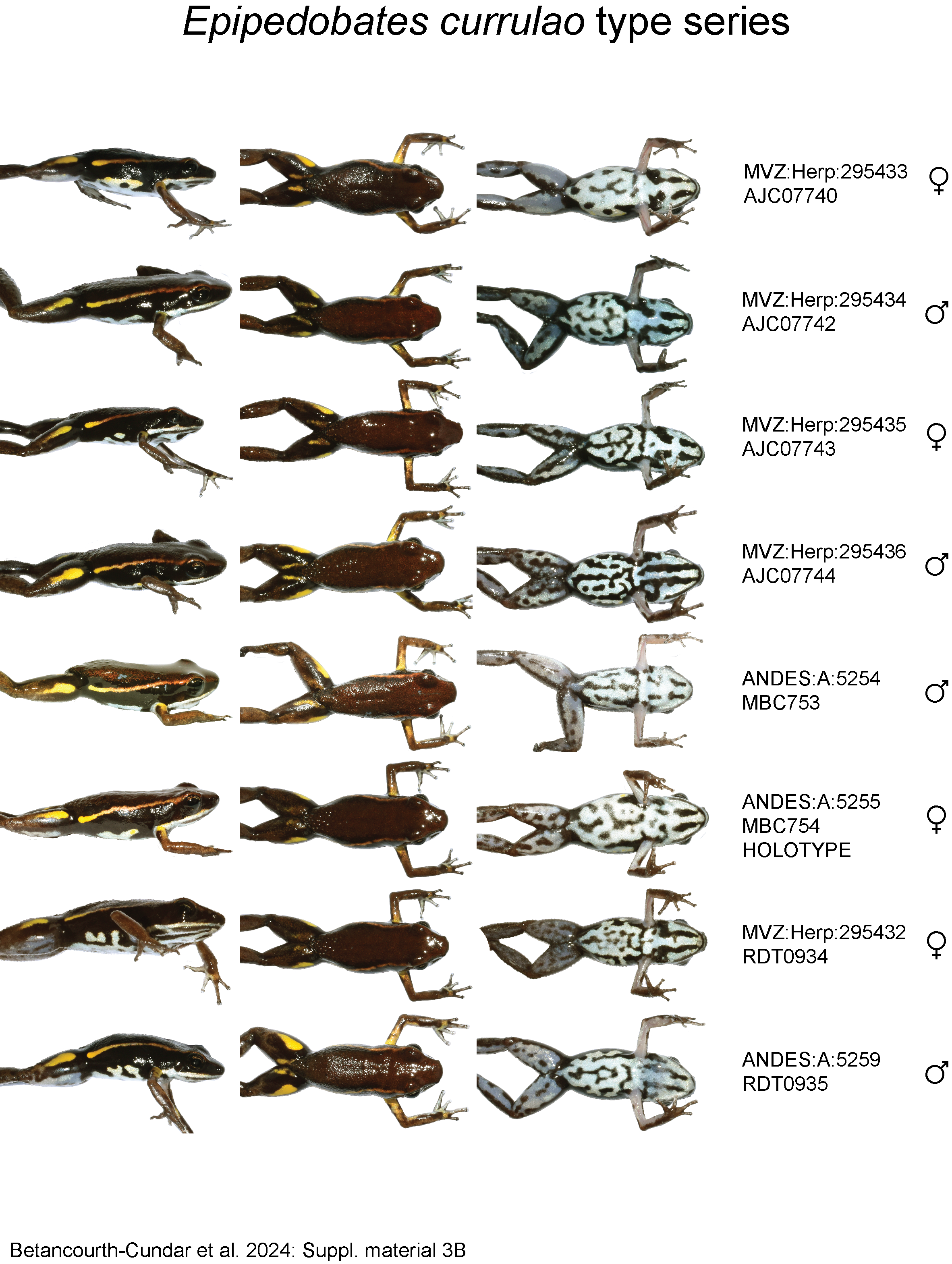

### S6_revised_2024-07-08.treefile.tre.pdf

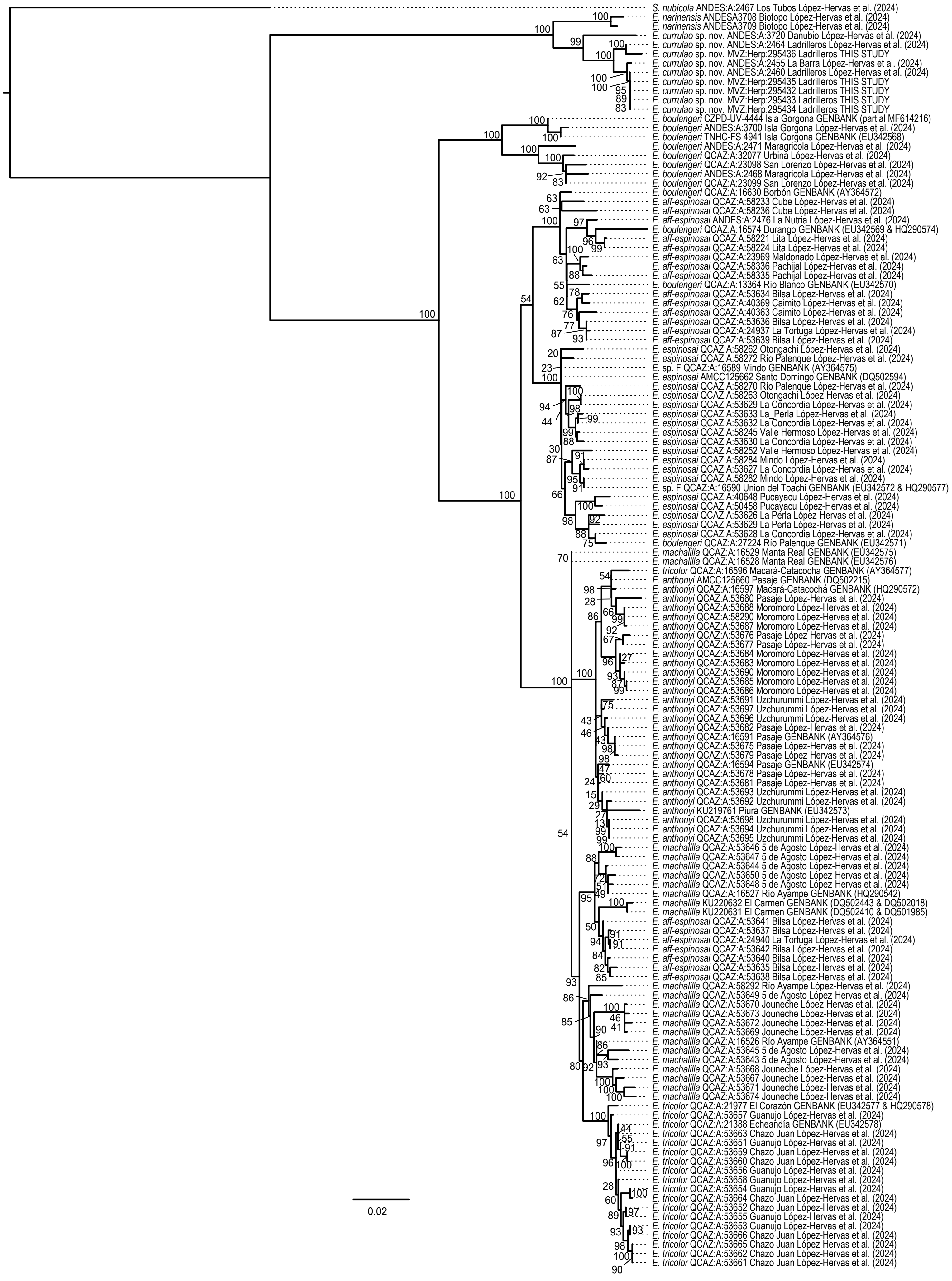
